# Cross regional coordination of neural activity in the human brain during autobiographical self-referential processing

**DOI:** 10.1101/2023.06.26.546582

**Authors:** James R. Stieger, Pedro Pinheiro-Chagas, Ying Fang, Zoe Lusk, Claire M. Perry, Anthony Wagner, Diego Contreras, Qi Chen, John Hugenard, Vivek Buch, Josef Parvizi

**Affiliations:** Laboratory of Behavioral and Cognitive Neuroscience, Human Intracranial Cognitive Electrophysiology Program, Stanford University School of Medicine, Stanford, USA; Department of Neurology and Neurological Sciences, Stanford School of Medicine, Stanford, USA; School of Psychology, South China Normal University, Guangzhou – China; Department of Psychology, Stanford University, Stanford, USA; Department of Neuroscience, University of Pennsylvania School of Medicine, Philadelphia, United States of America; Department of Neurosurgery, Stanford University, Stanford School of Medicine, Stanford, USA

## Abstract

For the human mind to operate, populations of neurons across remote regions of the brain need to coordinate their activity in the subsecond temporal scale. To date, our knowledge of such fast interactions involving cortical and subcortical structures in large brains, such as the human brain, remains limited. Here, we used stereo-electroencephalography (sEEG) recordings across four brain regions that are known, from decades of work, to be important for autobiographical memory processing. Our recordings involved 31 human participants implanted with intracranial electrodes in the hippocampus (HPC), posteromedial cortex (PMC), and ventromedial, as well as orbital subregions of the prefrontal cortex (OFC). In 14 subjects, we also recorded simultaneously in the anterior thalamus (ANT) across various experimental conditions and with direct electrical stimulations. Our observations provide new lines of correlative and causal evidence about the spatiotemporal profile of oscillatory coordination of cortical and subcortical activity during self-referential memory-based processing.

## INTRODUCTION

Past studies have reliably and consistently highlighted the importance of specific brain regions in memory operations ^1^. In the current study, we tapped into this large body of evidence and aimed to understand the fast (i.e., subsecond) spatiotemporal profile of activity within and across some of the brain regions that are known, from decades of work, to be co-activated during self-referential processing involving autobiographical memory, which requires the integration of a complex set of cognitive functions including not only episodic retrieval, but also, self-reflection, emotion, and semantic processes^2–8^.

Prior studies have shown that the hippocampus (HPC) and cortical structures beyond the medial temporal lobes (MTL), especially the posteromedial cortex (PMC, constituting retrosplenial and posterior cingulate regions) and the ventromedial prefrontal cortex ^3,4,9–15^ – along with some other regions of the brain - are co-activated during self-referential autobiographical memory processing. However, neuroimaging signals from a large mantle of the prefrontal cortex in its orbital surface are impacted by signal dropout^16^, constraining progress in understanding its functional role in the same cognitive processes while lesion studies have implicated the importance of orbital prefrontal cortex in such functions^17^.

While the majority of neuroimaging studies in human cognitive neuroscience has remained corticocentric^18^, recent work is beginning to elucidate the importance of subcortical structures, such as the anterior thalamus (ANT) for autobiographical memory processing^19,20^. Prior studies in the human brain, during rest or sleep, have shown that stimulations of the ANT, affects the hippocampal as well as cortical gamma activity^21^; thalamic spindles facilitate cortico-cortical and hippocampo-cortical co-rippling^22^; thalamic spindles are generated during sleep between cortical down and up-states^23,24^; and lastly, sleep spindles precede their neocortical counterparts and were initiated during early phases of thalamic slow oscillations (∼1Hz)^25^. Moreover, using direct recording from the ANT coupled with scalp EEG in human subjects, it has been shown that higher memory for complex photographic scenes was associated with theta phase synchrony as well as coupling between phase of theta oscillations recorded on the scalp and the power of gamma activity recorded directly in the ANT during encoding of stimuli^26,27^.

Motivated by the prior studies, and inspired by the extant evidence that the cognitive functions of the brain crucially depend on inter-regional coupling and precise timing of the coordinated activity of neuronal populations across multiple brain regions^28–33^, we aimed to explore the mode of cross regional co-engagement in the human brain by focusing on four brain areas that are already known, from decades of work, to be engaged during cognitive conditions of memory-based self-referential processing such as autobiographical memory retrieval. We emphasize that the aim of the current study was to leverage the temporal resolution, and the simultaneity of recordings across ANT, HPC, PMC, and OFC to understand cross-regional interplay. The experimental tasks used here are similar to autobiographical experiments employed in neuroimaging studies^3^, but not designed to decipher the precise cognitive mechanisms of memory retrieval per se or the importance of each specific brain region during each specific stage of a complex cognitive function such as autobiographical remembering. Future studies, with specific task designs are needed to explore the intricate regional specific mechanisms of autobiographical retrieval.

## RESULTS

### Demographics

In the following text, we only summarize the prominent findings of our analyses, while detailed statistics are provided in Supplementary **Tables S1-S12**. **Table S1** provides more detailed demographic and clinical data, our cohort consisted of 31 subjects with focal refractory epilepsy, 14 of whom had simultaneous recordings in the ANT. The rationale for multi-site thalamic recordings are detailed in our recent publication^34^. Twenty-six participants had implantation across at least 3 regions of interest (ROIs, i.e., ANT, HPC, PMC, or OFC) yielding sufficient data for *within-subject and within-hemisphere* analyses. All patients were undergoing invasive intracranial monitoring with stereo-encephalography (sEEG) electrodes (*AdTech In*c) as part of routine clinical evaluations. While clinicians probed the source of seizure activity in these patients’ brains, we invited them to participate in experiments in which they judged the accuracy of autobiographical episodic statements by retrieving memories of their past events (**Fig 1a**, detailed in Methods), or rested quietly in their bed while electrical stimulation procedures were applied. The research protocol was approved by Stanford University Institutional Review Board and all subjects provided informed consent.

**Fig 1:**
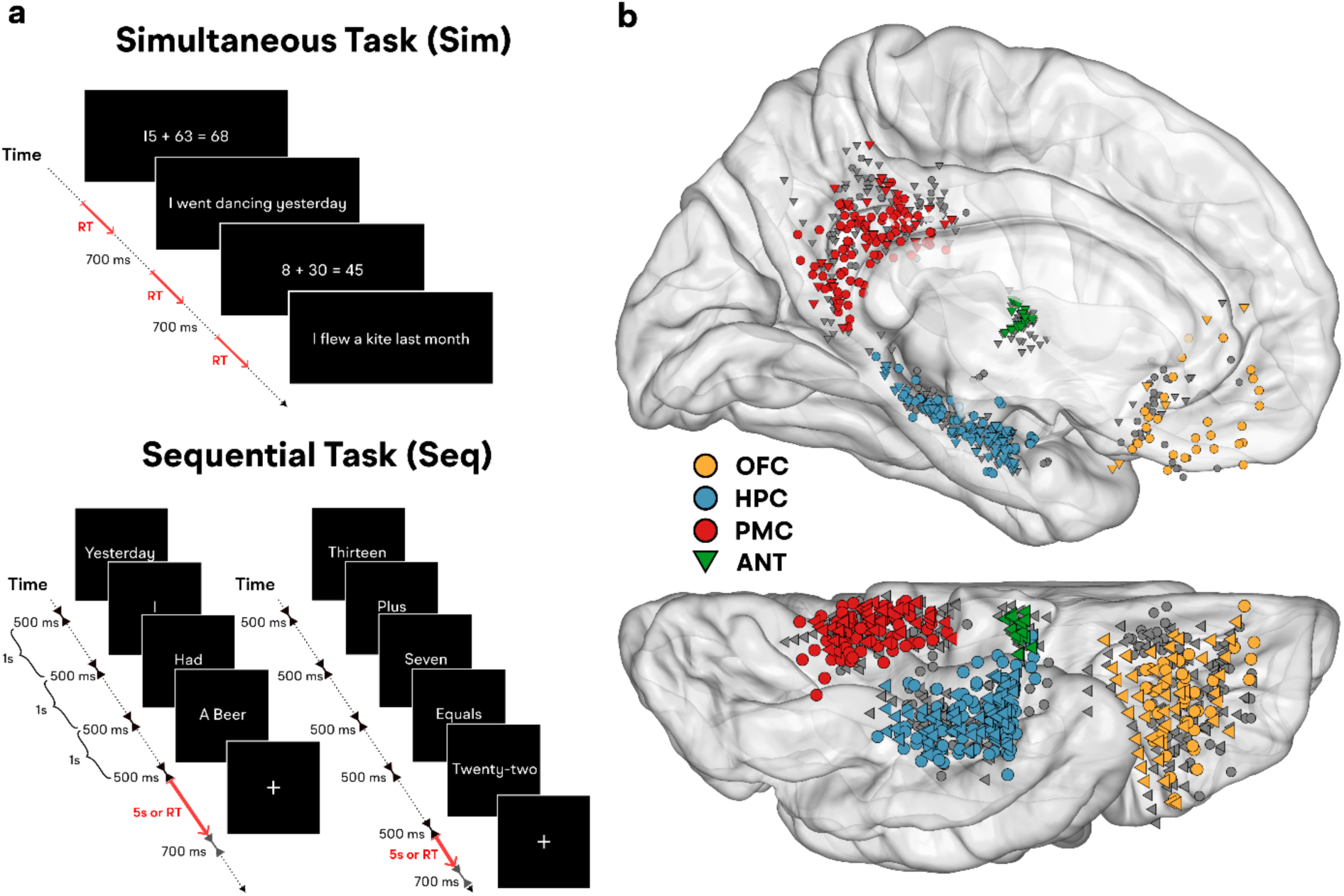
Experimental task design and recording sites: **a)** Task Design. Participants completed two tasks in which they decided whether a statement presented on the screen was true or false. The statements appeared in their entirety during the Simultaneous (Sim) Task (**a**-upper panel) or presented one word at a time during the Sequential (Seq) Task (**a**-lower panel). Note that a complete AM probe and non-AM probe in the Sequential Task unfolded across 4 and 5 screens, respectively. Non-AM conditions in the Simultaneous Task were either Math (in digit form only) or non-AM semantic statements (FACTS). Non-AM trials in the Sequential Task contained Math statements appearing in both number-word (presented) and digit form. **b)** Electrode locations. We recruited 31 participants with 812 electrodes across 4 ROIs —Orbitofrontal Cortex (OFC, n = 298), Hippocampus (HPC, n = 248), Posteromedial Cortex (PMC, n = 215), and Anterior Thalamic Nuclei (ANT, n = 51; note: ANT recorded in a subset of 14 patients; covered sites in these participants are identified with triangles). Colored markers represent sites that displayed significantly increased high frequency (HF) power above baseline and during AM compared to non-AM trials in at least one of the tasks. OFC electrodes were located either in the vmPFC or orbital region of the PFC. All vmPFC electrodes were below the callosal level in the individual brain, but when projected from native space to standard space, some of these electrodes may appear to fall dorsal to this level. Relatedly, for visualization purposes, the HPC electrodes are projected to the surface, and some may appear out of the HPC in the standard space. All HPC electrodes were confirmed to be within HPC in the native brain space.

### Electrode Coverage

We aggregated data from 812 recording sites across the left (N=447) and right (N=365) hemispheres and 4 ROIs: OFC [(total: per subject average ± standard deviation) 298: 9.61±6.47], HPC [248: 8.00±3.94], PMC [215: 6.94±5.73]. In these patients, 14 had recordings in the ANT [51: 1.65±2.04].

### Behavioral Data

Participants completed 268±123 (AVG±SD) trials in the Simultaneous Presentation (Sim) Task: 97 AM and 97 non-AM (**Fig 1a**) as well as Fact and Rest trials (i.e., a cross hair appearing at the center of the screen when subjects were instructed to rest). These trials were excluded from the analysis (except in **Fig 3c** where we show responses to Facts). In the Sequential Presentation (Seq) Task 220±81 trials were completed (110 AM, 110 non-AM; **Fig 1a**). The AM trials asked patients to judge the accuracy of common past experiences enabling their use with all participants (e.g., “Today I saw a doctor” or “Yesterday I took a shower”). The correct answer for the doctor and shower statement should be “Yes” and “No”, respectively because of the participant’s surgery. We found that the subjects’ responses were highly accurate for verifiable trials (Sim Task: subjects’ accuracy = 91%±8%; Seq Task = subjects’ accuracy= 83%±7%). Likewise, the high level of participants’ task engagement and the accuracy of their responses were documented by their high response accuracy during the non-AM (math) trials (Simultaneous Task: 92.7%±6.1%; Simultaneous Task: 89.5%±7.8%).

**Fig 2:**
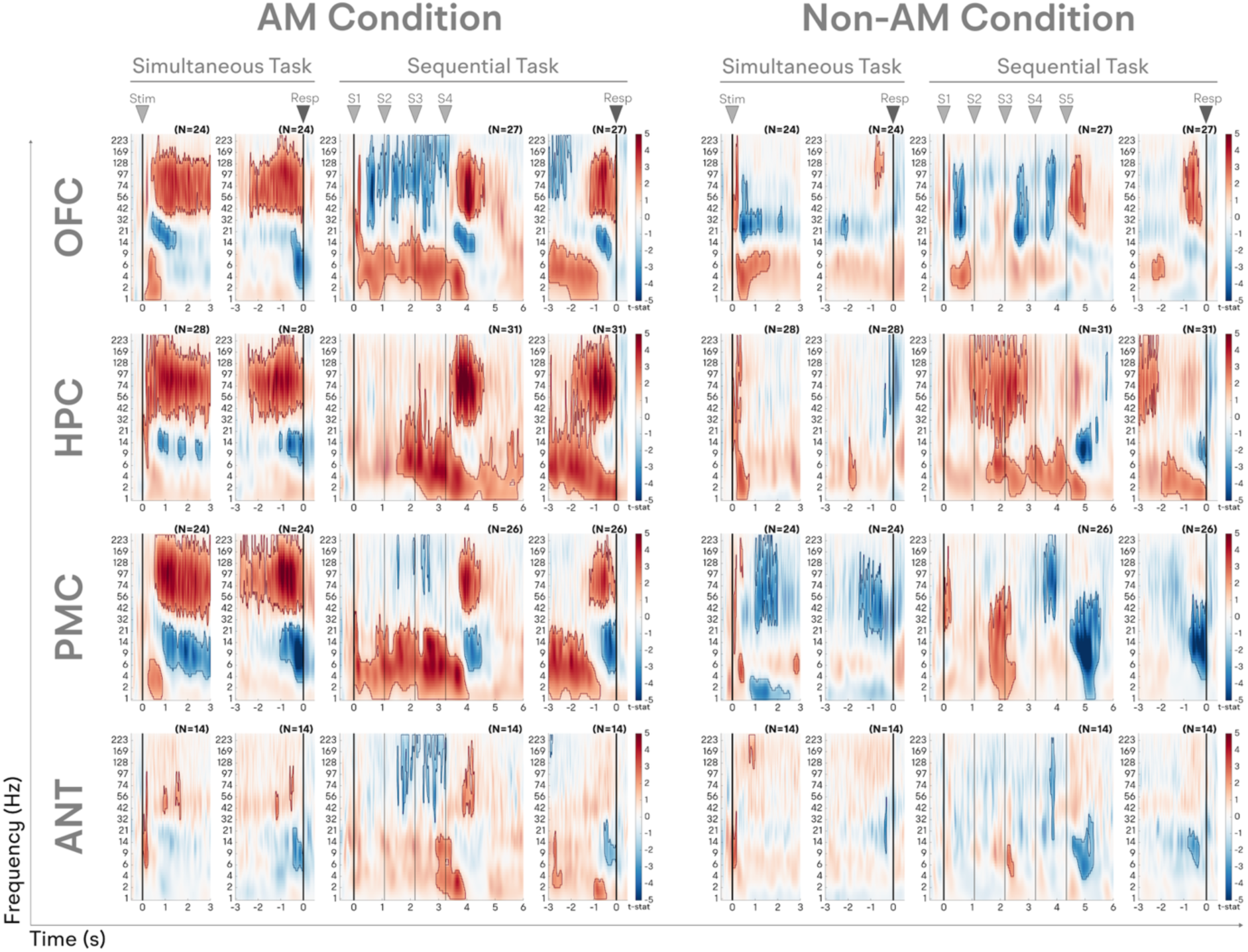
Similarities and differences in the profile of activity across the four regions of interest. Each panel displays the group-level stimulus (Stim or S1-5) and response (Resp) locked changes in power from baseline (prior to the presentation of the first stimulus) during the Simultaneous and Sequential Tasks. Note that the number of AM and non-AM stimuli in the Simultaneous Task were 4 and 5, respectively (see Fig 1). (Left) Autobiographical memory processing (AM Condition) (Right) Arithmetic calculation (Non-AM/Control Condition). Vertical black lines in the sequential stim-locked panels show the time each stimulus was displayed (S1-5). The color represents the statistical reliability of increases (Red) or decreases (Blue) in power from baseline computed across patients with coverage in a given region (t-value against the null hypothesis of no change in power from baseline; number of participants is presented above each panel). Significant deviations in power from baseline were identified with cluster-based permutation tests (CBPTs). Significant time-frequency clusters are highlighted and outlined in red and blue (all p < 0.0125, Bonferroni corrected for 4 ROIs).

**Fig 3:**
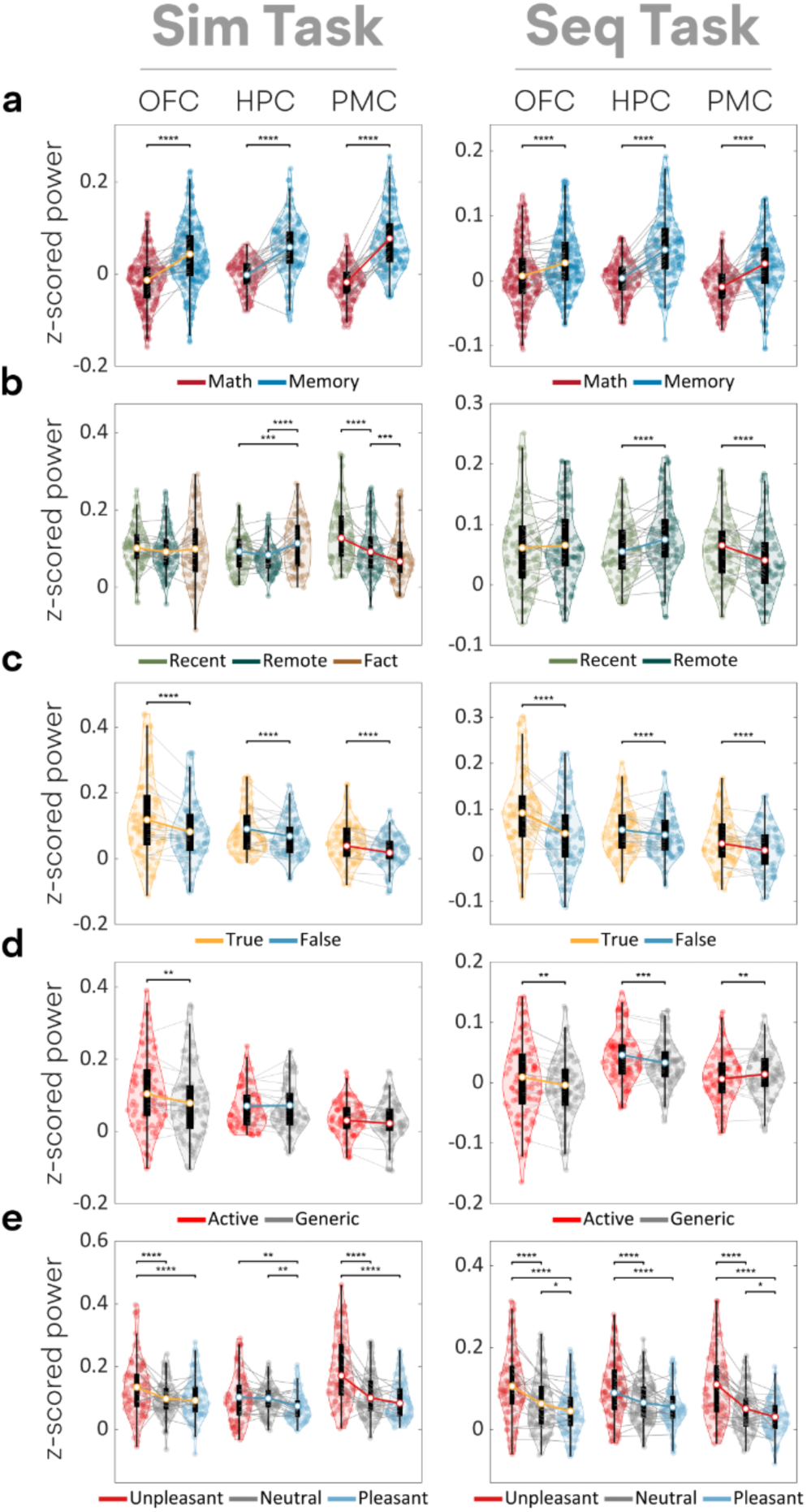
HF Power (70-170Hz) in each ROI is modulated by the memory content. Dissociable interactions between anatomical location and **a)** task condition; **b)** memory age; **c)** patient response; **d)** verb structure; and **e)** memory valence. Task content (**a**) was calculated across all electrodes in a given patient. Data for the other panels (**b-e**) were only calculated within the sites with significant memory related activations (colored sites in Fig 1b). In the violin plots, each colored dot represents one electrode, black boxes display the interquartile range, grey lines show individual subject averages, and colored lines show the average across subjects. (* p<=0.05, ** p<= .01, *** p<=0.001, **** p<0.0001-corrected for multiple comparisons). (See **Table S3-5** for full Statistics). See Methods for definition of each memory content type. See the main text and **Fig S1c** for data from the ANT.

These findings confirmed that the participants understood the task requirements and remained engaged throughout the experiments (more details in **Table S2**).

### Proportion of recording sites displaying significant responses during AM processing

We first identified the recording sites that displayed higher activation - as indexed by the power of *high-frequency activity* (HFA) during autobiographical memory (AM) trials compared to both baseline (i.e., a subset of the 700 ms inter-trial interval) and the non-AM (Math) condition. We emphasize that the rationale for choosing both baseline *and* the non-AM condition was not to claim selectivity but to exclude responses linked to motor (e.g., clicking mouse button) or to generic attention or visual processing. In the Simultaneous Task, subjects also evaluated semantic statements labeled as Facts (e.g., “Paris is in Europe”). These trials served as another non-AM control condition. Our analysis identified different proportions of sites per ROI that were engaged significantly higher during AM condition, with the highest proportion noted in the HPC (56%±33%) followed by PMC (54%±34%), OFC (37%±28%), and ANT (21%±37%) (colored sites in **Fig 1b**).

These findings demonstrate that in a given ROI, not all, but only select populations of neurons, are engaged during the studied cognitive process.

### Similarities and differences in power spectral patterns across regions

We included recorded signals from *every available site within every ROI* to assess the similarities and differences in power spectral patterns across all four ROIs during the AM condition (**Fig 2** – “AM-Condition”). In the Simultaneous Task, AM-related sites in HPC, PMC, and OFC responded with an initial and transient significant increase in low frequency power (LF; 1-6Hz) followed by a decrease in mid-range frequencies (∼8-30Hz, overlapping with the traditional alpha and beta ranges) and increase in high gamma (70-170Hz) power – akin to many similar findings during other cognitive tasks ^35–37^. Of note, 62% of ANT sites displayed greater power in gamma (32-58Hz) than high-gamma, range (T(50)=2.24,p=0.029). For the sake of simplicity, we will refer to high gamma activity in HPC, PMC, and OFC and gamma activity in the ANT as “*high frequency*, HF” activity. In the AM condition of the Simultaneous Task, when the presentation of each stimulus was timed and sequential, a significant increase in the HF range occurred in all ROIs *after* the last stimulus (Mixed-Effects Models (MxM): F(561)=11.47, p<0.0001). By comparison, LF activity in all ROIs was locked to the presentation of each stimulus (**Fig 2**). Mixed effects models (MxM) confirmed that this effect was stronger during AM than non-AM trials (**Table S3, S4;** LF memory effect (MxM): F(567)=7.39, p=0.0007, HF memory effect (MxM): F(561)=11.47, p<0.0001 – unless otherwise noted, claims made in the text apply to both tasks, but stats are reported only for the Simultaneous Task to reduce repetition).

These findings demonstrate that the average of recorded signals for every ROI (including all sites without any selection bias) displayed a similar profile of power spectral pattern in the AM condition.

### Modulation of high frequency activity by memory content

It is well known in the field of autobiographical memory research that the participants (when making judgments about autobiographical statements) are usually engaged in a myriad of cognitive processes such as episodic retrieval, self-reflection, emotion, and semantic reasoning^38^. However, such cognitive processes (imagination and reasoning) are expected to occur more during correct rejection statements than during correct hit statements. To examine if the recorded HF activity tracks certain aspects of memory processing, rather than generic decision making or response preparation processes, we measured the effect of rejection vs hit and the content of memory itself on the power of induced neural activity during AM trials in the AM-engaged electrodes of **Fig 1b**. To avoid a bias in selecting a specific temporal window for power analysis, we chose stim-locked (500-1500ms following the retrieval cue) as well as response-locked (1s prior to response) windows and compared the HF power as a function of memory content. These comparisons represent changes in HF power from the pre-trial baseline of 500-200ms before the first stimulus. We analyzed HF power across different conditions of memory age (“remote” such as “Last month” vs. “recent” such as “Today” etc.), subject’s response (responded as “True” i.e., correct hits vs. responded as “False” i.e., correct rejections), memory valence (“Pleasant” such as “Today, I had visitors” vs. “Unpleasant” such as “Last week I had a seizure” etc.), and verb structure (“active verbs” such as *kicked* or *drank* vs. “generic” such as *had* or *was* etc.) (**Fig 3** and **Table S4** for statistical details).

Across all four regions, correct hits elicited significantly more HF activity than correct rejections and trials with negative valence (patient response-MxM: F(286)=5.67,p=0.004, true-false: T(299)>4.49,p<0.0001). Moreover, as shown in **Fig 3**, a significant difference in the power of HF activity during AM compared to non-AM (semantic fact condition) is noteworthy. Additionally a significant effect was seen for pleasant vs unpleasant statements (memory valence-MxM: F(595)=8.48,p<0.0001, unpleasant-neutral: T(596)>5.32,p<0.0001). However, we observed an interaction between memory content and HF power suggesting regional HF activity was not homogenous across all ROIs. For instance, consistent with a recent study of hippocampal sharp ripples^39^, the HPC recording sites showed stronger HF activity during the processing of remote versus recent memories (memory age-MxM: F(295)=24.25,p<0.0001, HPC recent-remote: T(302)=-5.28,p<0.0001) but the opposite was the case in the PMC (PMC recent-remote: T(302)=+4.52,p<0.0001). As for ANT sites, a similar analysis revealed higher HF power for AM trials, “True” trials, and unpleasant trials (**Fig S1c**, **Table S5;** trial type-MxM: F(82)=22.6,p<0.0001, AM-Non-AM: T(80)=4.75,p<0.0001; patient response-MxM: F(30)=10.44,p=0.003, true-false: T(30)=3.23,p=0.003; memory valence-MxM: F(60)=17.94,p<0.0001, unpleasant-neutral: T(60)=5.45,p<0.0001).

These findings demonstrate that the recorded electrophysiological responses tracked the behavior of the participants. The consistency of these effects observed across both tasks suggests the following: 1) findings can be attributable to the AM process rather than a confound of task design, and 2) each ROI’s role in AM processing may at least be partially distinct since the modulation of activity by memory content is not homogenous and identical across sites.

### Cross-regional phase coherence following the retrieval cue

We confirmed the presence of oscillations in the power spectrum in the form of peaks in the LF range above the aperiodic background utilizing a method as described elsewhere^40^. Additional posthoc tests verified these results were not driven by volume conduction or choice of common average referencing (see supplemental material). To explore how oscillations across ROIs are coordinated and synchronized, we relied on the measure of *phase coherence,* which reflects the degree to which the phase of signals in two brain regions – that are themselves formed as a result of the summation of postsynaptic potentials generated in a large number of neurons^29^ – are coherent (**Fig S2**). For this, we measured 1) regional phase coherence across all trials in a given ROI (labeled as intertrial-phase coherence, ITPC; **Fig S3**), 2) phase coherence in a given ROI across the two hemispheres (labeled as inter-hemispheric within-ROI phase coherence, ISPC_within_; **Fig S4A**), and 3) phase coherence between two different ROIs (labeled as between ROI inter-site phase coherence, ISPC_between_; **Fig S4B**).

A significant ITPC was found in all ROIs in the LF range following the appearance of each stimulus. This effect was observed during both AM and non-AM trials in both Simultaneous and Simultaneous Tasks (Cluster based permutation tests (CBPT); p<0.003) (**Fig S3**-Left). The contrast between ITPC findings and ISPC_within_ and ISPC_between_ findings was made clear with data obtained during Sequential Task: Unlike significant increases in ITPC after each single stimulus, we observed increased ISPC *only* after the final stimulus was presented. As noted, this last stimulus was the associative memory cue that enabled the participant to be engaged in making memory-based decisions in AM and Math conditions (**Fig S3** -Right, CBPT; p<0.002; **Figs S4A,B**). Of note, in the non-AM (math) condition, the increse in cross-regional coherence did not occur when the subjects were adding numbers together, but rather when the number on the screen was being compared to the calculated value stored in memory. The ISPC_between_ appeared after the last stimulus in both experiments. Memory vs Math comparisons shown in supplementary **Figs S3-S5** and for ANT contacts in **Figs S1d and S5**.

The above findings demonstrate that consistent phase relations *within* ROIs occur non-selectively after the presentation of each stimulus regardless of its content while the phase relations *between* sites is selective to the time when participants were engaged in retrieving memory-based information (both AM and non-AM condition). We are confident that the ITPC findings were not simply due to visual evoked potentials (VEPs) since the plotted raw EEG signals during both AM and non-AM condition did not show similar time-locked changes after each visual stimulus (**Fig S6**). However, the ITPC and even ISPC findings could be related to the phenomenon of “*cognitive”* event related potentials (ERPs) that have been hypothesized to be caused by stimulus-induced increases in the phase-locking of ongoing EEG activity ^41,42^ but not visually-evoked changes in the EEG power alone ^43^.

### Timing of hippocampal LF power and cross-regional phase coherence

By leveraging the high temporal resolution of our approach, we used a measure of response onset latency (ROL, detailed in Methods and **Fig S7**) to explore how changes in the power of LF or HF activity and the coupling of phase between remote sites (ISPC_between_) relate to each other across time. We specifically examined whether these features occur simultaneously or progress in stages while the participants engaged in autobiographical processing (**Fig 4**). To compare the timing of events across sites, we only used data across pairs of sites *within the same hemisphere and within the same subject*. This analysis revealed a specific *temporal order* in that the rise of LF power, in all ROIs, was followed by coupling between sites (ISPC_between_; hereafter referred to as ISPC) and then the rise of HF power. Interestingly, the same statistical pattern was seen during AM condition in both experimental tasks (Mixed-effects model (MxM): F(1026)=12.73,p<0.0001; see **Table S6-8** for detailed stats).

**Fig 4:**
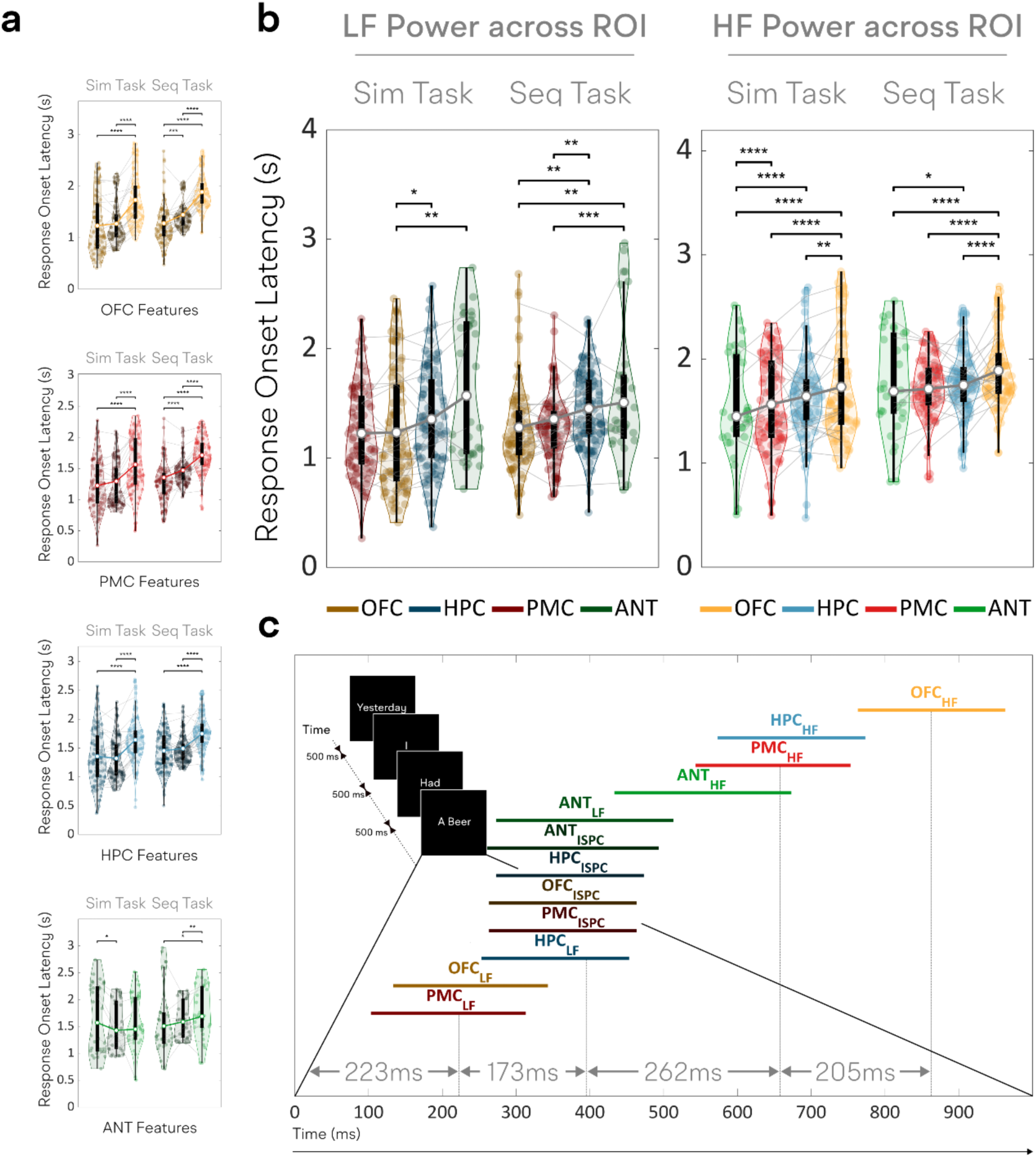
Temporal profiles of electrophysiological activity during autobiographical memory processing. **a)** ROL estimates for individual electrodes in each ROI for LF power, intersite phase coherence, and HF power (in that order) during Simultaneous and Sequential tasks. An interesting temporal order was seen when modeling response onset latency (ROL; p<0.0001). **b)** Response onset latency of LF (Left) and HF (Right) power compared across ROIs. Violin plots: Each colored dot represents the ROL of one feature within a given electrode, black boxes display the interquartile range, grey lines show individual subject averages, and colored lines show the average across subjects. (**** p<=0.0001). **c)** The lower panel shows the 95% confidence interval for the estimates of each ROI feature following the final stimulus in the Seq Task. The vertical lines represent significant differences between groups of features. Note: Results come from sites with significant AM related activations identified in Fig 1b (all ANT electrodes were included and reflect the ROL of gamma power).

The timing of LF activity in the HPC and ANT was more noteworthy in that it persisted longer and was present when the LF activity in the OFC and PMC had already subsided (**Fig 4b**-left, MxM: T(964)> 2.77,p<0.029; see **Tables S6-7**). However, the nature of the LF power differed between HPC and ANT. In HPC, the probability density function (PDF) of LF activity peaked after the recall cue and persisted along with the ISPC PDF, while the probability of ANT LF activity (like OFC and PMC) dropped sharply before the peak of the ISPC PDF but displayed a higher probability near the end of the trial.

In terms of the timing of HF activity, the ANT HF power appeared first while the OFC HF power appeared last (Early ANT HF – MxM: ANT-HPC/OFC: T(1029)<-2.63,p<0.04; ANT-PMC was only significant in the Sim Task: T(963)=4.93,p<0.0001; Late OFC HF – MxM: OFC-Others: T(1034)>5.65,p<0.0001). The time of HF activity in HPC and PMC did not differ (**Fig 4b**-right, MxM: T(1031)=0.86,p=0.83). As seen in **Fig 4c** these data highlight three important findings: First, the onset of HPC LF activity is contemporaneous with the global LF synchronization (ISPC); second, the HF activity in ANT is the first to erupt before the HF activity in other regions is triggered; and third, OFC is the last ROI in this HF cascade.

To validate the observations made with the ROL measure, we performed additional analysis of lagged correlation between signals by pooling data from 13 subjects who had optimal coverage across ANT, HPC and other ROIs. This provided further evidence to support our ROL findings (**Figs S8-S9**). First, HPC LF phase was seen to be leading the phases of both OFC (T(75)=2.96,p=0.004) and ANT (T(154)=3.45,p=0.0007). The same trend was observed for PMC (T(222)=1.06,p=0.29), but this was not significant, perhaps suggesting a tighter relationship between HPC and PMC. This supports our findings of HPC LF activity driving the coherence events observed in other ROIs (**Fig S8**). Finally, OFC HF activity was seen to significantly lag the HF activity of all other ROIs, with this effect being most significant for ANT (HPC: T(123)=5.95,p<0.0001, PMC: T(150)=4.98, p<0.0001,and ANT: T(153)=9.60, p<0.0001), which supports our findings of delayed response onset latency of OFC HF power compared to other ROIs (**Fig S9**). The above findings demonstrate a cascade of electrophysiological events within and across ROIs linking HPC LF activity with inter-site phase coherence before regional HF activity is induced.

#### Probing cross-regional interplay with causal measure

To provide causal information, we applied electrical stimulation using the well-known method of Single Pulse Electrical Stimulation (SPES; referred to as STIM for ease of understanding). In this procedure, repeated single pulses were delivered, during the resting state, between adjacent pairs of implanted electrodes while recording from all other implanted electrodes in an individual subject’s brain^44^. The presence and timing of evoked responses in a recorded region suggests that the stimulated seed region is physiologically connected with, and has the means to exert an effect upon, the target site^45–47^.

Using the data from the STIM approach, we collected descriptive data regarding the extent of causal effective connectivity across the four ROIs. We found a large proportion of stimulated sites within the HPC causing significant time-locked evoked responses in ANT (74.2%±6.5%), PMC (51.2%±10.2%) and OFC (36.2% ±7.1%). However, the proportion of sites in each of the other ROIs whose stimulation generated evoked responses in the HPC was smaller (details in **Fig 5a** and **Table S9).** ANT, on the other hand, appeared to have bi-directional effective connectivity with OFC (ANT→OFC: 56.8% ±10.2%; OFC→ANT: 58.3% ±12.2%) and PMC (ANT→PMC: 41.3% ±10.6%; PMC→ANT: 33.5% ±6.2%) suggesting ANT is strongly influenced by incoming signals from the HPC while it bi-directionally communicates with the OFC and PMC.

**Fig 5:**
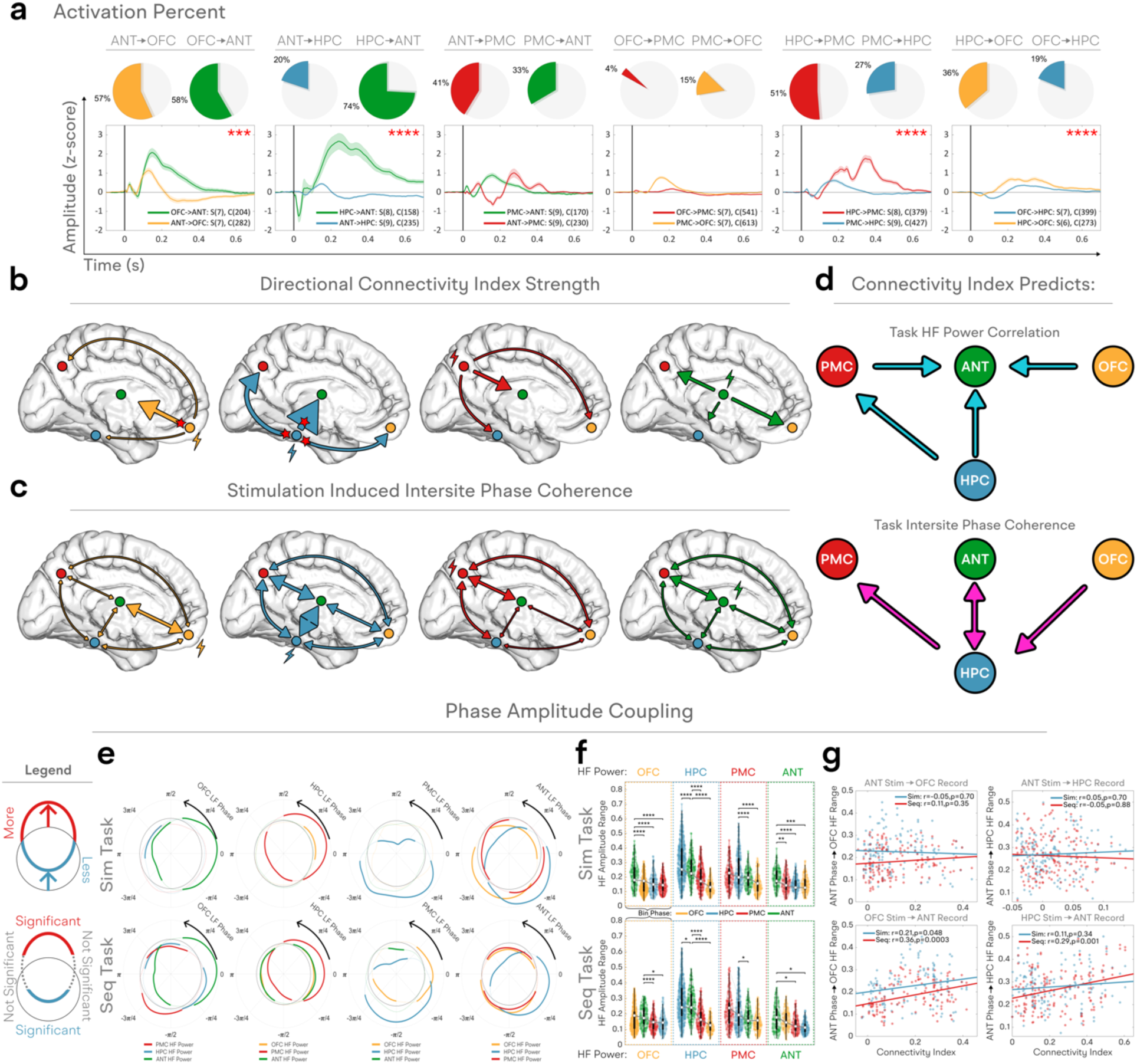
Profile of effective connectivity across ROIs reveals directionally asymmetric axis of influence between ROIs. **a)** Causal effective connectivity with single pulse electrical stimulation. (Upper) Percentage of stimulation sites causing significant voltage deflections in recording sites within distant ROIs (X→Y: stimulate X, record in Y). (Lower) Each plot shows the average voltage deflection response of a given region to single pulses of electrical stimulation in another ROI (S: number of subjects, C: number of connection pairs). Red stars indicate significant differences in the directionality of influence between ROIs as measured by connectivity index. **b**) A connectivity index based on intertrial phase coherence (ITPC) was derived as a measure of how consistently single pulses of electricity in one ROI affect the voltage in another ROI (Arrows are drawn in proportion to the strength of the connectivity index; Method: **Fig S11,** Statistics **Table S10**). The influence of HPC on other ROIs was significantly greater than the influence of other ROIs on HPC (all p<=0.0001). ANT displayed a strong influence on both OFC and PMC (greater than HPC, p<0.0001), and the OFC→ANT connection was significantly stronger than in the reverse direction (p=0.0001; no directional influence observed between ANT and PMC, p=1.0). **c**) Stimulation induced intersite phase coherence (ISPC) (Arrows are drawn in proportion to the strength of induced coherence between ROIs; Statistics: **Table S11**). Note: 1) Stimulation of a given ROI causes the greatest increase in ISPC between that ROI and ANT, suggesting when an ROI is stimulated, ANT is likely to synchronize its activity with this ROI. 2) HPC stimulation causes the greatest increase in coherence across all electrode pairs suggesting that when HPC is stimulated, in general, all cross regional pairs will become more phase coherent. Note that the stimulation induced artifacts were removed before the calculation of ISPC. **d**) Here we combine data from two modalities: experimental AM-condition and STIM. Arrows are drawn when the X→Y connectivity index significantly predicted HF-HF correlations (above) or ISPCs (below) in BOTH tasks (See **Figs S12**-**S13** for the strength of correlations across all pairs). The upper cartoon summarizes only statistically significant relationships between HF-HF correlations during task and strength of causal effective connectivity (i.e., connectivity index) induced electrically during rest. The data presented in this cartoon should be interpreted as follows: a given HPC neuronal population had highly correlated HF activity during AM retrieval with another neuronal population in the PMC if the stimulation of that HPC population showed strong and reliable evoked responses in the PMC population (but not the other direction); a given site had correlated HF activity during AM retrieval with ANT only if the site had strong causal connectivity to the ANT but not the other way around; or if the HF activity of an ANT and OFC site was similar during AM processing, it was more likely that the OFC site would evoke a reliable response in ANT when electrically stimulated. This noteworthy because the HF correlation during AM processing is a symmetrical statistical relationship that was predicted by the direction of causal effective connectivity. In the lower cartoon, we use the ISPC values instead of HF-HF correlation values. Here, the causal connectivity of a site to the HPC predicts the degree of that site’s phase coherence with the HPC. **e**) Significant cross regional phase amplitude coupling (PAC) between LF phase (1-6Hz) and HF power was observed across all ROIs and both tasks. In these plots, the thin black circle at the center represents the expected uniform distribution of HF amplitude if there were no relationship between LF phase and HF amplitude (which can be thought of as a chance level). If one ROI’s HF amplitude is greater than the expected uniform distribution at a specific LF phase of another ROI, this line will extend outside the inner circle. Similarly, if one ROI’s HF amplitude is less than the expected uniform distribution at a specific LF phase of another ROI, this line will be found inside the inner circle. When certain phases were found to be consistently related to greater or less HF amplitude in another ROI (compared to a permutation distribution where the HF amplitude and LF phases of different trials were shuffled) the significant phases are drawn with a thick line. If there was no evidence of greater or lesser HF amplitude for a specific phase, this part of the circle is drawn with a dashed line. The phase specific effect appeared consistency across both tasks in that most ROIs show significant increases and decreases in HF power at the same phase of the reference signal across tasks. For these plots, HF power was averaged across 18 uniform bins spanning −π to +π of an ROIs’ LF phase. Thick lines represent phases where HF power is significantly increased (outside middle circle of expected uniform distribution), or decreased (inside middle circle of expected uniform distribution), compared to a permutation distribution where HF power was binned according to LF phase from a different randomly selected trial (CBPT, p<0.004). While the circle diagrams present evidence that there is a significant asymmetrical relationship between HF amplitude of one ROI and another ROI’s LF phase during the task, the reliability of these findings across subjects was measured with mixed effects models and is shown in panel **f**. Significant interactions of the strength of PAC were observed between ROIs (p<0.0001). The important thing to note for both the circle diagrams in panel **e** and panel **f** is that the same values of HF amplitude in one ROI show differential relationships to the LF phase of other ROIs. In particular the timing of OFC and HPC HF activity appeared to have a consistent relationship with ANT LF phase (p<0.05), while PMC displayed a closer relationship with HPC LF phase (p<0.05). **g**) the extent to which OFC and HPC HF activity locks to ANT LF phase was predicted by OFC’s and HPC’s ability to influence ANT through electrical stimulation (p<0.05), but not the other way around(p>0.3). This again supports our findings from panel **d** in that, for the same measure of statistical relationship during AM processing (ANT LF – OFC HF PAC), PAC could be predicted by inflow to ANT (OFC→ANT connectivity index) but not outflow from (ANT→OFC).

The descriptive data, presented above, only reflect the extent of connectivity (i.e., proportion of sites within an ROI influencing sites in another ROI). In a separate analysis, we quantified the robustness of connections between a pair of sites using the measure of “*Connectivity Index*” (**Methods**, and **Fig S11).** This measure is based on the prediction that if A → B connections are robust, then stimulations in A will reliably evoke non-jittered and coherent responses in B across trials of stimulation.

Across all pairs of stimulated-recorded sites we confirmed significantly greater measures of *Connectivity Index* across pairs of sites that happened to be co-activated during the AM condition (Recording site AM coactive-nochange – MxM: T(3897)=2.24,p=0.025; Stim site AM coactive-nochange – MxM: T(3888)=4.709,p<0.0001). However, the strength of connectivity was not the same in both directions between ROIs (MxM: F(3890)=25.65, p<0.0001) suggesting an asymmetric directionality of signal flow across different nodes of the network. For instance, the strength of connectivity from HPC to all other ROIs was significantly stronger than from other ROIs to HPC (MxM: T(3890)>8.77, p<=0.0001; red stars in **Fig 5a,b**– the size of arrows in **Fig 5b** scales with Connectivity Index), suggesting that the stimulation of the HPC changes the activity of other ROIs but the stimulation of other regions does not equally affect the HPC. By contrast, ANT appears to receive the greatest proportion of evoked responses following the stimulation of any other ROI (**Fig 5a**). The asymmetry of inflow to ANT is not only significant for HPC, but also OFC (OFC→ANT > ANT→OFC MxM: T(3891)=4.92, p=0.0001). Importantly, we validated these findings by showing the thalamic recording sites closer to the ANT proper were more likely to display evoked responses following OFC stimulation (OFC→ANT vs. distance: r(24)=-0.46,p=0.041), and vice versa (ANT→OFC vs. distance: r(24)=-0.52,p=0.027; see **Fig S1f**).

Remarkably, the strength of connectivity from an ROI to ANT correlated with the degree of their co-engagement (each site’s power of HF activity) during the AM-condition (OFC→ANT r(100)=0.29,p=0.01, HPC→ANT r(106)=0.33,p=0.003, PMC→ANT r(64)=0.33,p=0.015; **Fig 5d** and supplemental figure **S12**). This is particularly noteworthy because it demonstrates that the strength of co-engagement of two sites during AM processing (HF power correlation; itself a symmetrical statistical measure) displayed a significant *directional* relationship with an ROI’s ability to *causally* influence ANT in one, but not the other, direction. This highlights the profile of ANT’s as a receiver (possibly for the purpose of integrating information). In other words, the strength of co-engagement between ANT and a contact X during memory processing is related to the site X’s ability to evoke responses in ANT when stimulated, but not ANT’s ability to evoke activity in site X. We emphasize that in **Fig 5d**, arrows are drawn if there was a significant correlation *across both tasks* between memory task engagement and an ROI’s ability to evoke reliable activity in another ROI measured by the Connectivity Index. By contrast, the strength of causal connectivity of HPC with other ROIs correlated consistently with the degree of their phase coherence during AM condition (ANT→HPC r(162)=0.37,p=0.0001, HPC→ANT r(106)=0.27,p=0.02, OFC→HPC r(313)=0.2,p=0.002, HPC→PMC r(282)=0.14,p=0.04; **Fig 5d** and **S13**). One exception was the HPC→PMC connectivity index that correlated also with the power of HF co-activation at the two sites (HPC→PMC r(282)=0.25,p=0.0002).

The measure of stimulation-induced intersite phase coherence (ISPC) also affirmed the generalizations mentioned above (**Fig 5c, Table S11**). In this analysis, we measured the increase in phase coherence *between pairs of sites* resulting from stimulating a given ROI. This is different than the measure of the Connectivity Index which relies on trial-by-trial coherence between responses within a single electrode as a result of stimulating another seed region (i.e., ITPC). Here, following stimulation of a seed region, we measured ISPC between all ROI pairs (after excluding contacts affected by the stimulation artifact). We found that the stimulation of the HPC, on average, produced the greatest observed increases in ISPC across *all* electrode pairs (HPC→(X-Y ISPC) > Z→(X-Y ISPC) MxM: T(1170)>7.13, p<0.0001) while the stimulation of each ROI caused the greatest ISPC between that particular ROI and ANT (X→(X-ANT ISPC) > X→(Z-Y ISPC) MxM: T(1168)>4.50, p<0.0001). This again highlights the importance of ANT’s role as a universal receiver of information in that, following stimulation of any ROI, ANT was most likely to synchronize its activity with the stimulated ROI. Once again, ISPC measures were stronger between sites that were co-engaged during AM condition (i.e., increased HF activity compared to baseline and non-AM condition) than sites that were not (Recording site AM coactive-nochange – MxM: T(1170)=3.611, p=0.0003; Stim site AM coactive-nochange – MxM: T(1170)=2.355, p=0.019). These results are shown with arrows proportional to the stimulation induced ISPC in **Fig 5c** (Further statistical details found in **Table S11**).

Finally, we calculated the speed of signal flow across ROIs by measuring the time to first prominent peak in sites displaying significant evoked responses (details **Table S12**). These results highlighted the centrality of the ANT in the studied AM-network. For instance, the speed of connectivity from an ROI to ANT was significantly faster than the speed of connections from the same ANT to the other ROI sites (T(44)=3.43, p=0.0026). Furthermore, connections from ANT to the other ROIs (on average) were significantly faster than the connections among the other ROIs (T(61)=2.62,p=0.01).

The above findings demonstrate that only a select proportion of populations of neurons are connected across different ROIs, and that the proportionality and strength of these connections are both asymmetric with more robust connections being present across pairs of populations that are co-engaged during the experimental condition. More notably, the outward connections of the HPC to all ROIs stand in contrast to inward and fast connection of all ROIs towards ANT.

#### Phase amplitude relationships

The measure of causal connectivity relies on the injection of electricity in one population of neurons while recording the evoked responses in other regions as a result of such artificial perturbations. To validate these findings with a relatively more naturalistic measure of connectivity we relied on the well-known measure of phase amplitude coupling (PAC) ^29,48^. In this analysis, we pooled data from only 13 subjects in whom optimal coverage was ensured across the HPC, ANT, and other two ROIs. This analysis revealed a significant relationship between HPC LF phase and the HF power of *all* ROIs (most pronounced for PMC). We found that, compared to a permutation distribution across channels, there were significant effects of PAC observed across all ROIs (cluster-based permutation tests (CBPT) p<0.004; **Fig 5e**). However, there appeared to be a differential relationship among ROIs (**Fig 5f**; MxM: F(3191)=38.84, p<0.0001). The strongest PAC observed was between HPC LF phase and HPC HF power [for the following X→Y PAC means binning Y’s HF power by the LF phase of X] confirming prior similar observations^30^.

Providing further evidence of the close link between HPC and PMC, we found the HPC→PMC PAC was significantly stronger than OFC→PMC (MxM: T(3250)=3.71,p=0.001) and ANT→PMC (MxM: T(3170)=2.97,p=0.016). However, this relationship was reversed with OFC. ANT→OFC PAC was significantly stronger than HPC→OFC (MxM: T(3156)=3.24,p=0.007) and PMC→OFC (MxM: T(3118)=4.59,p<0.0001), suggesting that OFC HF activity is more tightly linked to the phase of LF activity in ANT than the other ROIs. Furthermore, the OFC→ANT connectivity index significantly predicted how much OFC HF activity locked to ANT LF phase (r(129)=0.36,p=0.0003; **Fig 5g**). This again provides further support of our findings in **Fig 5d**, in that for the same measure of statistical relationship during AM processing (PAC between ANT LF phase and OFC HF power), PAC could be predicted by inflow to ANT (OFC→ANT connectivity index) but not outflow from (ANT→OFC); in these posthoc analyses we focused exclusively on the ANT/OFC/HPC relationship because of the prior statistically significant directional influence measured by the connectivity index. Interestingly, ANT HF activity did not preferentially lock to any other ROI’s LF phase (MxM: T(3190)<0.68,p>0.9). Taken together, these results demonstrate that the same HF amplitude during AM processing in one ROI shows asymmetric coupling to the LF phase in another ROI. Specifically, the HF activity of all ROIs was locked to HPC LF activity; OFC HF activity is most likely to pair with the ANT phase, and the HF activity in PMC is most likely to pair with HPC phase.

## DISCUSSION

In the current study, we simultaneous recordings across four brain regions gave us a unique opportunity to observe the profile of responses, as well as their timing, across the regions of interest. In the following text we first summarize the key observations before we focus on a discussion of those findings that are unique and relatively novel compared to extant literature, namely the temporal order of events, causal connections, and the results from the thalamus.

### Correlative Evidence

We analyzed signals across *all* electrodes present within the boundaries of each ROI and observed that only select populations of neurons were engaged during the experimental task (**Fig 1**, colored vs gray sites) – a finding that cannot be appreciated with neuroimaging tools based on group-based analyses. Next, we observed a remarkably similar “signature” of electrophysiological activity across all four regions in the subsecond space (**Fig 2**) – an observation that could have not been achieved if we had used a method with low temporal resolution. The profile of HF and LF activity in the Simultaneous Task was similar to the one obtained during the Sequential Task – i.e., in the time window after the presentation of the memory cue - hence offering replication of findings across two complimentary tasks and providing mechanistic information about the cognitive stage of at which these regions are co-engaged. Further, we confirmed that the stimulus-locked (500-1500ms following the retrieval cue) as well as response-locked (1000ms prior to response) power of HF activity was higher in hit statements compared to correct rejections and modulated by the cognitive content of memory statements including memory age (remote vs. recent), memory valence (pleasant vs. unpleasant), and verb structure (active vs. generic) – suggesting that the observed electrophysiological features were not independent from the cognitive content of the stimuli (**Fig 3**). Importantly, our findings revealed a few noteworthy differences in the signature of activity across regions during the stimuli preceding the presentation of the memory cue and across trial types with different memory content. We take these regionally specific idiosyncrasies as preliminary hints suggesting that the contribution of each region during different stages of memory retrieval is unique and can be decoded with goal-directed task designs. This hypothesis needs further exploration in the future. We used the above observations based on the data collected from *all sites* to identify the electrophysiological features differentiating the AM from non-AM condition. Once this was accomplished, we selected only the AM-activated sites across the ROIs to study how the presumed AM-related responses unfolded in time across the ROIs, and how the AM-activated sites across regions were coupled together. This analysis revealed a unique temporal order of events across frequencies (LF to HF) and across regions – a finding that was made possible because of simultaneous recordings with high temporal resolution and more importantly, high signal-to-noise ratio of observed physiological responses that enabled within-subject and trial-by-trial comparisons. When participants attempted to construct mental representations of cued personal past events, a cascade of temporally orchestrated electrophysiological events unfolded across the studied regions: regional LF activity → cross-regional phase coherence of LF activity → regional HF activity. (**Fig 4**).

### Causal Evidence

As temporal regularities do not necessarily imply causation^49^, it was important for us to replicate and validate the key findings from our correlative approach by the causal information obtained with direct causal perturbations of the sites of interest. As detailed in **Fig 5** and related supplementary material, our causal approach revealed several key findings: across a given pair of ROIs, many populations of neurons in one ROI can affect the other (A →B) while only a few populations of neurons may do the same in return (B → A); a majority of sites across a given pair of regions (e.g., OFC and ANT) may have strong causal effective connectivity, while only a minority of sites across another pair (e.g., OFC to PMC) may be influencing each other; causal connectivity between a pair of sites was strongest if the two sites showed strong co-activations during the experimental AM-condition; stimulation of the HPC, on average, produced the greatest observed increases in ISPC across all electrode pairs, and the strength of causal connectivity of HPC with other ROIs correlated consistently with the degree of their phase coherence during AM condition; by contrast, a site in ANT had strongly correlated co-activations with HPC or cortical sites during AM condition if those sites were able to evoke strong effects on the ANT when stimulated (but not the other way around).

The above causal measures were based on the measure of the strength of responses evoked by repeated electrical stimulations of a given seed region. Some might argue that the connectivity across regions during task and or rest may go beyond such artificial conditions. To explore the validity of our STIM-based findings, we measured phase amplitude coupling and lagged correlations which suggested that the same values of HF amplitude in one ROI had different relationships to the LF phase of other ROIs and highlighted the strong relationship between HPC– ANT and HPC–PMC as well as ANT–HPC and OFC–ANT.

### ANT and HPC in the Order of Electrophysiological Events

Our results resonate with the current scientific understanding of the memory system ^19^. Hippocampal LF activity was seen to lead a network-wide cross-regional phase coherence followed by thalamic “ignition” before a near-simultaneous HF co-activation of the HPC and PMC and, finally, a significantly delayed OFC HF activity. Our causal modulation approach validated the crucial role of the HPC in synchronizing the activity of other ROIs. Single perturbations of the HPC resulted in a network-wide cross-regional phase coherence. In addition, the causal effective connectivity between the HPC and other regions of the brain was clearly asymmetric and favored HPC→ other ROIs direction than the ROIs →HPC direction. By comparison the ANT, similar to other cortical regions, displayed trial specific HF modulation, robust ITPC following task stimuli, and consistent cross-regional ISPC following the presentation of the retrieval cue. However, unlike the HPC or other regions, ANT operated uniquely in that it was the first to increase HF power following the cross regional LF phase synchronization event – even before the rise of HF power in the HPC; ANT was the first to show an evoked peak following electrical stimulation of other ROIs [i.e., when a region X is stimulated, evoked responses are seen in X→ANT, ANT→Y, and then X→Y]; ANT was the most likely to synchronize with an electrically stimulated seed region with the strength of effective connectivity between a given ROI and the ANT predicting the degree of their co-engagement following the retrieval cue; and lastly, the phase of LF activity in ANT exhibited a robust coupling with the HF amplitude of all other ROIs’ which was predictable by the strength of OFC→ANT and HPC→ANT connectivity.

These findings highlight the important role of the HPC and ANT in enabling cross regional interplay in the memory network. Based on our results, one can view the HPC as a “universal synchronizer”, and by contrast, the ANT as a “universal receiver” within the memory network – a view that is compatible with the notion of ANT being a key player in memory based cognitive processes beyond serving as a simple relay station for the HPC output ^19^. In this model, the HPC can be seen as a mnemonic “hub” coordinating the engagement of other regions by phase resetting and creating cross-regional phase coherence to synchronize their co-engagement during retrieval. This is consistent with the evidence suggesting that the phase of LF activity (including hippocampal theta activity) reflects changes in the excitability of neural structures, and thus a key mechanism for timing the synchronization of neuronal assemblies that fire together^29,31,50–52 53,54^.

In the recent literature, ANT has been portrayed as an integrative hub and gatekeeper within the memory network^20^. The ANT’s central location within the brain allows it to effectively receive and integrate information from various sources^55^. Our data shows that following a global inter-site coherence of LF phase, the sites in the ANT were the first to initiate HF activity, which may suggest that the ANT plays a key role in the transition to memory processing following the HPC binding in the LF range (**Fig 4**). Importantly, our data supports the notion that the ANT contributes uniquely to memory processing, beyond duplicating hippocampal functions^19,56^. Further support of ANT’s presumed integrative role was given by our results that the stronger the HF functional connectivity between an ROI and ANT, the more likely that the ROI was to evoke a response in ANT following electrical stimulation (**Fig 5d**). When we stimulated a given ROI, the ANT were consistently the most likely to synchronize their activity with the stimulated region (**Fig 5**) and the first to respond to the stimulation (**table S12**). This further emphasizes the active role of the ANT in coordinating and integrating neuronal activity across the brain^57^. However, we are mindful that the specific role of the ANT may vary depending on the context and other factors, suggesting that more research is required to fully understand the ANT’s involvement in memory processes.

### PMC and Autobiographical Memory

Another important observation in our study pertains to the finding of simultaneous engagement of HPC and PMC and the strong effective connectivity between them highlighting a direct and strong functional and anatomical relationship between the HPC and PMC beyond the known Papez circuit and bypassing the ANT during retrieval of stored personal memory information. This is in agreement with our own recent electrophysiological findings showing strong coupling between MTL and retrosplenial cortex ^58^ and anatomical tracing studies in the primate brain documenting strong direct connections between the MTL and PMC ^59^. Of note, there is evidence that approximately half of HPC projections to the mammillary bodies in the rodent brain send direct collateral projections to the PMC^60^. Lastly, our findings lend support to a recent MEG study, in which inhibitory stimulation of the human PMC led to a network-level alteration of MTL-driven oscillatory coupling with the PMC itself and with other posterior cortical structures^61^.

### OFC and Autobiographical Memory

Consistent with the current ideas about the prefrontal cortex being important for schema-related processes^13,62,63^, we expected to find stronger HF activity (denoting increased averaged neuronal firing) in the OFC. Instead, we found enhanced HF activity in the ANT (even before the HPC), which raises an intriguing new hypothesis: a yet different system of the brain, that was not included in our study, may be driving the ANT HF activity and inducing an “episodic-retrieval mode”. Given the known anatomical connections of the ANT in rodents and primates^19^, candidate structures may include medial PFC areas (located more dorsal to the orbital and ventral regions that we studied here). In keeping with this, in an optogenetic study it was shown that the rodent prefrontal (anterior cingulate) neurons are causally important for inducing contextual memory retrieval^64^; studies in primates have shown that, in the absence of bottom-up visual inputs, the prefrontal cortex is causally important for recall^65^; and a recent MEG study^66^ used source reconstruction to examine the changes in 1–30 Hz power during AM retrieval and found engagement in the medial PFC above the level of the corpus callosum and including the anterior cingulate area.

### Limitations and Future Directions

Our study has several important limitations. First, we focused only on 4 ROIs primarily because of lack of sufficient coverage in other ROIs as the sites of recordings had been motivated by the clinical needs. We acknowledge that autobiographical processing involves additional regions beyond the four ROIs studied here. For instance, angular gyrus is another key region of interest ^15,67–69^, where we have previously reported^70^ similar activations during autobiographical processing. Second, we treated the ROIs as if each ROI is a unitary functional region. Different populations of neurons within the anatomical boundaries of a given structure may play very different functional roles. We have already documented a remarkable heterogeneity of responses within the boundaries of PMC^71^ and recently several subdivisions within the PMC have been suggested by others^72^. Additionally, we grouped posterior and anterior HPC together while our own recent observations have revealed autobiographical memory related ripples appearing clearly stronger in the anterior than posterior HPC^39^. Prior work has also claimed differential activity along the anterior-posterior axis of the structure^73^. A potential avenue of future research would be to compare timing and connectivity analyses across different populations of a given ROI. Lastly, as we have noted, the process of autobiographical memory processing - instead of being a memory pure process – blends several kinds of self-referential constructive processes^5,38^. As such, one may not expect autobiographical memory experimental paradigms to probe episode-specific and memory-pure processes as other lab-based experiments do. However, these statements should not be taken to imply that the electrophysiological responses reported in our work are not related to autobiographical memory processing as they clearly tracked with the subject’s behavior in several ways: Hit trials eliciting significantly higher physiological responses than non-autobiographical fact statements or correct rejection AM trials (Fig 3). Moreover, it is widely believed that inferential reasoning and imagination processes should occur more during correct rejections than during correct hit statements. Yet, we showed statistically stronger HF responses during hits vs correct rejections. Future studies are needed to explore the causal and differential contribution of each of the ROIs (and their sub-regions) in different aspects of self-referential processing.

In closing, we hope that a more informed knowledge about the temporal cascade of physiological events and their location within the circuitry of AM-network may prove essential for designing future neuromodulation studies to test to change or enhance retrieval processes through careful manipulation of cross-regional oscillatory dialogues.

## Data availability statement

All data will be available for sharing upon reasonable request.

## Code availability statement

The code used to derive the connectivity index is available here: https://github.com/JRStieger/ConnectivityIndex/

## METHODS

### Participants

Thirty-one subjects (age range: 19-52 years old, sixteen female, **Table S1**) with refractory epilepsy were implanted with stereotactic electrodes (sEEG) to localize their seizure onset zone. Compared to neuroimaging studies, clinical invasive recordings in humans have clear limitation in terms of small number of recruited subjects and sparse coverage across the brain within each individual. We sought to overcome the sparsity of the sEEG method by aggregating data across a cohort of patients. All the participants had at least one electrode site placed over at least two of the regions surrounding the hippocampus (HPC), posteromedial cortex (PMC), orbitofrontal cortex (OFC), and/or the anterior nuclei of the thalamus (ANT).

Moreover, the location of electrodes in each participant were decided purely by clinical evaluation since the invasive procedure of intracranial EEG was primarily conducted for pre-surgical clinical purposes. As our data were recorded in subjects suffering from epilepsy, this may be a clear limitation. We note, however, that there is strong evidence documenting that non-lesional epileptic tissue exhibits normal physiological responses to incoming relevant stimuli, which are “seized” or diminished only at the time of an ongoing epileptic pathological activity^74,75^. Given the number of trials in each condition, we believe only a minority of trials were presented during epileptic discharges. Moreover, in each patient with implanted electrodes, only a minority of electrodes (<20%) show epileptic activity, while the majority of electrodes show no signs of epileptic discharges^76^. To mitigate the confounding effects of epilepsy and seizures, we recruited patients with focal seizures who do not have diffuse brain disease and conducted our experimented outside the window of seizures. Moreover, we reviewed the EEG tracings of the patient with the clinical team prior to testing and made sure that the patient did not have subclinical or clinical seizures prior to testing.

Each subject was monitored in the hospital for approximately 6 to 10 days after the implantation surgery. The Institutional Review Board of Stanford University approved the experiments and all subjects provided verbal and written informed consent before experiments. All 31 participants completed the sequential task and 27 participants completed the simultaneous task (tasks described below).

### Electrode Localization

In order to precisely determine the anatomical locations of the electrode recording sites, we co-registered a structural MRI brain scan with a post-operation CT scan to highlight the precise location of the electrode contacts. A high-resolution T1 was performed on a 3T GE scanner. The scanning parameters were as follows: 256 × 256 matrix, 186 slices, 0.90 ×0.90 ×0.90 mm voxel size, 240 ms field-of-view, 7.60 ms TR. To reconstruct the cortical surfaces, the T1 image was processed via FreeSurfer (recon-all command)^77^. The post-implant CT image was co-registered to the space of the high resolution T1 volume. For each individual, the electrode locations were manually labelled on the co-registered CT using BioImage Suite^78^ then projected to the individual reconstructed 3D Brain by iElVis toolbox^79^.

Locations within the anatomical boundaries of HPC, PMC, OFC, and ANT were chosen as the regions of interest and contacts falling within these boundaries were identified by a trained medical professional. Electrodes falling within any of the regions of interest (in native anatomical space) were selected for this study, which resulted in 240 electrodes in the Hippocampus, 234 in the PMC and 320 in the OFC across all subjects (**Table S3**). The 13 most recent patients, in addition to having electrodes placed in the HPC, PMC, and OFC additionally had electrodes implanted in the anterior nuclei of the thalamus (ANT) (47 total electrodes across the ANT) i.e., in the anterior part of the thalamus between the limbs of the internal medullary lamina. Note: due to the heavy computational burden of some analyses, the most recent patients with ANT electrodes were not included in some analyses that were sufficiently powered and simply focused on OFC, PMC, and HPC (**Figs 2-4**).

The Individual ANT contact’s center of mass was defined in native T1 space for each subject. A linear affine transformation of each subject’s T1 image to MNI space was performed to convert center of mass of each contact into a scalar X,Y,Z coordinate in MNI space. The THOMAS atlas^80^ was used to define the left and right thalamus, as well as the left and right Anteroventral (AV) nuclei constituting each Anterior Nucleus of the Thalamus (ANT) in the THOMAS atlas. A contact neighborhood was created by assigning a 0.5−0.5−0.5mm voxel space around each contact center of mass in MNI space. Each contact neighborhood was then analyzed against the entire voxel space of the AV nucleus. For each contact, a Euclidean distance in millimeters MNI space was computed by taking the minimum of the Euclidean distance between each voxel of the contact neighborhood and each voxel of the AV nucleus (**Figs S1f**).

To mitigate the problem of sparse recordings in each individual and un-identical coverage across individuals, in the analysis of regional activity we included *all* individuals who had coverage in that particular ROI, but very importantly, for the analysis of cross-regional analysis, we only analyzed pair-wise interactions across two ROI *within the same individual* when both ROIs were covered.

### Experimental Design

In order to closely monitor the neural dynamics of autobiographical memory (AM) processing we designed two tasks that leveraged the precise temporal resolution of the sEEG method and contrasted AM condition with another inwardly directed cognitive task (arithmetic calculation). In these tasks, arithmetic calculation, or “Math” represented a distractor rather than a perfect control condition.

#### The Simultaneous Presentation Task

Required subjects to make true/false judgments on a series of autobiographical memory (AM) statements (e.g., “I drank a beer this week”) and arithmetic statements (e.g. “33+5=38”), which were visually presented in the center of the laptop monitor (**Fig 1a**-upper). Subjects indicated their response by pressing one of two keypad buttons. AM and math trials were randomly shuffled and lasted until the subject’s response, or for 15s for trials where subjects made no response. Each subject performed between 50 - 120 trials for each condition. A 700 ms inter-trial interval (ITI) separated consecutive trials and was also used as a baseline period for several analyses. The Simultaneous Task also included a third condition with fact statements (e.g., “Paris is in Europe”). These statements were not presented in the Sequential Task. Therefore, we did not include them in our analysis. Lastly, the task also included a rest condition during which a cross hair appearing at the center of the screen when subjects were instructed to rest. These trials were excluded from the analysis. For trial numbers, see **Table S2**.

#### The Sequential Presentation task

Also required subjects to make true/false judgments on a series of AM and arithmetic statements. However, in this task, each statement was separated into four AM (e.g., Last year, I, took, a test.) or five non-AM statements (e.g., “Fifty-seven”, “plus”, “seven”, “equals”, “sixty-four”) fragments. Hence S1-4 in AM condition and S1-5 in non-AM condition in **Fig 2**. This allowed us to track activity across the different stages of sentence and AM processing. Each statement fragment was presented for 500 ms followed by a 500 ms inter-stimuli-interval (**Fig 1a**-lower). A 700 ms inter-trial-interval (blank screen) separated consecutive trials. Trials were organized into blocks of 12 consecutive AM or math trials, but the order of blocks was randomly shuffled for each subject. Each subject performed 48 - 96 trials for each condition. As in the simultaneous task, participants indicated their response to each statement by pressing one of two keypad buttons. Further description of task stimuli can be found in **Table S2.**

Both the simultaneous and sequential tasks were programmed via Psychtoolbox^81^ in MATLAB (http://psychtoolbox.org/) and were run on an Apple Macbook Pro or HP laptop, which was placed ∼70 cm in front of subjects, while they were sitting up in their hospital bed.

### Intracranial EEG data acquisition and preprocessing

Data were collected using two multichannel recording systems (Tucker Davis Technologies, sampling rate 3051.76 Hz for subjects 1-3; and the Nihon Kohden recording system, sampling rate 1000 Hz, for subjects 4-26). Stimulus onset times of the visually presented task stimuli were marked via a photodiode for participants 1-3, and via an RT box for participants 4-26 and were synced with the iEEG signals.

Following data collection, a preprocessing pipeline was implemented to remove noise from the electrophysiological signals with as little distortion as possible. First, signals above 1000 Hz were downsampled to 1000 Hz. Next, we applied spatial and notch filters centered at 60, 120 and 180 Hz to the downsampled signals, to remove electrical line noise^82^. We then identified noisy channels to be excluded from subsequent analyses. Noisy channels were defined as those whose raw amplitude was larger than 5 times or less than one-fifth of the median raw amplitude across all channels, or that exhibited more than 3 times the median number of spikes across all channels (spikes defined as jumps between consecutive data points larger than 80 μV). After identifying pathological and noisy channels, the signal from each site was re-referenced to the average of the non-noisy channels (common-average referencing).

We are mindful that the procedure of common average effectively increases the extent of each electrode leadfield, and may introduce spurious correlativity, particularly between nearby contacts in posterior HPC and ventral PMC^83,84^. We performed quality measures to demonstrate that our core findings are not an artifact of the referencing scheme. We observed qualitatively similar results with common average and bi-polar referencing, and believe our results would generalize across referencing choice (see supplemental material and **Fig S14**).

Different cognitive operations have been found to be correlated with activity in different frequency bands of the raw voltage signal. Therefore, frequency decomposition of each re-referenced signal was performed using Morlet wavelet filtering (at log-spaced frequencies between 1 and 256 Hz [59 total frequencies]; each wavelet having a width of 5 cycles)^85^. We extracted the instantaneous power at each frequency, then z-scored the power for each frequency separately, across the time dimension.

### Selection of AM-related Features

In order to determine which electrophysiological features could be consistently detected across subjects during AM we used Cluster Based Permutation Tests (CBPT)^86^ to evaluate whether there were significant changes from baseline in these features that were locked to particular clusters in the time-frequency feature space. Subject responses were averaged across a given ROI, then baseline corrected. Clusters were identified by finding pixels in the time-frequency space that T-tests across subjects (Or electrodes, for ANT) determined were significantly different from baseline (−500:−200ms before the first stimulus; alpha = 0.05). T-values were summed across the identified clusters. Subject responses were then circularly shifted 5000 times in frequency and time (preserving their structure) and the maximum (and minimum) cluster t-value sum was recorded for each iteration to create a null distribution. Clusters more extreme than 95% of the null distribution clusters were determined to be significant (Bonferroni corrected). All subjects (N=31) were included in this analysis.

Once the significant time-frequency clusters were identified, we sought to find which channels and connections displayed these particular features as well as how task dynamics effected these features. For power features, time-frequency pairs were averaged across two bands in each trial: low frequency (LF: 1-6 Hz) and high frequency broadband (HF: 70-170 Hz) power. It should be noted that we selected the 1-6Hz range of LF activity instead of focusing purely on the theta band (4-7Hz) because our initial observations as presented in **Fig 2** showed a significant change in this broader LF range beyond the narrower band of theta oscillations. As noted in the main text, we confirmed the presence of oscillations in the LF range in the power spectrum plots in the form of peaks above the aperiodic background - utilizing the same method as described elsewhere ^40^.

Memory related (AM related) channels were identified by significant increases in HF power above baseline and significantly greater HF power during AM trials compared to math trials in at least one of the two tasks. Permutation tests determined the channels that displayed significant activations above baseline by permuting the value labels (baseline vs trial values). The same process was used to identify AM selectivity by permuting the trial labels (math vs AM trials). The baseline window was defined as 500ms-200ms before first stimulus while the trial window was defined as 500ms-1500ms following last stimulus (where the greatest density of HF power was observed, see **Figs 2,5**). This window was chosen due to it containing the highest density of HF power, however similar results were observed with response locked analyses. To mitigate false discoveries, all channel and connection p-values were Benjamini-Hochberg corrected^87^. All statistical tests used 5000 permutations. All future analyses focused on channels that we identified as AM related.

### Memory Content Modulation

To test whether HF power was consistently modulated by the AM statements observed, we averaged the stimulus-locked trial by trial response in the window defined above (500ms-1500ms following the last stimulus) across different memory categories. Statements presented to the subjects were grouped according to *memory age* (recent, remote, fact), *subject response* (did subjects respond “True” or “False”?), *memory content* (were the statements assumed to be interpreted as pleasant, unpleasant, or neutral?), and *verb structure* (active or generic verbs, i.e., a verb indicating an action without an object (e.g., kicked) vs. verbs that needed an object to illustrate an event (“had” + a beer or + headache). We note that this analysis was not designed to test how our ROIs processed the semantic categories listed above, but rather to highlight that the content of the memories (inferred from the different semantic categories of the statements) influences activity in the HF band. This analysis was performed on data collected from the first 26 subjects before we recruited additional five subjects with thalamic coverage to enhance the statistical power of our thalamic data.

### Phase Coherence

Phase coherence, or phase synchrony is thought to capture the consistency in brain responses across trials or time. Intertrial Phase Coherence (ITPC) is a measure of similarity of phase response within an electrode across trials. Higher values of ITPC will be found when electrodes present consistent phase profiles in response to stimuli or cognitive operations. Intersite Phase Coherence (ISPC) measures the difference in phase angles between two electrode sites and is a weak measure of functional connectivity. Higher values of ISPC will be found when the phase of signals in two different are more synchronized (displaying a consistent offset in phase angle). We used ITPC as a measure of the reliability of the response in a given site to an incoming stimulus and ISPC as a measure of how reliably two different regions respond to the same stimuli or cognitive operation. ITPC was calculated by taking the complex phase component of the wavelet decomposition, averaging this value across trials within each channel, and then taking the absolute value^88^. ISPC was calculated by taking the difference between the complex phase of two separate channels and then averaging this value across trials and then taking the absolute value ^12^. Within region ISPC was only examined if a patient had electrodes in both left and right hemispheres. Each channel’s ITPC and ISPC values were normalized by a permutation distribution (500 permutations) of circularly shifted phases (ITPCz, ISPCz). Cluster based permutation tests were computed across subject averages as above to determine significant increases in ITPC and ISPC. To estimate timing of ISPC during individual trials, ISPC was calculated across a 500ms window shifted with 99% overlap. This analysis was based on the cohort of 26 initial subjects recruited to our study. Again, we performed posthoc tests to lend support that our ISPC results were not driven by volume conduction or referencing choice (see supplemental material).

### Response onset latency (ROL)

In order to infer the flow of information across regions, we estimated the response onset latency (ROL) of task-related activations at each electrode site of interest at the single trial level using a method reported previously in our group^70^. See supplemental figure **Fig S7** for a visual description of the method. Time-frequency signals were first averaged across the frequencies of interest mentioned above. Neural signals, namely low-frequency (LF) power, Inter-Site Phase Coherence (ISPC), and high-frequency (HF) power, were initially smoothed by applying a 100ms moving average. Subsequently, a sampling process was employed using 10,000 random windows, each of 300ms duration, to compute the average expected signal value across these windows. The averages obtained from the 300ms windows were utilized to define a permutation distribution, representing the expected average value across a 300ms window. A threshold was then established at the 95th percentile of this distribution. Significant events were identified as contiguous timepoints where the signal amplitude exceeded the threshold for more than 100ms. The response onset latency (ROL) was then defined for each of these significant events. This was computed as the average time of the event, with the calculation being weighted by the signal amplitude. For each channel, its average ROL was determined as the average over all significant events for that channel that coincided with signal detection in another ROI within the same hemisphere for the given trial. In essence, for a significant event to contribute to the average ROL, it needed to be detected concurrently in at least two different ROIs within the same hemisphere and during the same trial. This analysis was based on the cohort of 26 initial subjects recruited to our study.

### Single Pulse Electrical Stimulation (SPES; STIM)

While our results suggested the ROIs examined coordinate their activity during AM, we used Single Pulse Electrical Stimulation (SPES; referred to as STIM for ease of understanding) of the brain as a strong and causal measure of the connectivity of two brain regions. Otherwise known as the study of corticocortico evoked potentials ^44^, we recorded STIM by electrically stimulating one ROI while recording from all other distal contacts. The electrical stimulus consisted of a 6mA constant current, biphasic square-wave pulse lasting 200us at 0.5Hz totaling 39-40 pulses for each electrode pair. Raw voltage traces were computed by time locking to the electrical stimulus.

In order to ensure our results were not driven by stimulation artifacts, we employed an automatic artifact rejection scheme to account for the off chance that a minor artifact could still be present in the bipolar traces. To automatically remove these artifacts, we first examined a 40ms window surrounding each stimulation pulse for extreme differences in voltage values. We examined the derivative of the signal for extreme values under the assumption that physiological changes in voltage would proceed more smoothly while the gradient would be much sharper for artifactual changes. If we detected derivative peaks within this window that exceeded 99% of the voltage derivative distribution, these data points were removed and interpolated with an autoregressive moving average (fillgaps in MATLAB). After automatic artifact rejection, we manually checked the traces of all evoked responses and excluded those displaying a stimulation artifact, while keeping those without a significant artifact (see **Fig S10** for further details).

STIM activation was defined using a subject specific thresholding procedure. The MATLAB findpeaks function was used to define a distribution of the raw voltage and prominence of all peaks of the time-locked average voltage traces within the first 500ms following stimulation across all stimulation-recording pairs. Significant physiological voltage deflection caused by electrical stimulation resulted in high voltage and prominence values, which decayed across time (most physiological responses occur within the first 200ms). Significant activation in a stimulation-recording channel pair was determined when a peak in the average voltage trace exceeded 2 standard deviations of the raw voltage and prominence distributions. Since the early sharp negative potential (referred to the N1 potential) is often less prominent than the later slow-wave like N2 potential^44^ when a stimulation-recording pair was found to have a significant activation, the onset of the N1 potential (or first detected voltage deflection) was defined as the first detected peak that exceeded 1 standard deviation of the voltage and prominence distributions.

We then defined a ***connectivity index*** that attempted to capture how reliably electrical stimulation produced responses in other brain regions by combining STIM and ITPC (See **Fig S11** for detailed methods).

The derivation of the connectivity index proceeded as follows: We first automatically removed potential stimulation artifacts by examining a 40ms window surrounding each stimulation pulse for extreme differences in voltage values (**Fig S10**). We chose to examine the derivative of the signal for extreme values under the assumption that physiological changes in voltage would proceed more smoothly while the gradient would be much sharper for artifactual changes. If we detected derivative peaks within this window that exceeded 99% of the voltage derivative distribution, these points were removed and interpolated with an autoregressive moving average. This data was then decomposed with a wavelet transform, and the intertrial phase coherence for each stimulation/recording electrode pair was calculated across stimulation trials. As can be seen in **Fig S11**, a prominent cluster of ITPC was observed in response to stimulation. A cluster-based permutation test across all stimulation/recording electrode pairs identified the boundaries of the average significant change in ITPC in response to stimulation. Then, ITPC values within this cluster were summed across the time-frequency values in each stimulation/recording pair which provided a single metric to quantify the effective connectivity strength between two pairs of electrodes. We then fit mixed effects models to the effective connectivity strength to determine direction of influence as measured by electrical stimulation (**Figs 6b,c**). The same procedure was used to calculate the stimulation induced intersite phase coherence.

Correlation analyses compared this connectivity index to functional connectivity (FC) measures during AM (**Figs S12-13**) using Pearson’. FC was calculated across a 2s window with the leading edge defined by the subject’s response. The two measures examined were HF power correlation, the correlation between the average HF power in two ROIs across trials, and ISPC, the intersite phase coherence across time in the LF range (1-6Hz) between two ROIs averaged across trials. As can be seen in **Fig 5d** and **Figs S12-13**, arrows were drawn when significant correlations were observed between memory task engagement and the directional causal connectivity index *across both tasks*.

### Phase Amplitude Coupling

We calculated phase amplitude coupling (PAC) similar to Tort et al., to examine the relationships between LF phase and HF power^48^. We first examined whether there were significant cross regional effects of phase amplitude coupling. We extracted the phase of the LF signals 1-6Hz and binned the HF power from different ROIs in 18 uniform bins spanning −π to +π. We then computed cluster-based permutation tests across channel connections for the 13 subjects with ANT connections to determine whether there were consistent LF phases in one ROI that produced significantly more or less HF power in a different ROI than would be expected by chance. The permutation distribution was calculated by using the LF phase from one trial to bin the HF power in a different randomly selected trial. Once this was established, we extracted the PAC range for each channel pair, defined as the maximum HF power across LF phase minus the minimum HF power across LF phase. Mixed effects models (described below) then compared the strength of these effects. These analyses were based on 13 subjects with ANT coverage.

### Lagged Correlation

We used lagged correlation as directed measure of functional connectivity to corroborate some of our response onset latency results. The two signals we examined were LF phase and HF power (HF power was smoothed by 100ms). In both cases, we computed the lagged correlation of these signals across channel pairs for different ROIs in the 13 subjects with ANT contacts. We then used cluster-based permutation tests as described above for the PAC analysis to determine whether there were lags of significantly more or less correlation between these signals than would be expected by chance. In both cases, we found a peak significantly greater correlation than expected by chance with, on average, the maximum correlation between signals at zero lag. We then examined each channel pair, and if we found a positive peak that included the origin (zero lag), we extracted the positive values of the peak, then integrated the correlation values to the left of the peak (negative lag; reference signal leads other ROIs) and correlation values to the right of the peak (positive lag; reference signal lags other ROIs). Finally we used T-tests to determine whether we found evidence of significant leads or lags between ROIs in these two signals. These analyses were based on 13 subjects with ANT coverage.

### Statistical Analysis

One limitation of the statistical analysis of sEEG signals is that electrodes are not uniformly distributed in all brain regions. Mixed Effects Models (MxM) are able to perform regression analyses while accounting for differences in group size. We used MxMs to examine how electrophysiological features are related to the interaction between ROIs and cognitive content, task demands, and functional connectivity. Mixed effects analysis was performed in R using the LME4 package^89^. The LmerTest package was used to calculate p-values of the mixed models^21^. Outliers were removed with the Grubbs’ method in Matlab. The unit of analysis was typically individual patients, but in some cases, the unit of analysis was individual electrodes (see main text). Data were fitted with a random intercept model with the relevant fixed factors and a random factor of ‘Electrode’ nested within ‘Patient’. Fixed effects structures of the mixed-effects models were reduced stepwise by excluding nonsignificant interaction terms/predictors and compared using ANOVA ratio tests until the respectively smaller model explained the data significantly worse than the larger model (significant χ2-test)^90^. Follow-up tests were run by comparing marginal means using the emmeans package^91^. AM related power responses were fit with the model:

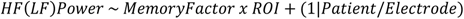

The memory factors examined were task condition (Levels: Math and AM), memory age (Levels: Recent, Remote, and Fact (fact only in simultaneous task)), subject response (Levels: True and False), memory content (Levels: Pleasant, Unpleasant, and Neutral), and verb structure (Levels: active, generic). The regions of interest (ROIs) examined were OFC, HPC, and PMC. The same analysis was performed on the trial level for one subject with AM selective ANT electrodes.

Response onset latency was fit with the model:

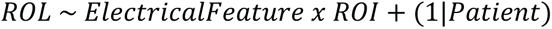

The electrical features examined were LF power, HF power, and ISPC over time. ROIs included OFC, HPC, PMC, and ANT.

Effective connectivity analyses were fit with the model:

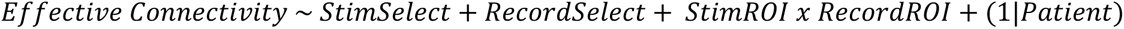

Two models were fit. The first was based on the ITPC connectivity index, and the second was based on the stimulation induced ISPC. Due to the large number of connections for the second model, each stimulated site had one value that averaged all the recording site pairs. StimSelect and RecordSelect were 1 if the electrode in question displayed AM related activity. Finally phase amplitude coupling analyses were fit with the model:

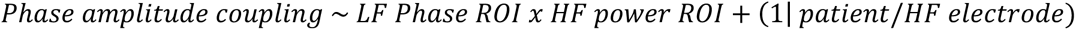

## SUPPLEMENTARY MATERIAL

### Volume Conduction and ISPC

To determine the possible influence of volume conduction on our Inter-Site Phase Coherence (ISPC) results, we initiated an examination by calculating the average minimum distance between all pairs of contacts across different Regions of Interest (ROIs). The distance values were found to vary among different pairs of ROIs. The average minimum distances (± standard deviation), in millimeters, between pairs of ROIs are as follows:

- HPC to ANT: 32.74 ± 2.65
- HPC to OFC: 50.02 ± 8.39
- HPC to PMC: 49.55 ± 12.17
- OFC to ANT: 46.35 ± 10.56
- OFC to PMC: 79.53 ± 9.66
- PMC to ANT: 35.13 ± 6.99

The magnitude of these minimum distances suggested a limited likelihood of volume conduction distorting our ISPC results. However, to further consolidate this proposition, we explored a potential correlation between distance and ISPC values for the two ROI pairs presenting the smallest average minimum distances, namely, HPC to ANT and PMC to ANT.

For the Sequential (Seq) task and the Simultaneous (Sim) task, we found no significant correlation between distance and ISPC values in either HPC_ANT (Seq task: r(220)=-0.0,p=0.969, Sim task: r(220)=-0.07,p=0.29) or PMC_ANT (Seq task: r(151)=0.07,p=0.43, Sim task: r(151)=0.11,p=0.18) pairs, reinforcing our earlier confidence that our results were not predominantly influenced by volume conduction.

It should be noted that, in certain instances, there were no ipsilateral pairs of ROI contacts. Thus, these average minimum distances should not be interpreted as a precise representation of the patients’ actual anatomical structures, but rather as a useful metric for our analytical purposes.

### Impact of Referencing Method on ISPC Results

In order to discern the potential influence of the referencing choice on our ISPC results, we conducted an analysis focusing on our thalamus subjects. As our ISPC results were more pronounced in the Sequential (Seq) task, this task became the focus of our analysis. The aim was to examine if our ISPC results were artifacts or driven by the choice of referencing.

A comparison was made between the two referencing methods: common average referencing (AVG) and bipolar referencing (BP). The t-tests showed that, overall, the strengths of ISPC results with AVG were qualitatively similar to those with BP. The following table summarizes the test results:

| ROI Pair | AVG Result | BP Result |
| --- | --- | --- |
| **HPC_ANT vs HPC_OFC** | t(147)=8.75,p<0.0001 | t(147)=7.03,p<0.0001 |
| **HPC_ANT vs HPC_PMC** | t(89)=6.22,p<0.0001 | t(89)=3.73,p=0.0003 |
| **HPC_ANT vs OFC_ANT** | t(208)=6.02,p<0.0001 | t(208)=3.36,p=0.0009 |
| **HPC_ANT vs OFC_PMC** | t(180)=10.18,p<0.0001 | t(180)=7.54,p<0.0001 |
| **HPC_ANT vs PMC_ANT** | t(158)=2.04,p=0.0432 | t(158)=5.83,p<0.0001 |
| **HPC_OFC vs HPC_PMC** | t(136)=-0.23,p=0.8156 | t(136)=-1.68,p=0.0951 |
| **HPC_OFC vs OFC_ANT** | t(255)=-4.85,p<0.0001 | t(255)=-2.96,p=0.0034 |
| **HPC_OFC vs OFC_PMC** | t(227)=0.72,p=0.4724 | t(227)=0.60,p=0.5524 |

Given these findings, we conclude that the choice of referencing does not significantly influence the ISPC results in the Seq task. This consistency across different referencing methods supports the robustness of our ISPC measurements.

## Author contributions

JRS contributed to the acquisition, analysis, and interpretation of data and creation of new codes and measures used in the work and prepared the first draft of the manuscript; PRC contributed to the acquisition, analysis, and interpretation of data; YF contributed to the analysis and interpretation of initial set of data; ZL and CMP contributed to the acquisition of data; AW, DC, QC, and JH contributed to interpretation of data; VB contributed to surgical operations and data collection and interpretation of data; All authors made comments on the manuscript. JP designed the study and supervised all aspects of the work.

---

**Fig S1.**
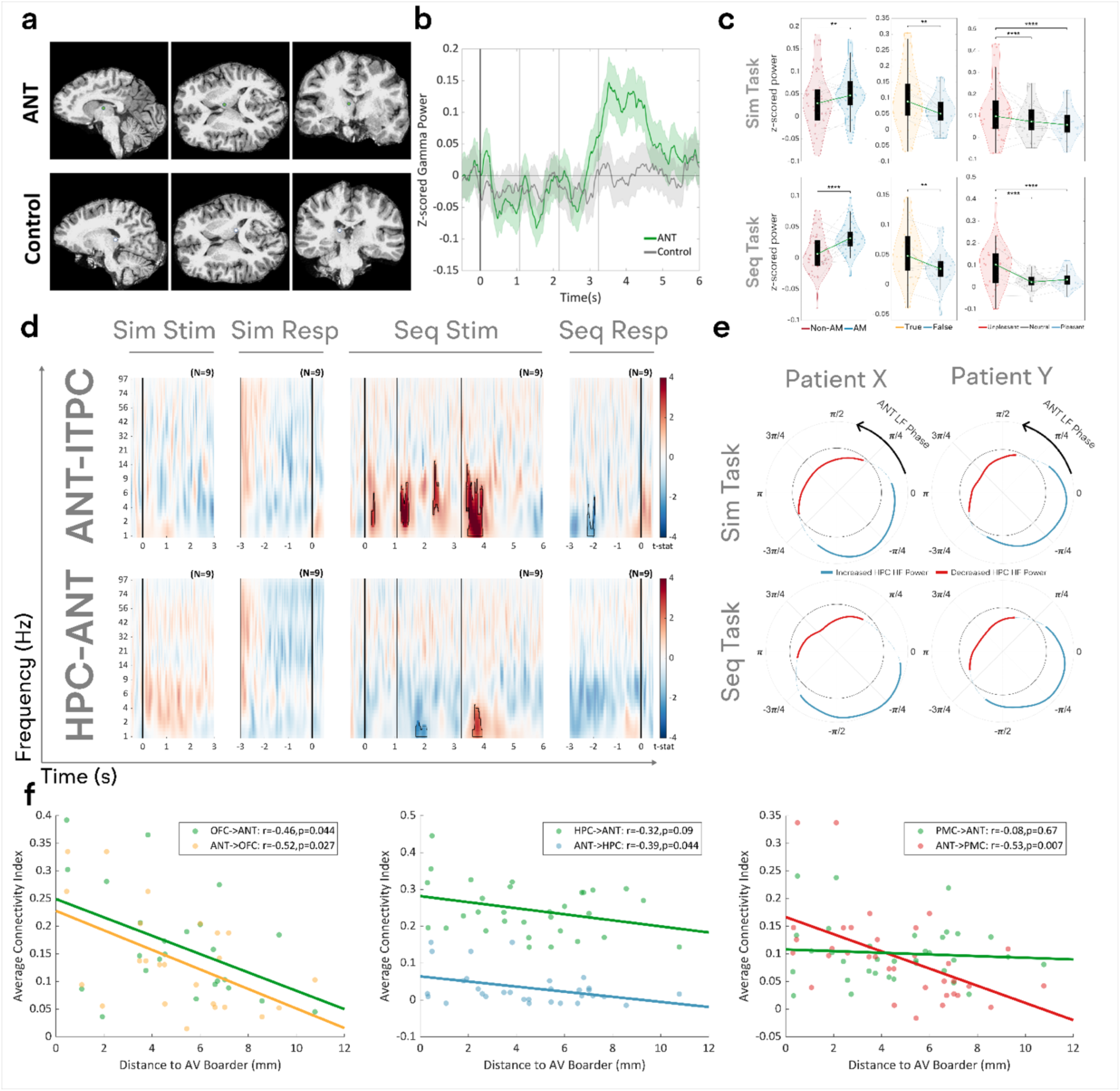
Human thalamus recordings. **a**) Anatomical locations of ANT electrode (upper) and ipsilateral control thalamic electrode (lower) in the same patient. **b**) The average gamma power (32-58 Hz) during AM trials in the Seq task shows that the AM-related responses are specific to the ANT anatomical location. **c**) Group level ANT HF modulation by trial type. (Left) On average ANT displays more HF power during AM trials compared to Non-AM trials (MxM:T(80)=4.75,p<0.0001). (Middle) When selecting the AM specific electrodes (either activated above baseline or memory active) ANT electrodes display similar patterns of trial type HF modulation as the other ROIs such as increased activation for true vs false trials (MxM: T(30)=3.23,p=0.003), and unpleasant vs neutral trials (MxM: T(60)=5.45, p<0.0001). However, the effects for recency (MxM: T(30)=1.62, p=0.21) and verb structure (MxM: T(30)=0.71, p=0.41) were not significant. **d)** Each panel displays the group-level stimulus locked (Stim) and response locked (Resp) change in ANT phase coherence from baseline during the simultaneous (Sim) and sequential (Seq) presentation tasks during AM condition. Within electrodes (intertrial phase coherence; ITPC-First row), ANT also shows increased phase coherence compared to baseline in the low to mid frequency range (∼1-20Hz) across trials following each stimulus, while the intersite phase coherence (ISPC) between ANT and HPC was strongest in the LF range (∼1-6Hz) after the last stimulus i.e., during the retrieval process (see **Fig S5** for all pairwise comparisons). All channel pairs for a given connection type were averaged for each subject and statistics were performed on the group level (permuted across subject averages). Both ITPC and ISPC were calculated across trials. Statistical significance was determined using cluster-based permutation tests (CBPTs). Significant time-frequency clusters are highlighted and outlined in red and blue (ITPC: all p<0.003, ISPC: all p<0.002, Bonferroni corrected for 3 connection types). **e**) Significant phase amplitude coupling between ANT LF phase (1-6Hz) and HPC HF power was observed across multiple subjects. For these plots HPC HF power was averaged across 18 uniform bins spanning −π to +π of the ANT LF phase. Thick blue lines represent the ANT phases where HPC HF power is significantly increased, and thick red lines, decreased compared to a permutation distribution where HPC HF power was binned according to ANT LF phase from a different randomly selected trial (CBPT, p<0.001). Significant effects were seen at the same phases for the Sim (upper) and Seq (lower) tasks. **f**) The average connectivity index (see **Fig S11**) between ANT electrodes and other ROIs significantly correlated with the Euclidean distance in MNI space of each thalamic contact to the border of anteroventral (AV) nucleus. Correlations were corrected for multiple comparisons (6 connection types).

**Fig S2:**
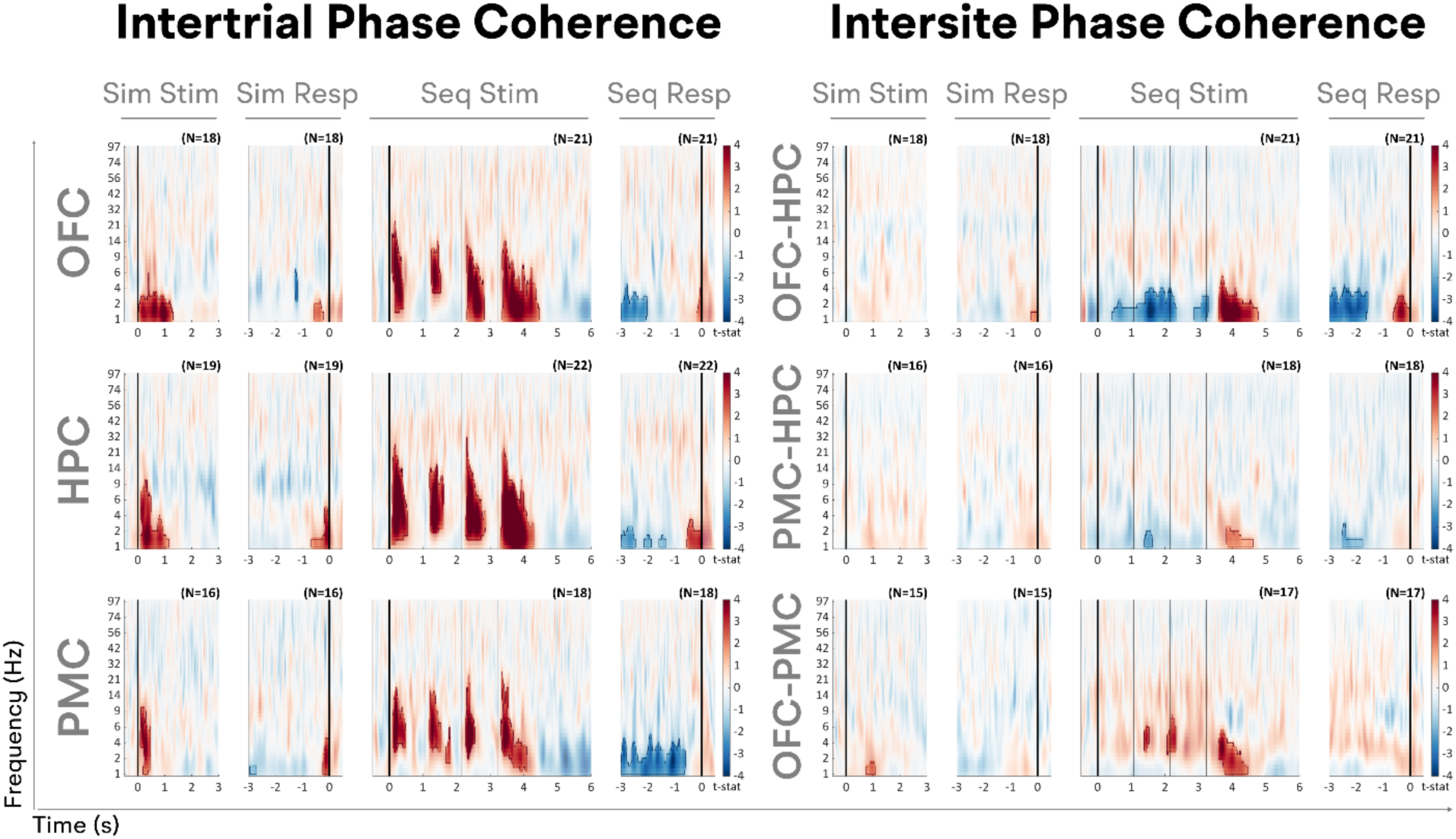
Unique patterns of phase coherence across trials and sites. Each panel displays the group-level stimulus locked (Stim) and response locked (Resp) change in phase coherence from baseline during the simultaneous (Sim) and sequential (Seq) presentation tasks during AM processing. Within electrodes (intertrial phase coherence; ITPC), each ROI shows increased phase coherence compared to baseline in the low to mid frequency range (∼1-20Hz) across trials following each stimulus, while across electrodes of different ROIs (intersite phase coherence; ISPC), increased phase coherence was strongest in the LF range (∼1-6Hz) after the last stimulus i.e., after the cue stimulus. As seen in **Figs S2-5**, similar findings were also seen during the non-AM (math) condition suggesting that the ITPC (after each stimulus; **Fig S2**) and between-sites ISPC (after the last stimulus; **Fig S4**) synchrony events contribute to both conditions. ITPC was calculated individually for each electrode and averaged across electrodes in a given ROI within each subject. ISPC was calculated between different ROIs, but the same results were seen within the same ROIs, but across hemispheres (**Fig S4**). All channel pairs for a given connection type were averaged for each subject and statistics were performed on the group level (stats computed across subjects). Both ITPC and ISPC were calculated across trials. Statistical significance was determined using cluster-based permutation tests (CBPTs). Significant time-frequency clusters are highlighted and outlined in red and blue (ITPC: all p<0.003, Bonferroni corrected for 3 ROIs, ISPC: all p<0.002, Bonferroni corrected for 6 connection types). Similar coherence results were observed in/with ANT contacts; see **Figs S1d,S5**.

**Fig S3:**
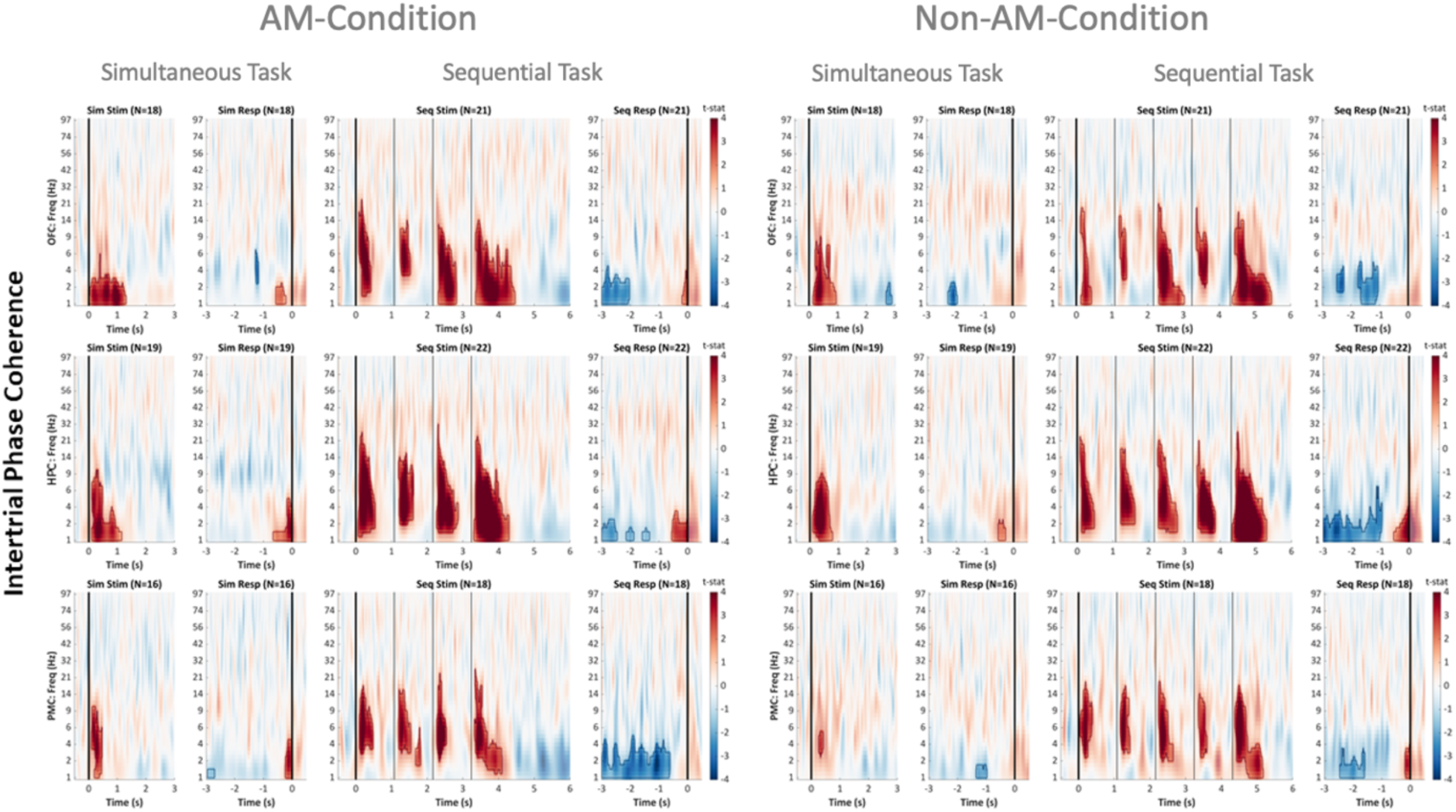
Local phase coherence. Each panel displays the group-level stimulus locked (Stim) and response locked (Resp) change in phase coherence from baseline during the simultaneous (Sim) and sequential (Seq) presentation tasks. At the local level, each ROI shows increased phase coherence compared to baseline in the low to mid frequency range across trials following each stimulus. ITPC was calculated individually for each electrode and averaged across electrodes in a given ROI within each subject. ITPC was calculated across trials. Statistical significance was determined using cluster-based permutation tests (CBPTs). Significant time-frequency clusters are highlighted and outlined in red and blue (all p<0.003, Bonferroni corrected for 3 ROIs).

**Fig S4A:**
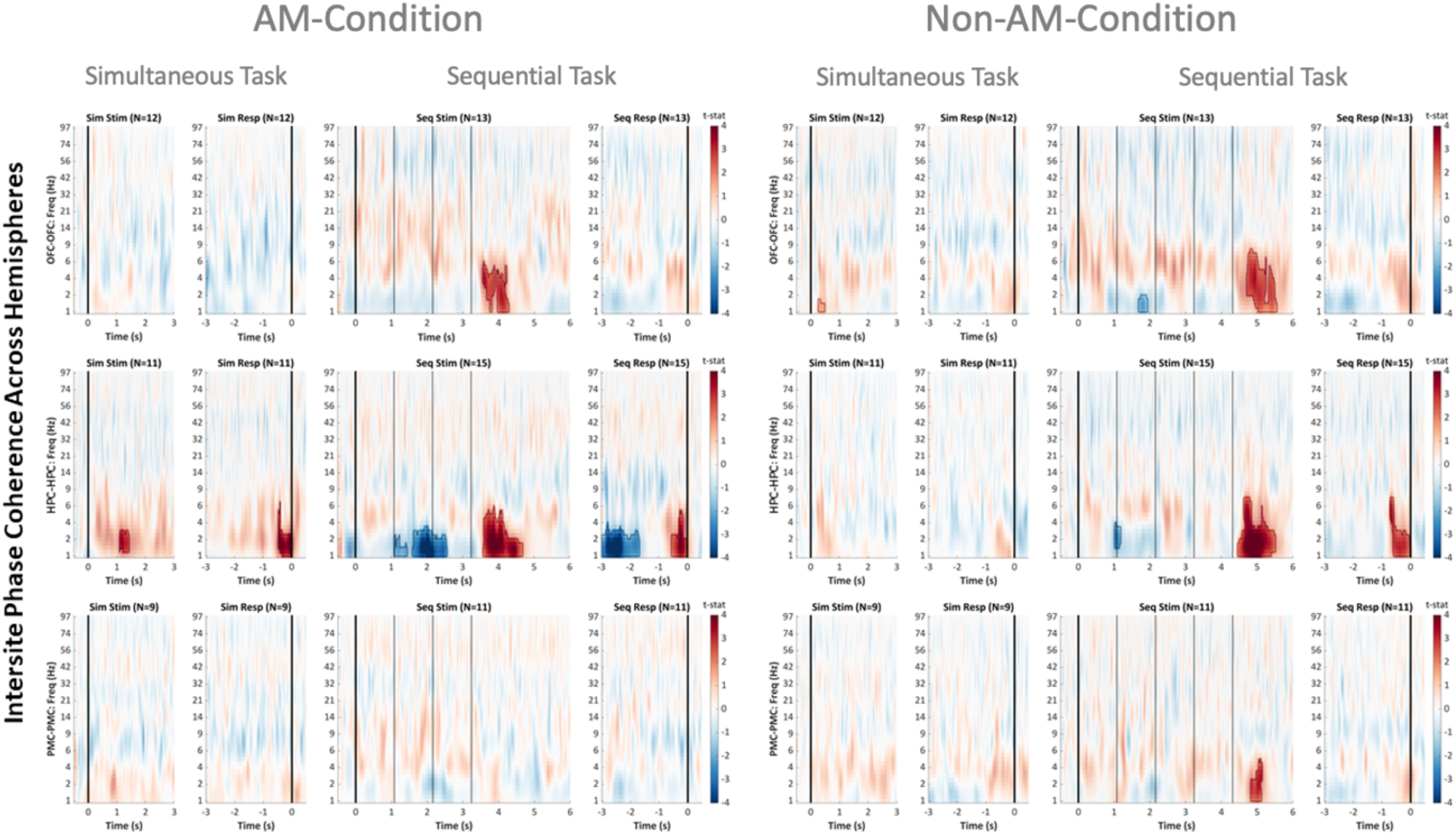
Global phase coherence across hemispheres within the same ROI. Each panel displays the group-level stimulus locked (Stim) and response locked (Resp) change in phase coherence from baseline during the simultaneous (Sim) and sequential (Seq) presentation tasks. At the global level, increased phase coherence was only observed in the LF range after the last stimulus and during the retrieval process. ISPC was calculated between different channel pairs for a given connection type and then averaged for each subject. Statistics were performed on the group level. ISPC was calculated across trials. Statistical significance was determined using cluster-based permutation tests (CBPTs). Significant time-frequency clusters are highlighted and outlined in red and blue (all p<0.002, Bonferroni corrected for 6 connection types). Note: in the Seq Task, lines correspond to the general equation “2 + 2 = 4” therefore, the increased cross-regional coherence occurs when the last number observed on the screen is compared to the calculated value stored in memory.

**Fig S4B:**
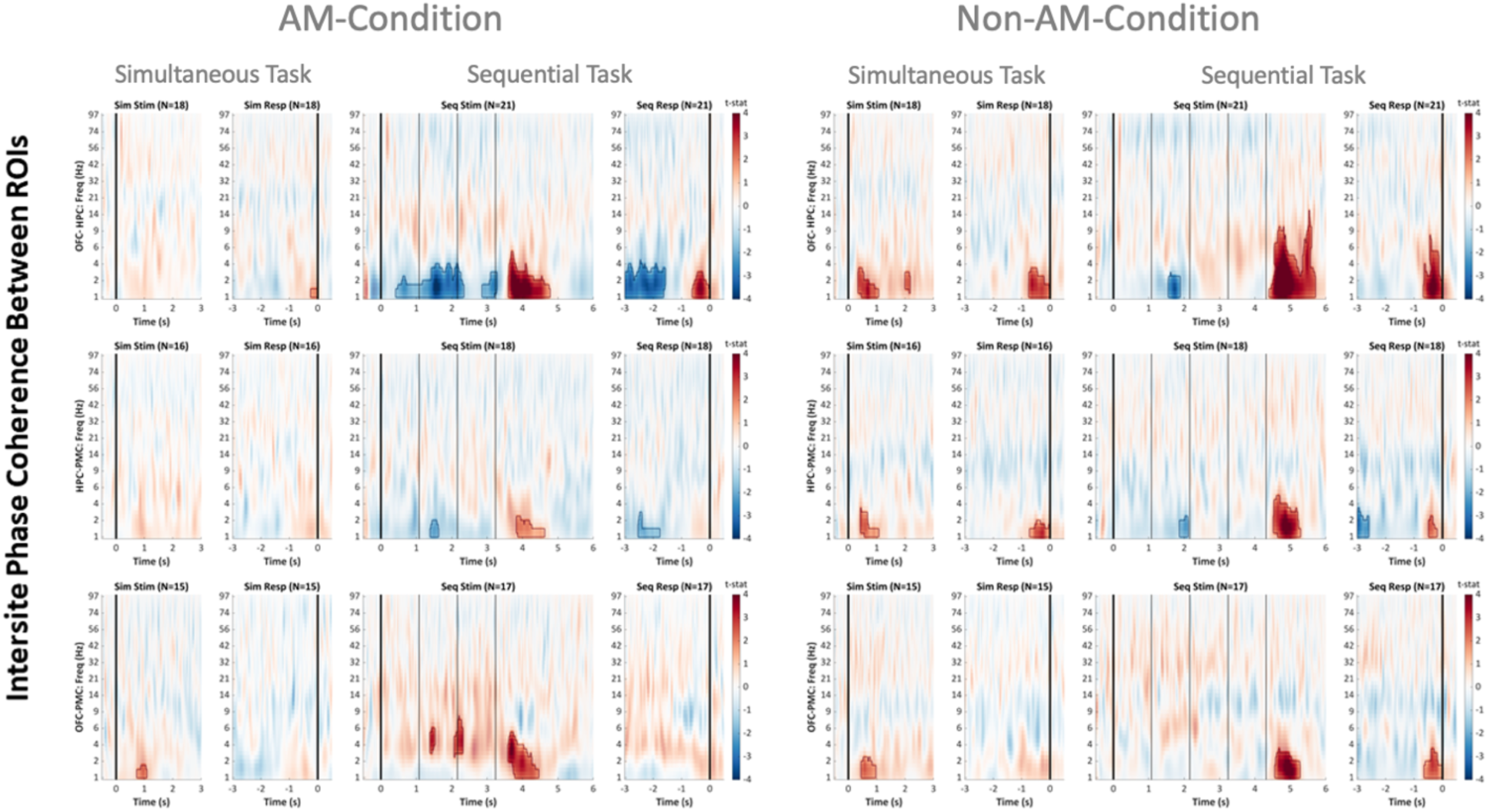
Global phase coherence across different ROIs. Each panel displays the group-level stimulus locked (Stim) and response locked (Resp) change in phase coherence from baseline during the simultaneous (Sim) and sequential (Seq) presentation tasks. At the global level, increased phase coherence was only observed in the LF range after the last stimulus and during the retrieval process. ISPC was calculated between different channel pairs for a given connection type and then averaged for each subject. Statistics were performed on the group level. ISPC was calculated across trials. Statistical significance was determined using cluster-based permutation tests (CBPTs). Significant time-frequency clusters are highlighted and outlined in red and blue (all p<0.002, Bonferroni corrected for 6 connection types). Note: in the Seq Task, lines correspond to the general equation “2 + 2 = 4” therefore, the increased cross-regional coherence occurs when the last number observed on the screen is compared to the calculated value stored in memory.

**Fig S5:**
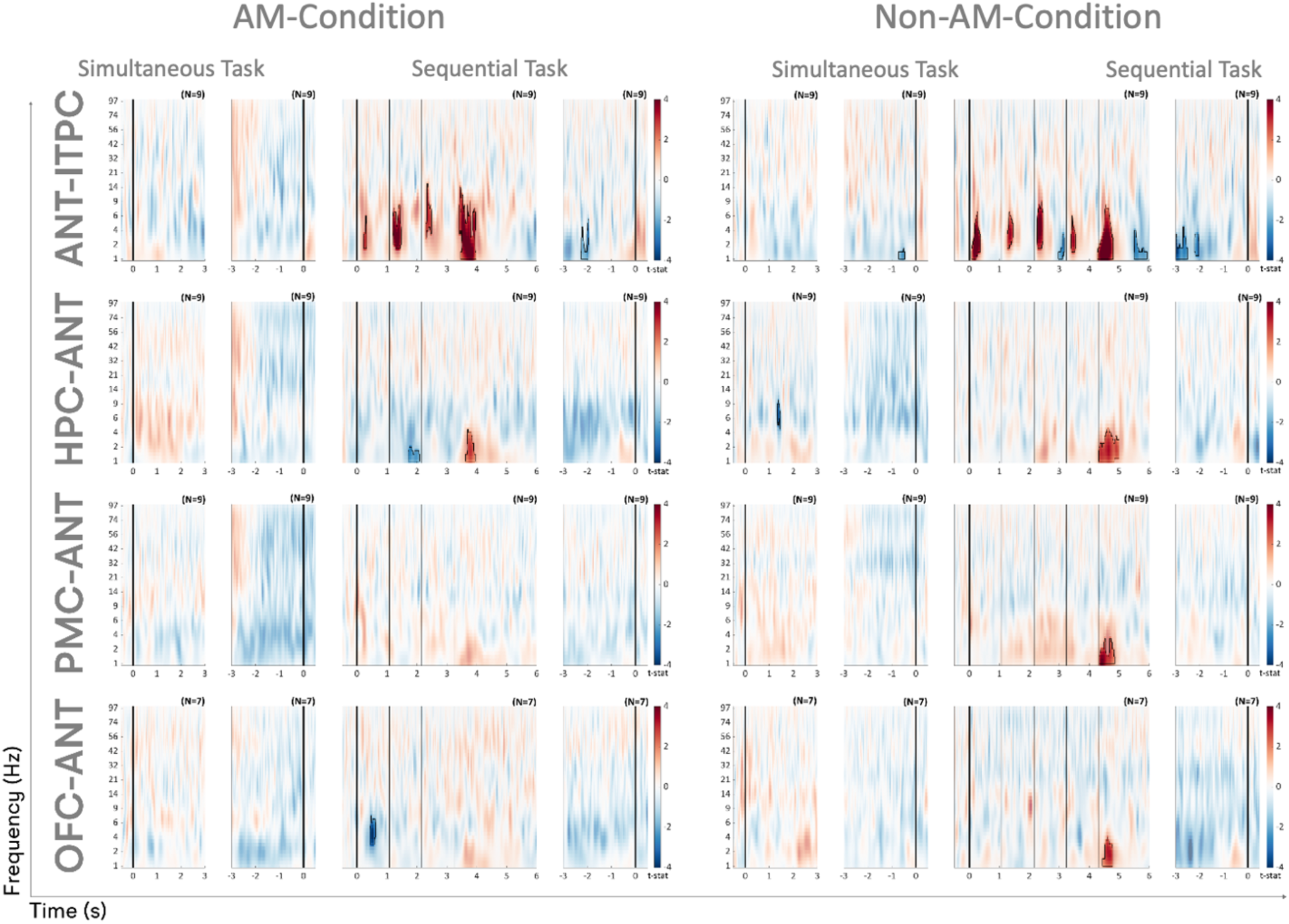
ANT displays a similar pattern of coherence as other regions. Each panel displays the group-level stimulus locked (Stim) and response locked (Resp) change in phase coherence from baseline during the simultaneous (Sim) and sequential (Seq) presentation tasks during AM (left) and Non-AM (right) conditions. Within electrodes (intertrial phase coherence; ITPC-First row), ANT also shows increased phase coherence compared to baseline in the low to mid frequency range (∼1-20Hz) across trials following each stimulus, while across electrodes of different ROIs (intersite phase coherence; ISPC), increased phase coherence was strongest in the LF range (∼1-6Hz) after the last stimulus i.e., during the retrieval process. This trend is consistent with the observations from the main text, but may not be significant between PMC/OFC and ANT in the Memory condition because of the smaller subject number. ITPC was calculated individually for each electrode and averaged across electrodes in a given ROI within each subject. ISPC was calculated between different ROIs. All channel pairs for a given connection type were averaged for each subject and statistics were performed on the group level. Both ITPC and ISPC were calculated across trials. Statistical significance was determined using cluster-based permutation tests (CBPTs). Significant time-frequency clusters are highlighted and outlined in red and blue (ITPC: all p<0.003, ISPC: all p<0.002, Bonferroni corrected for 3 connection types).

**Fig S6:**
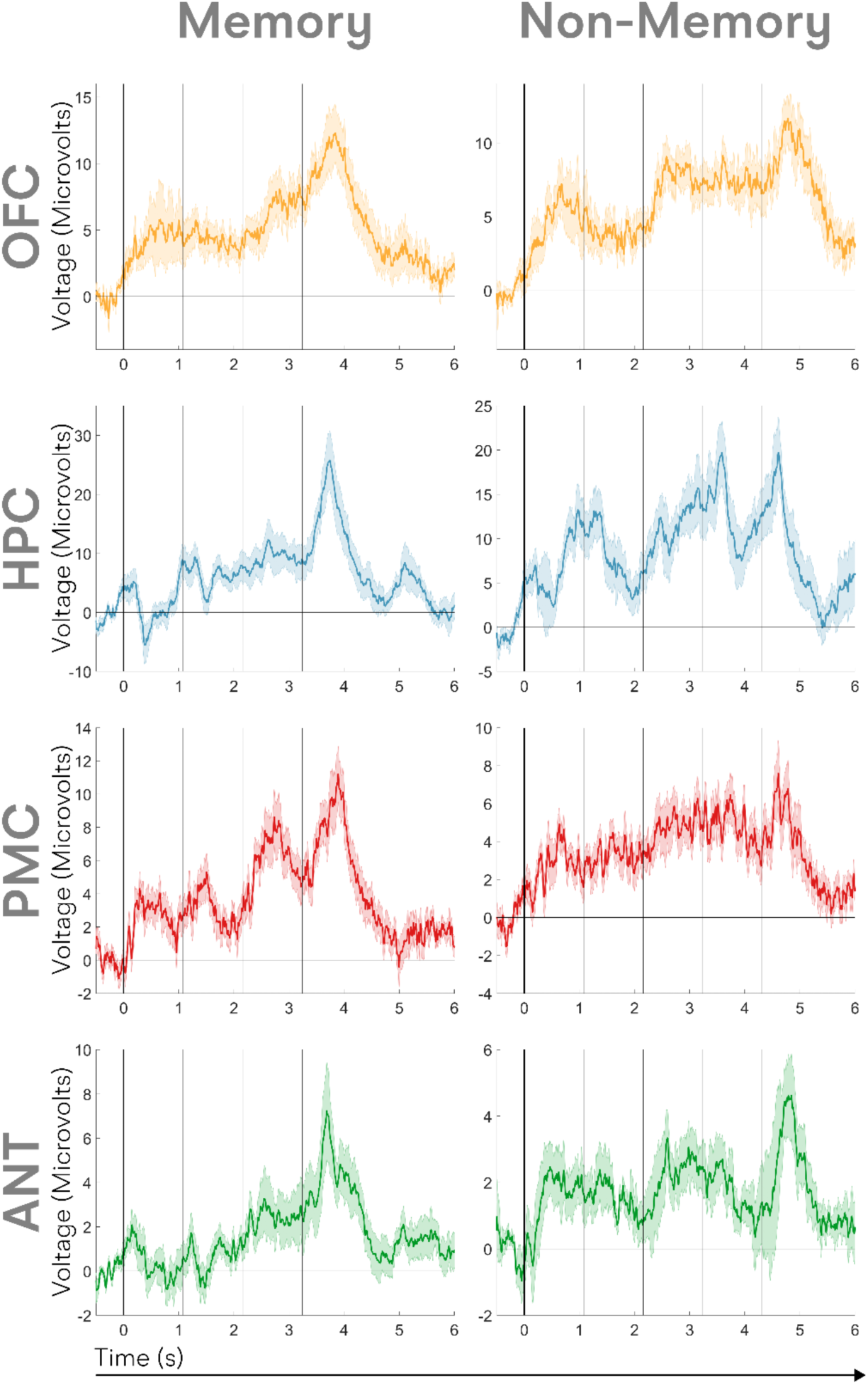
Phase coherence results cannot be completely derived from ERP phenomena. Each panel displays the group-level stimulus locked change in voltage from baseline during the sequential presentation tasks for the AM condition (memory; left) and Non-AM condition (non-memory; right). The colored shading displays the standard error of the mean. While clear increases in ITPC are seen in response to each new stimulus, the average voltage responses do not show clear peaks following every stimulus indicating this phenomenon isn’t completely an artifact of event related potentials. Further, cross regional synchrony is only observed following the final stimulus (compare **Fig S3** and **Figs S4-S6**). While there is an ERP peak in the average voltage response, this cannot be an artifact of visual responses per se because the average latency of this peak is 862ms compared to the expected average peaks of visual responses around 100ms.

**Fig S7:**
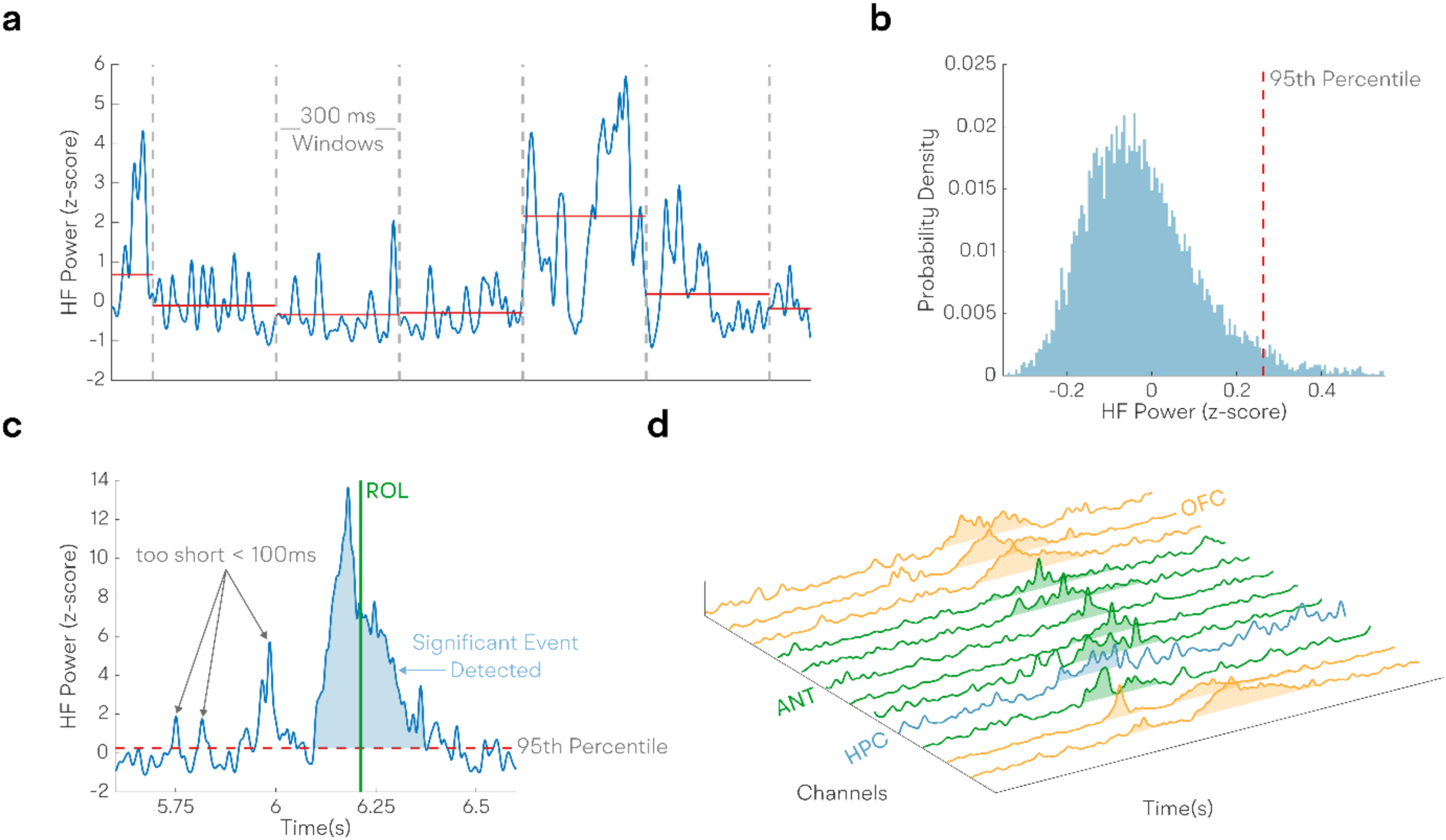
ROL Methods. This figure illustrates our method for detecting response onset latency (ROL) from neural signals.a) **Signal Smoothing and Sampling**: Neural signals, namely low-frequency (LF) power, Inter-Site Phase Coherence (ISPC), and high-frequency (HF) power, were initially smoothed by applying a 100ms moving average. Subsequently, a sampling process was employed using 10,000 random windows, each of 300ms duration, to compute the average expected signal value across these windows. b) **Permutation Distribution and Threshold Definition**: The averages obtained from the 300ms windows were utilized to define a permutation distribution, representing the expected average value across a 300ms window. A threshold was then established at the 95th percentile of this distribution. c) **Identification of Significant Events and ROL Definition**: Significant events were identified as contiguous timepoints where the signal amplitude exceeded the threshold for more than 100ms. The response onset latency (ROL) was then defined for each of these significant events. This was computed as the average time of the event, with the calculation being weighted by the signal amplitude. d) **Channel-specific ROL Identification**: For each channel, its ROL was determined as the average over all significant events for that channel that coincided with signal detection in another ROI within the same hemisphere for the given trial. In essence, for a significant event to contribute to the average ROL, it needed to be detected concurrently in at least two different ROIs within the same hemisphere and during the same trial.

**Fig S8:**
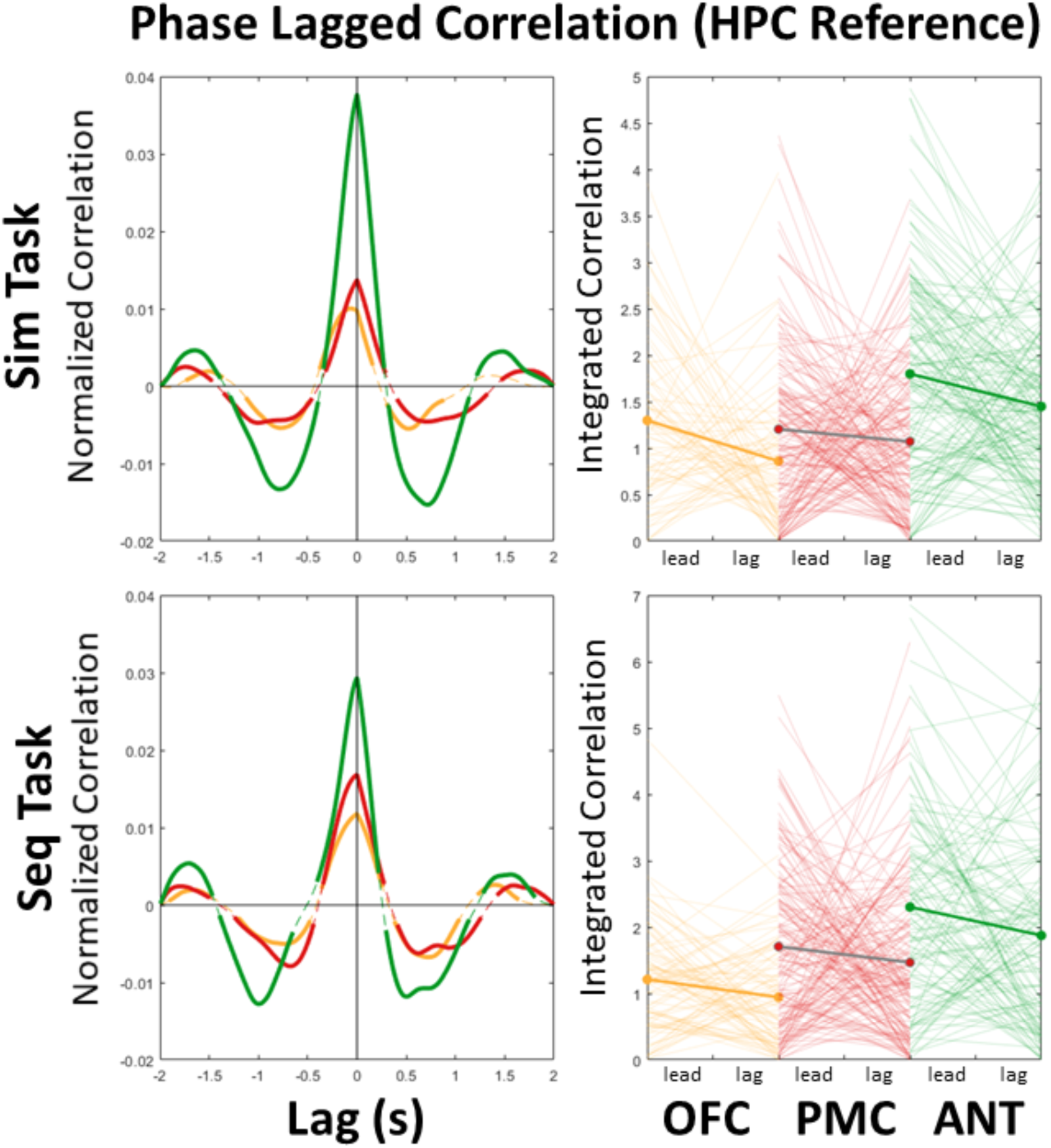
Lagged correlation of LF phases across regions suggests HPC leads LF phase. Colors are the same in the original manuscript (green = ANT; Red = PMC, and yellow = OFC) For this analysis we computed the lagged correlation between HPC LF phase as a reference and the LF phases of other ROIs during the retrieval stage. We found statistically stronger correlation values than would be expected by chance (Cluster based permutation test across electrodes; clustsum>607, p<0.0001; represented by thick lines) suggesting there is a relationship between the LF phases of these ROIs during memory retrieval phase. We then integrated the correlation values for the main positive peak in the left panels to determine whether correlation was stronger for a negative lag (left of zero; HPC LF phase leads other ROIs) or positive lag (right of zero; HPC LF phase lags other ROIs). Line plots show individual connections for the left/lead integration and the right/lag integration. We found HPC LF phase significantly leads LF phase in OFC (T(75)=2.96,p=0.004) and ANT (T(154)=3.45,p=0.0007). The same trend was observed for PMC (T(222)=1.06,p=0.29), but this was not significant. This supports our findings of HPC LF activity driving the coherence events observed in other ROIs.

**Fig S9:**
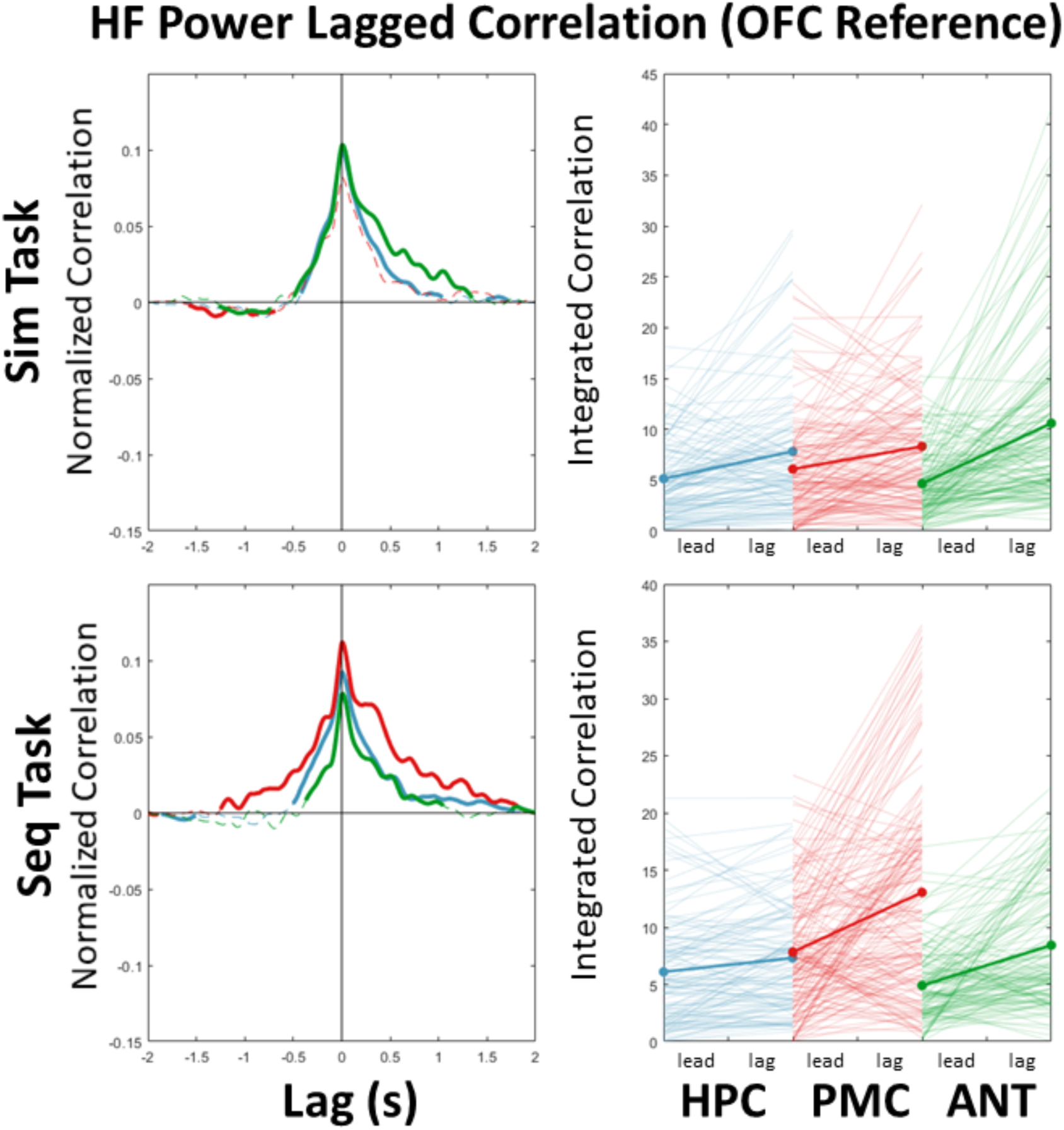
Lagged correlation of HF power across regions suggests OFC lags HF power. Colors are the same in the original manuscript (green = ANT; Red = PMC, and Blue = HPC). For this analysis we computed the lagged correlation between OFC HF power as a reference and the HF power of other ROIs. We found statistically stronger correlation values than would be expected by chance (Cluster based permutation test across electrodes; clustsum>1714, p<0.0001; represented by thick lines) suggesting there is a relationship between the HF power of these ROIs during memory retrieval stage. We then integrated the correlation values for the main positive peak in the left panels to determine whether correlation was stronger for a negative lag (left of zero; OFC HF power leads other ROIs) or positive lag (right of zero; OFC HF power lags other ROIs). Line plots show individual connections for the left/ lead integration and the right/lag integration. We found OFC HF activity significantly lags the HF power in all other ROIs (HPC: T(123)=5.95,p<0.0001, PMC: T(150)=4.98, p<0.0001,and ANT: T(153)=9.60, p<0.0001). This supports our findings of Late OFC HF power compared to other ROIs.

**Fig S10:**
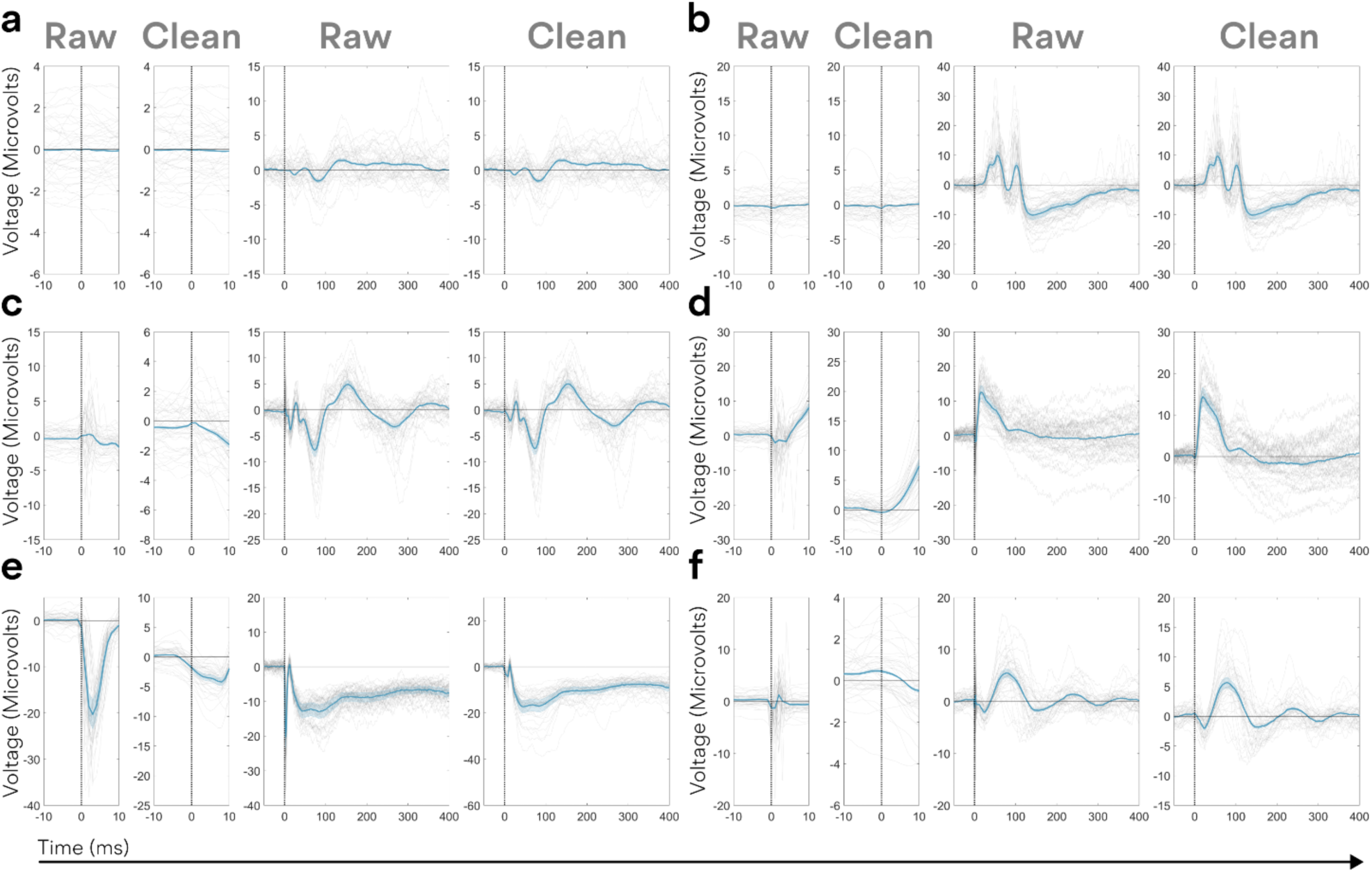
Stimulation artifact removal. In order to ensure our single pulse electrical stimulation (SPES) results were not biased by the stimulation artifact, we employed an automatic artifact rejection algorithm followed by manual artifact rejection. We first examined a 40ms window surrounding each stimulation pulse for extreme differences in voltage values. We examined the derivative of the signal for extreme values under the assumption that physiological changes in voltage would proceed more smoothly while the gradient would be much sharper for artifactual changes. If we detected derivative peaks within this window that exceeded 99% of the voltage derivative distribution, these points were removed and interpolated with an autoregressive moving average (fillgaps in MATLAB). In these panels, the thin grey lines represent data from a single trial while the blue lines represent the average and standard error across trials. The thin panels on the left show a 20ms window around the stimulation onset and the wider panels show the traditional view across hundreds of milliseconds. “Raw” panels show the voltage traces before artifact removal and “Clean” panels show the voltage trace after artifact removal. **a)**&**b)** There is generally not a significant artifact present when examining SPES across ROIs. **a)** Selected electrode PMC→OFC **b)** Selected electrode HPC→ANT **c)**&**d)** Sometimes a minor stimulation artifact is present (note the sharp triangles under the first “Raw” windows). However, the automatic artifact removal algorithm appears to remove these effects without distorting the signal. **c)** Selected electrode ANT→OFC **d)** Selected electrode ANT→PMC. **e)**&**f)** SPES traces were manually reviewed and connections showing a significant artifact still present after the automatic artifact rejection was performed were excluded from the data. **e)** Excluded electrode PMC→PMC **f)** Non-excluded electrode PMC®PMC

**Fig S11.**
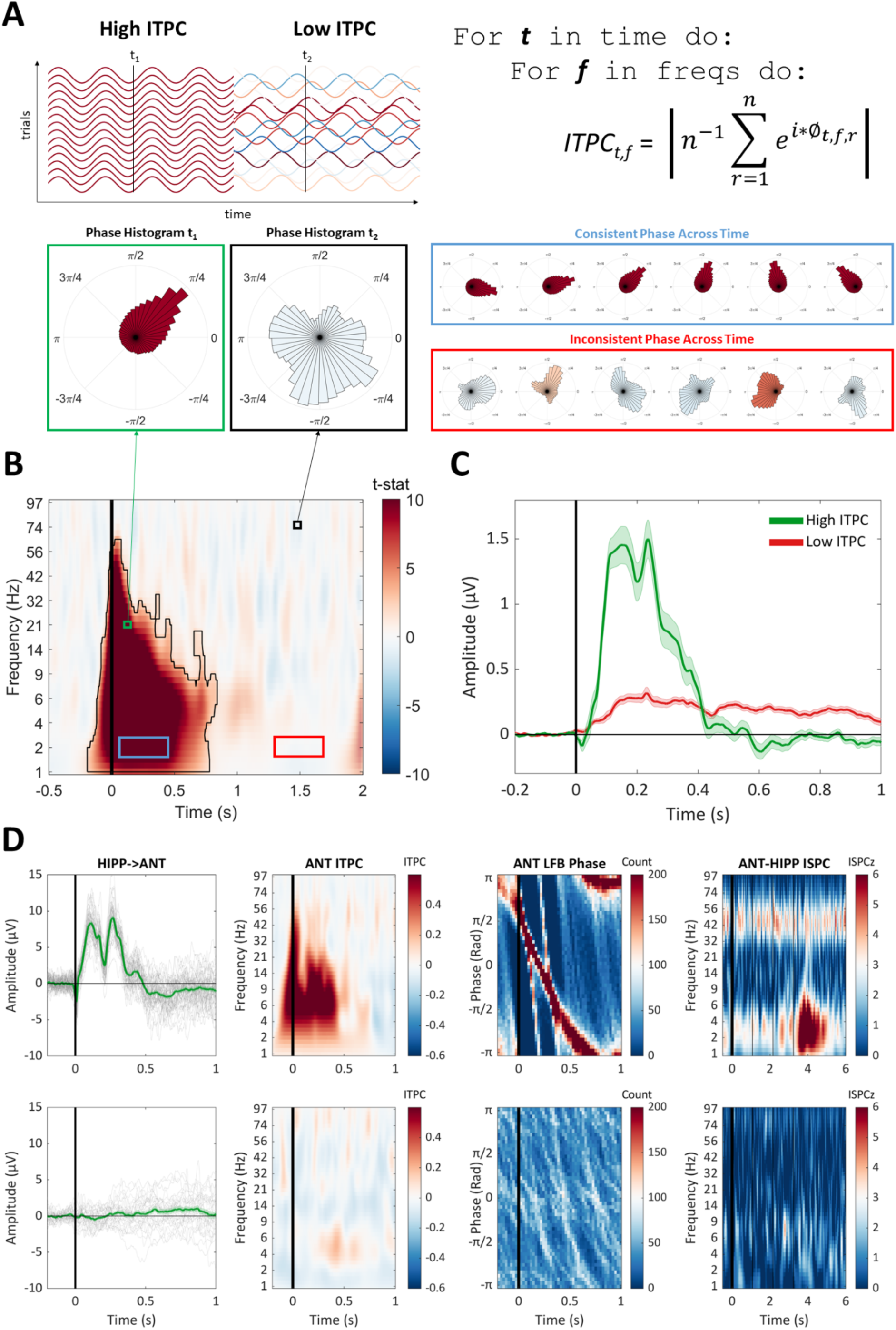
Connectivity Index: **A)** Intuition behind intertrial phase coherence (ITPC) as a measure of connectivity strength. ITPC (also known as phase-locking value) measures the consistency of the phase of a signal across trials. An estimate of the instantaneous phase (Ø) is extracted for each time point (t), frequency (f), and trialI) through wavelet decomposition. This phase is used to define a unit vector for each trial (e raised to the complex numIer i times the instantaneous phase). ITPC is calculated as the length (absolute value of a complex number) of the mean of these unit vectors across trials. When phase is consistent across trials, the length of this unit vector will be longer (ITPC closer to 1) than when the phase angles are random or uniformly distributed (ITPC closer to 0). If the voltage recorded in response to stimulation in a distal site follows the same phase trajectory every trial, the phase will continue to cyclically change, therefore we would expect large values of ITPC to propagate across time (Blue box), whereas if electrical stimulation produces a less consistent effect in the recording site (random phase throughout trials), ITPC would be lower and show less propagation through time (red box). **B)** Cluster used to calculate the connectivity index. We were interested in determining whether electrical stimulation could produce a statistically detectable change in ITPC across all stimulation-recording sites in our subjects. The ITPC values for each time-frequency point were calculated with the formula above for each stimulation-recording pair and a cluster-based permutation test was used to assess the statistical reliability across stimulation-recording pairs for the change in ITPC from baseline (baseline ITPC values were subtracted from each stimulation-recording pair and t-tests were performed at each time-frequency point with the null hypothesis that the change in ITPC from baseline would be zero). We found electrical stimulation reliably increases ITPC in distal recording sites in the time-frequency cluster shown above. ITPC was then averaged across time-frequency points in this cluster to calculate the connectivity index for each stimulation-recording pair. **C)** The connectivity index distinguishes rapid onset and consistent responses (Green) from delayed and jittered responses (Red). This plot was generated by separating then averaging the connections with the highest and lowest connectivity index. (Shaded area represents the standard error of the mean). **D)** The connectivity index is related to the consistency in the trial by trial response to stimulation and measures of functional connectivity. Two stimulation-recording pairs are shown here one for a large connectivity index (upper panels) and one for a low connectivity index (lower panels). Left to right are shown the voltage recorded in ANT in response to HIPP (HPC) stimulation, ANT ITPC calculated from these trials, the histogram of phase values in the LF band across trials, and the inter site phase coherence (ISPC) between these two electrodes during the AM condition of the Seq task. The high connectivity index stimulation-recording pair displayed a highly consistent voltage and phase profile that persisted across time, leading to larger values of ITPC across the cluster identified in **B.** As described in the main text, higher values of connectivity index between HIPP and ANT predicted stronger ISPC between these sites during memory tasks. A similar procedure was used to calculate ISPC between sites in response to stimulation. In this case, the difference in phase values between two sites replaces the instantaneous phase (Ø) in the equation above.

**Fig S12:**
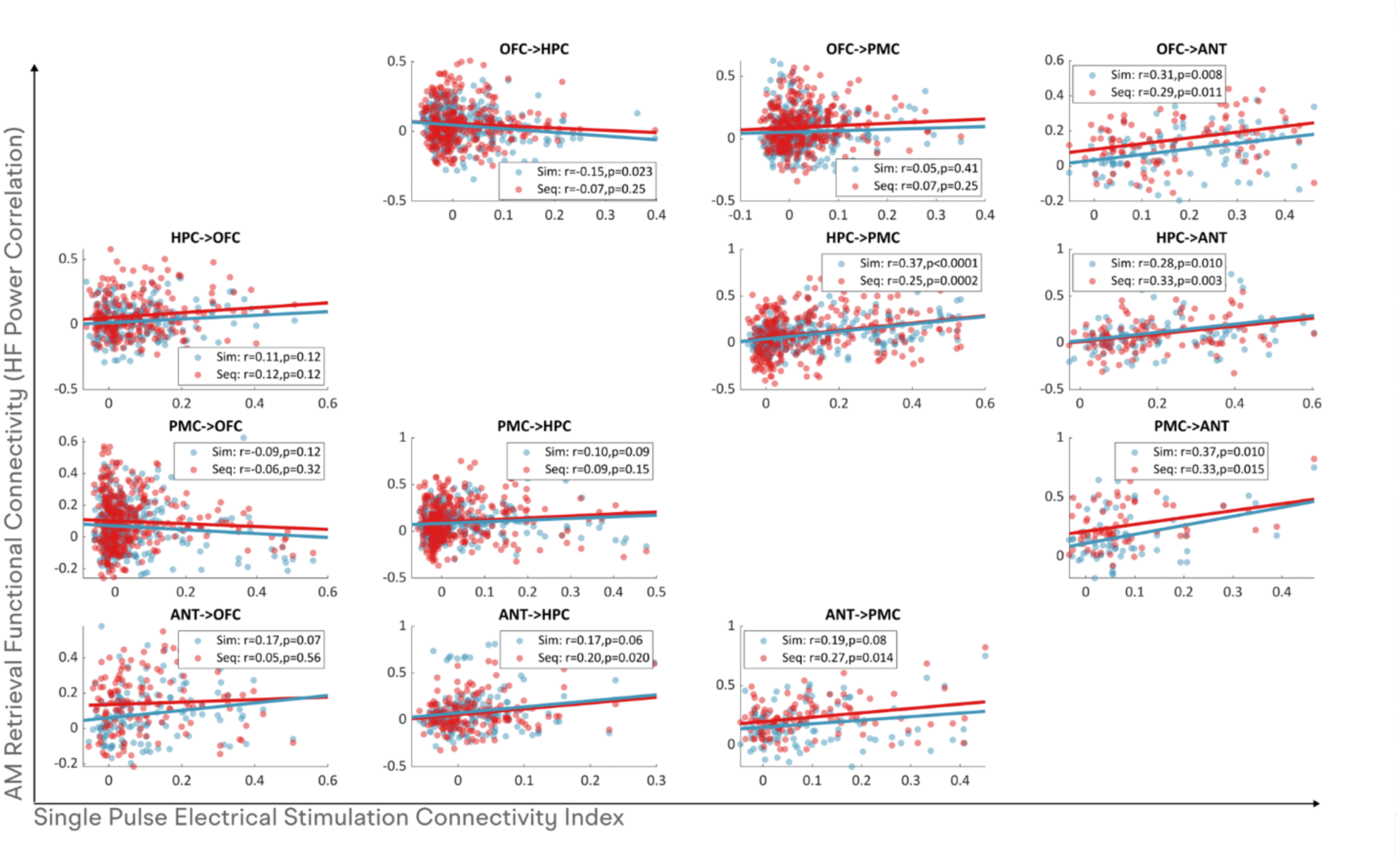
Correlations between connectivity index and HF power correlation. This figure shows the pairwise correlations between connectivity index and trial by trial HF power across a pair of ROIs. All p-values were corrected for multiple comparisons using the Bonferroni-Hochberg procedure. Note: Clumping around zero connectivity index occurs when many recording-stimulation pairs are present, but few of these stimulation sites result in significant voltage deflections in the distant recording sites.

**Fig S13:**
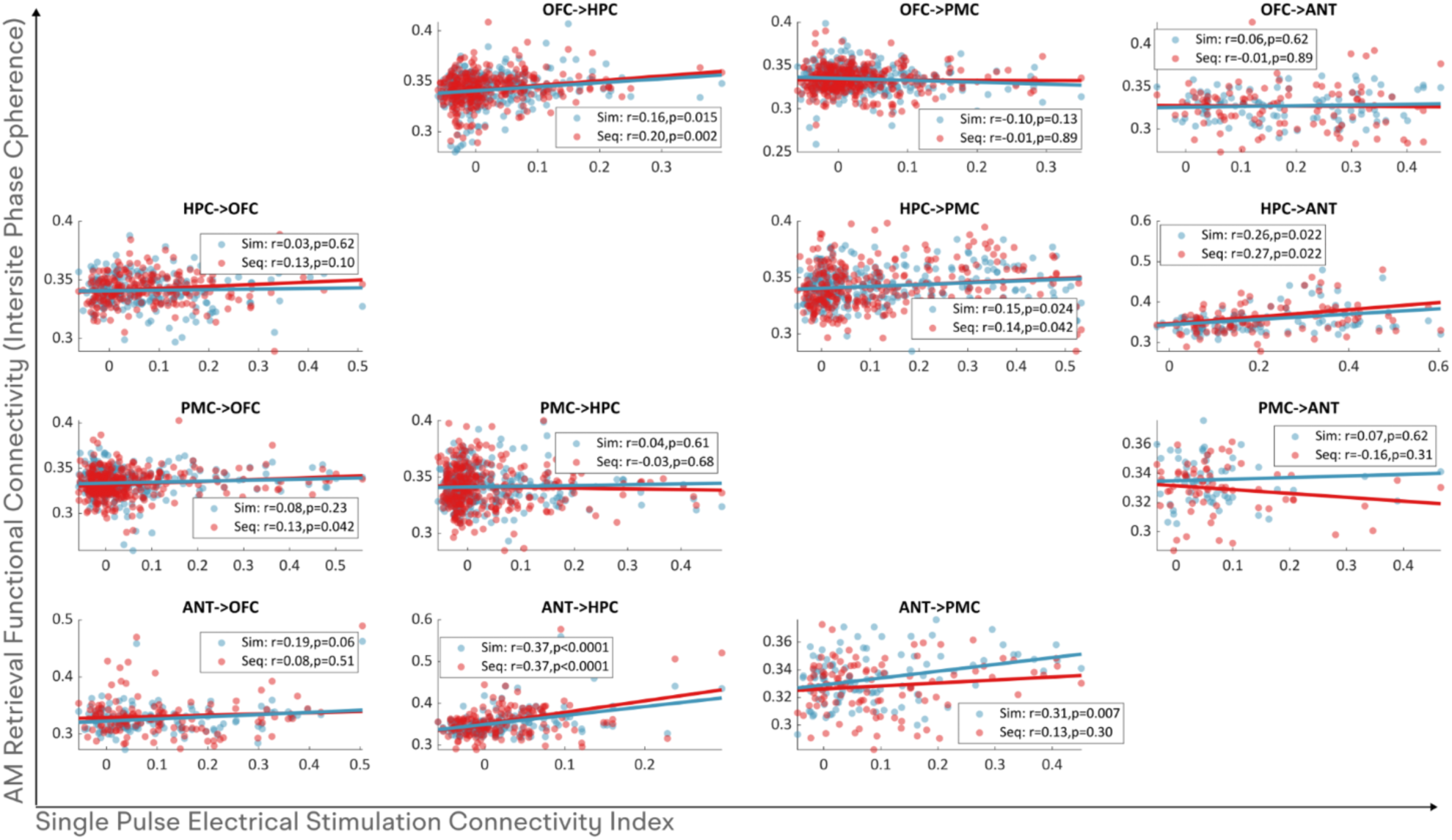
Correlations between connectivity index and LF Intersite phase coherence (ISPC). This figure shows the pairwise correlations between connectivity index and LF ISPC across a pair of ROIs. All p-values were corrected for multiple comparisons using the Bonferroni-Hochberg procedure. Note: Clumping around zero connectivity index occurs when many recording-stimulation pairs are present, but few of these stimulation sites result in significant voltage deflections in the distant recording sites.

**Fig S14:**
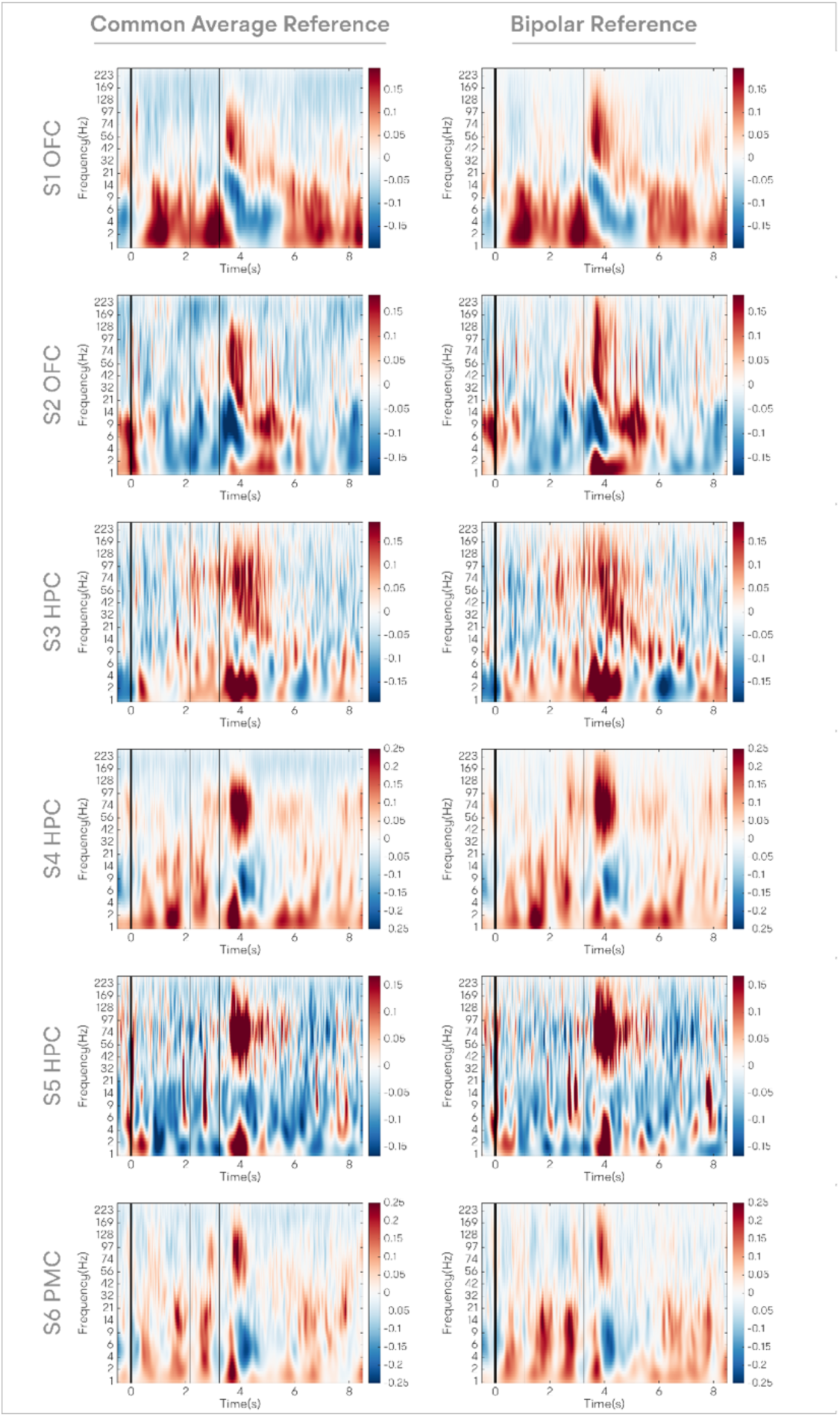
Common Average vs Bipolar References. This figure compares the results of using two different referencing methods, namely common average referencing and bipolar referencing, in analyzing neural signals. Both methods yield qualitatively similar results, demonstrating the robustness of the measurements across the two referencing techniques. **Left Panels**: Display spectrograms averaged over all channels of a given Region of Interest (ROI) for a single subject using common average referencing. **Right Panels**: Present spectrograms averaged over all channels of a given ROI for the same subject but using bipolar referencing. The color coding on the spectrograms corresponds to variations in z-scored power from the baseline. Warm colors indicate an increase in z-scored power from the baseline, while cool colors signify decreases in z-scored power from the baseline. Taken together, these side-by-side comparisons demonstrate the overall consistency of the results regardless of the referencing method employed.

**Table S1:** Patient Demographics. This table presents the demographic and clinical details for the patients included in our study. Note: relatively few patients had seizure onset zones outside the medial temporal lobe. N/A for IQ when information was not available. Patients **S18-S31** represent the thalamic cohort in our study.

| Subject ID | Data type | Seizure Onset Zone | Seizure Hemisphere | Gender | Age | Handedness |
| --- | --- | --- | --- | --- | --- | --- |
| S1 | sEEG | Insula | right | F | 21 | R |
| S2 | sEEG | Temporal lobe | left | M | 30 | R |
| S3 | sEEG | Temporal lobe | left | F | 30 | R |
| S4 | sEEG | Temporal lobe | left | M | 30 | R |
| S5 | sEEG | Insula | left | F | 44 | Both |
| S6 | sEEG | Temporal lobe | left | M | 30 | R |
| S7 | sEEG | Temporal lobe | right | F | 23 | R |
| S8 | sEEG | Temporoparietal Junction | left | F | 19 | R |
| S9 | sEEG | Temporal lobe | left | F | 32 | R |
| S10 | sEEG | Temporal lobe | right | F | 35 | R |
| S11 | sEEG | Frontal Lobe | right | F | 41 | L |
| S12 | sEEG | Temporal lobe | left | M | 49 | R |
| S13 | sEEG | Parietal lobe | both | M | 26 | R |
| S14 | sEEG | Temporal lobe | left | M | 35 | R |
| S15 | sEEG | Parietal lobe | right | F | 45 | R |
| S16 | sEEG | Unknown | N/A | F | 29 | R |
| S17 | sEEG | Insula | both | M | 46 | R |
| S18 | sEEG | Temporal lobe | right | F | 37 | R |
| S19 | sEEG | Insula | right | F | 23 | L |
| S20 | sEEG | Occipital lobe | right | M | 23 | R |
| S21 | sEEG | Temporal lobe | left | M | 40 | R |
| S22 | sEEG | Temporal lobe | both | M | 52 | R |
| S23 | sEEG | Temporal lobe | left | M | 48 | R |
| S24 | sEEG | Temporal lobe | left | F | 34 | R |
| S25 | sEEG | Temporal lobe | both | M | 19 | R |
| S26 | sEEG | Temporal lobe | both | M | 28 | R |
| S27 | sEEG | Occipital lobe | right | F | 23 | R |
| S28 | sEEG | Temporal lobe | both | F | 37 | R |
| S29 | sEEG | Temporal lobe | right | M | 33 | R |
| S30 | sEEG | Temporal lobe | left | F | 36 | R |
| S31 | sEEG | Temporal lobe | right | M | 31 | R |

**Table S2:** Patient Stimuli and task Performance. Trial numbers and reaction times (RTs) for each trial category are listed with their mean and standard deviation (SD) for the Sim and Seq tasks. Significant differences in RTs are listed in subsequent columns. Data were compared with T-tests and corrected for multiple comparisons with Bonferroni-Hochberg correction.

| Task | Trial Category | Trial Type | Trial Number (mean±SD) | Reaction Time (s) (mean±SD) | Longer than (ms) (mean±SD) | T(DF)=X | p | Significance |
| --- | --- | --- | --- | --- | --- | --- | --- | --- |
| Sim Task | Total Trials | AllTrials | 268±123 | 3.14±0.97 |  |  |  |  |
| Sim Task | MathVsMemory | Autobio | 97±43 | 2.62±0.89 |  |  |  |  |
| Sim Task | MathVsMemory | Math | 97±43 | 3.66±1.23 | Autobio: +1033±930 | (18)=4.84 | 0.0001 | **** |
| Sim Task | MemoryAge | Recent | 48±21 | 2.56±0.88 |  |  |  |  |
| Sim Task | MemoryAge | Remote | 48±21 | 2.68±0.94 |  |  |  |  |
| Sim Task | MemoryAge | Fact | 48±21 | 3.66±1.71 | Recent: +958±874, Remote: +771±977 | (12)=3.95, (12)=2.84 | 0.002, 0.015 | **, * |
| Sim Task | MemoryResponse | True | 39±20 | 2.77±0.96 | False: +213±366 | (18)=2.54 | 0.02 | * |
| Sim Task | MemoryResponse | False | 57±26 | 2.55±0.88 |  |  |  |  |
| Sim Task | MemoryContent | Pleasant | 29±13 | 2.50±0.90 |  |  |  |  |
| Sim Task | MemoryContent | Unpleasant | 8±4 | 2.74±1.13 |  |  |  |  |
| Sim Task | MemoryContent | Neutral | 59±26 | 2.67±0.88 | Pleasant: +265±208 | (18)=3.44 | 0.0029 | ** |
| Sim Task | VerbStructure | Active | 65±29 | 2.57±0.87 |  |  |  |  |
| Sim Task | VerbStructure | Generic | 31±14 | 2.74±0.96 | Active: +165±290 | (18)=2.49 | 0.02 | * |
| Seq Task | Total Trials | AllTrials | 220±81 | 1.55±0.48 |  |  |  |  |
| Seq Task | MathVsMemory | Autobio | 110±40 | 1.65±0.47 | Math: +205±362 | (21)=2.66 | 0.01 | * |
| Seq Task | MathVsMemory | Math | 110±40 | 1.45±0.56 |  |  |  |  |
| Seq Task | MemoryAge | Recent | 55±20 | 1.62±0.45 |  |  |  |  |
| Seq Task | MemoryAge | Remote | 55±20 | 1.69±0.50 | Recent: +68±147 | (21)=2.66 | 0.04 | * |
| Seq Task | MemoryAge | Fact | NaN | NaN | NaN | NaN | NaN |  |
| Seq Task | MemoryResponse | True | 41±19 | 1.75±0.60 | False: +135±302 | (21)=2.09 | 0.049 | * |
| Seq Task | MemoryResponse | False | 68±24 | 1.62±0.44 |  |  |  |  |
| Seq Task | MemoryContent | Pleasant | 41±15 | 1.62±0.47 |  |  |  |  |
| Seq Task | MemoryContent | Unpleasant | 17±6 | 1.83±0.58 | Pleasant: +208±222, Neutral: +211±172 | (21)=4.39, (21)=5.76 | 0.0003, <0.0001 | ***, **** |
| Seq Task | MemoryContent | Neutral | 53±20 | 1.62±0.46 |  |  |  |  |
| Seq Task | VerbStructure | Active | 50±18 | 1.57±0.49 |  |  |  |  |
| Seq Task | VerbStructure | Generic | 60±22 | 1.73±0.47 | Active: +162±130 | (21)=5.86 | <0.0001 | **** |

**Table S3:**
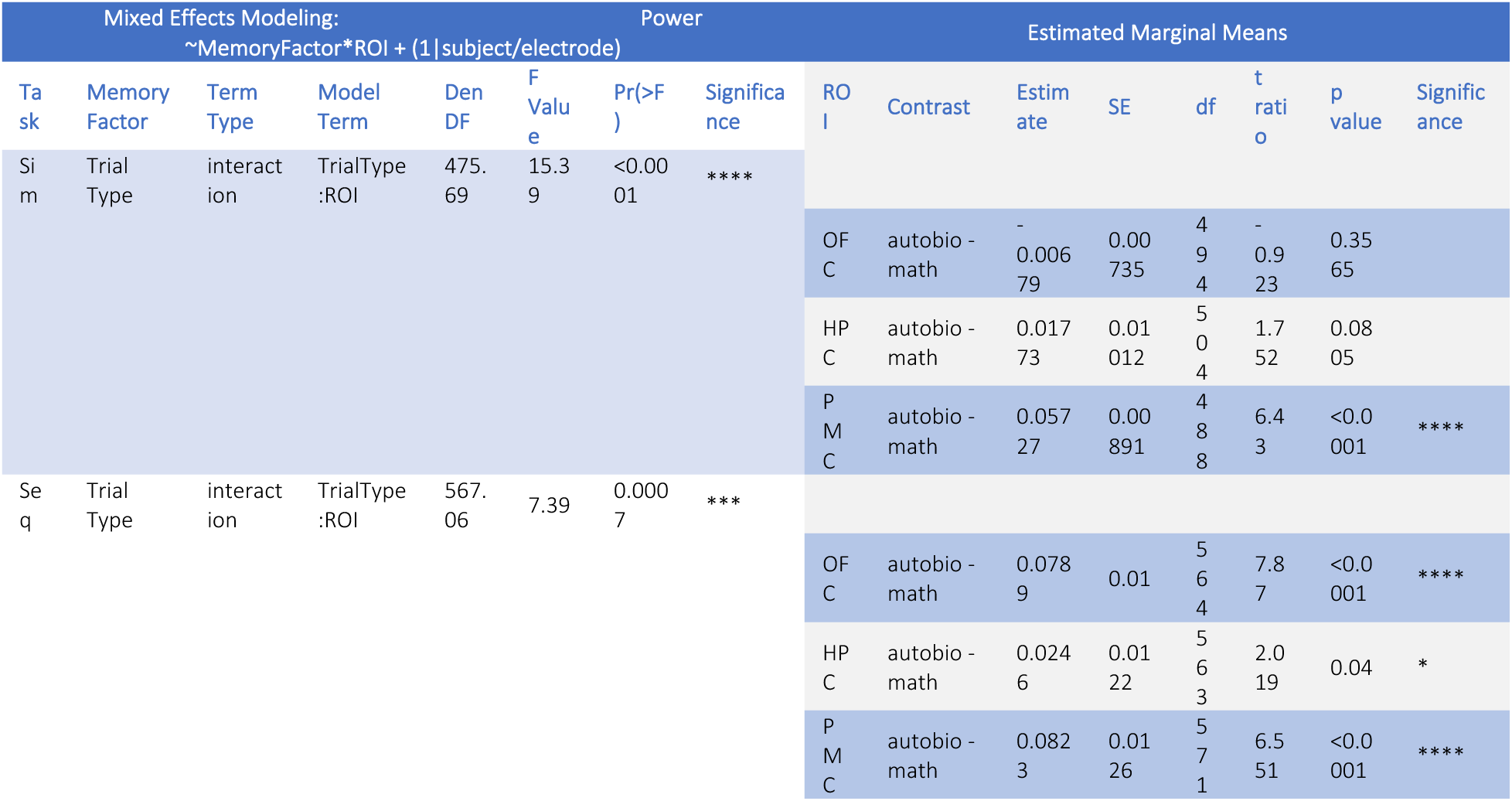
Mixed-Effects modeling of LF Power. This table displays the significant interactions observed between ROI and trial type when modeling LF power. The results suggest, on average, more LF power is present during autobiographical memory trials (autobio/AM) than during math, especially in the Seq task.

| Mixed Effects Modeling:<br>~MemoryFactor*ROI + (1\|subject/electrode) |  |  |  |  |  |  | Power | Estimated Marginal Means |  |  |  |  |  |  |  |
| --- | --- | --- | --- | --- | --- | --- | --- | --- | --- | --- | --- | --- | --- | --- | --- |
| Ta sk | Memory Factor | Term Type | Model Term | Den DF | F Value | Pr(>F ) | Significa nce | ROI | Contrast | Estimate | SE | df | t ratio | p value | Significance |
| Sim | Trial Type | interaction | TrialType :ROI | 475.69 | 15.39 | <0.0001 | **** |  |  |  |  |  |  |  |  |
|  |  |  |  |  |  |  |  | OF C | autobio - math | -0.00679 | 0.00735 | 494 | -0.923 | 0.3565 |  |
|  |  |  |  |  |  |  |  | HP C | autobio - math | 0.01773 | 0.01012 | 504 | 1.752 | 0.0805 |  |
|  |  |  |  |  |  |  |  | P M C | autobio - math | 0.05727 | 0.00891 | 488 | 6.43 | <0.0001 | **** |
| Seq | Trial Type | interaction | TrialType :ROI | 567.06 | 7.39 | 0.0007 | *** |  |  |  |  |  |  |  |  |
|  |  |  |  |  |  |  |  | OF C | autobio - math | 0.0789 | 0.01 | 564 | 7.87 | <0.0001 | **** |
|  |  |  |  |  |  |  |  | HP C | autobio - math | 0.0246 | 0.0122 | 563 | 2.019 | 0.04 | * |
|  |  |  |  |  |  |  |  | P M C | autobio - math | 0.0823 | 0.0126 | 571 | 6.551 | <0.0001 | **** |

**Table S4:**
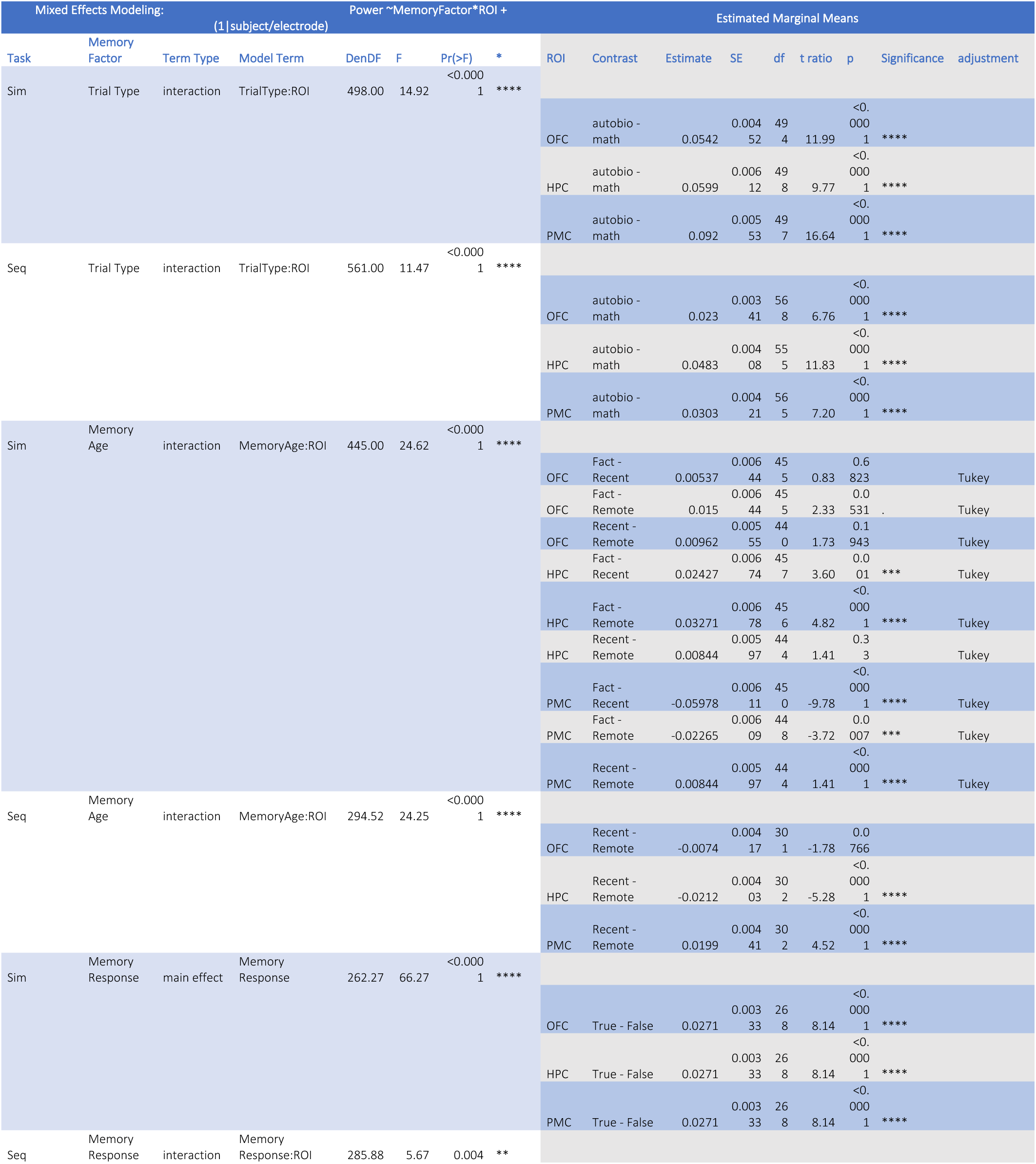
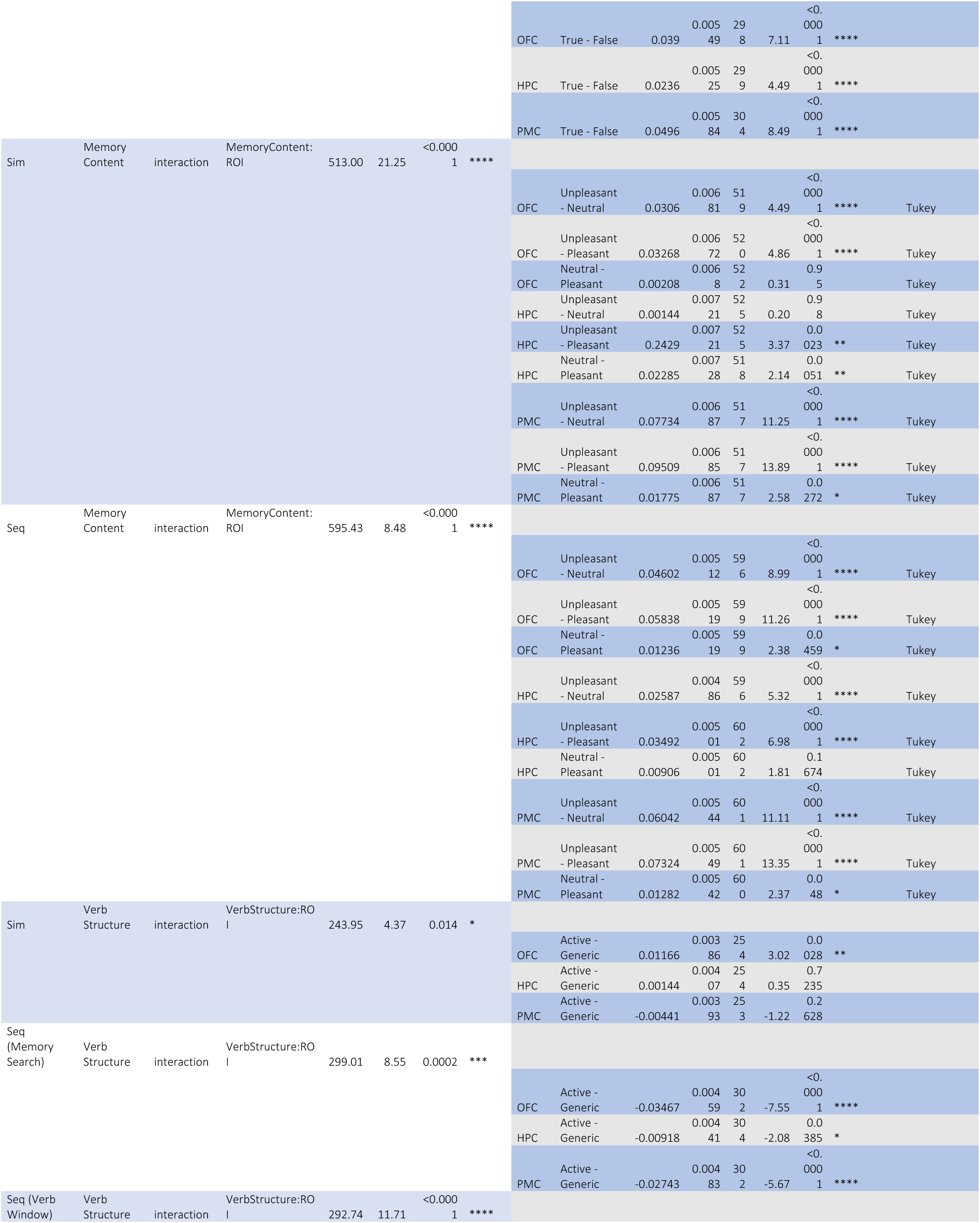
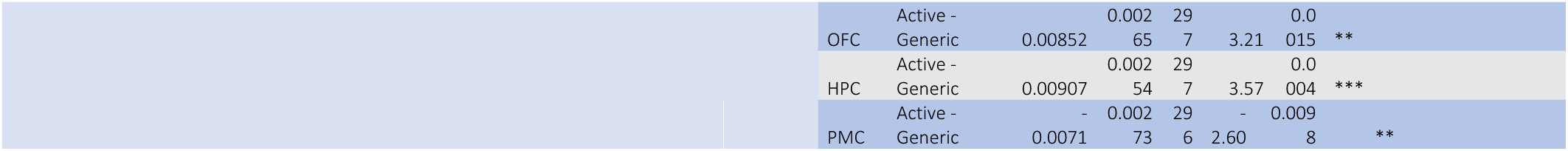
Mixed-Effects modeling of HF Power. This table displays the significant interactions observed between ROI and trial type when modeling HF power. Many significant interactions were observed suggesting the semantic content of the AM statement can causally influence the HF power in each ROI differentially. Note-Verb structure was calculated for two different windows: 1. Memory search is the standard window used during the Seq Task (0.5-1.5s following the last stimulus), 2. Verb window was calculated during the 1s following the verb presentation in the seq task.

**Table S5:**
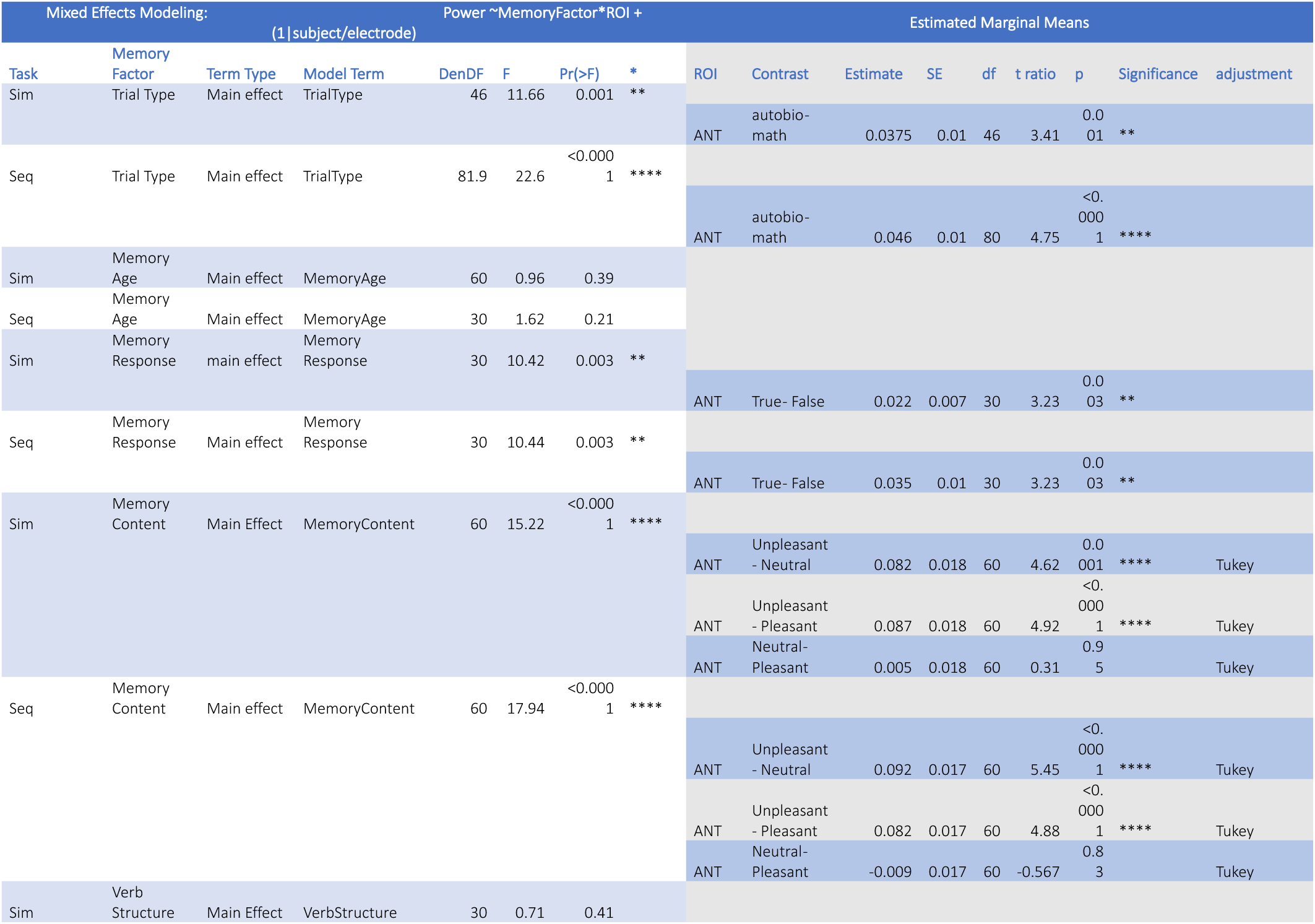
Mixed-Effects modeling of HF Power in ANT electrodes. This table displays the significant main effects of trial type when modeling HF power in ANT contacts. Many significant main effects were observed suggesting the semantic content of the AM statement can causally influence the HF power in ANT contacts.

**Table S6:** Mixed-effects modeling of electrophysiological event timing for the sim task. This table displays the significant interactions observed between ROI and feature when modeling the response onset latency (ROL) of different electrophysiological events during the sim task. A significant interaction was observed suggesting the timing of electrophysiological features differs between ROIs. Note: ISPC was assumed to have the same timing across connection types (See Table S8).

| Mixed Effects Modeling: Response Onset Latency ~ROI*Feature + (1\|subject) |  |  |  |  |  | Estimated Marginal Means |  |  |  |  |  |  |  |
| --- | --- | --- | --- | --- | --- | --- | --- | --- | --- | --- | --- | --- | --- |
| Task | Model Term | DenDF | F Value | Pr(>F) | Significance | Contrast | Estimate (ms) | SE | df | t ratio | p value | Significance | adjustment |
| Sim | ROI*Feature | 961.33 | 12.51 | <0.0001 | **** | HF - ISPC \| OFC | 442.7 | 33 | 962 | 13.44 | <0.0001 | **** | Tukey |
|  |  |  |  |  |  | HF - LF \| OFC | 494.3 | 31.7 | 961 | 15.59 | <0.0001 | **** | Tukey |
|  |  |  |  |  |  | ISPC - LF \| OFC | 51.5 | 33 | 962 | 1.56 | 0.26 |  | Tukey |
|  |  |  |  |  |  | HF - ISPC \| HPC | 314.5 | 32.1 | 962 | 9.79 | <0.0001 | **** | Tukey |
|  |  |  |  |  |  | HF - LF \| HPC | 290.5 | 32.1 | 961 | 9.03 | <0.0001 | **** | Tukey |
|  |  |  |  |  |  | ISPC - LF \| HPC | -24 | 32.1 | 962 | -0.75 | 0.74 |  | Tukey |
|  |  |  |  |  |  | HF - ISPC \| PMC | 302.7 | 33 | 961 | 9.186 | <0.0001 | **** | Tukey |
|  |  |  |  |  |  | HF - LF \| PMC | 327.7 | 33.7 | 961 | 9.724 | <0.0001 | **** | Tukey |
|  |  |  |  |  |  | ISPC - LF \| PMC | 24.9 | 33.1 | 961 | 0.75 | 0.73 |  | Tukey |
|  |  |  |  |  |  | HF - ISPC \| ANT | 111.9 | 58.5 | 961 | 1.91 | 0.14 |  | Tukey |
|  |  |  |  |  |  | HF - LF \| ANT | -52.8 | 57.4 | 961 | -0.92 | 0.63 |  | Tukey |
|  |  |  |  |  |  | ISPC - LF \| ANT | -164.6 | 58.5 | 961 | -2.81 | 0.01 | * | Tukey |
|  |  |  |  |  |  | HPC - OFC \| HF | -109.7 | 33.9 | 964 | -3.236 | 0.007 | ** | Tukey |
|  |  |  |  |  |  | HPC - PMC \| HF | 44.8 | 34.1 | 963 | 1.311 | 0.56 |  | Tukey |
|  |  |  |  |  |  | HPC - ANT \| HF | 285.5 | 47.8 | 962 | 5.971 | <0.0001 | **** | Tukey |
|  |  |  |  |  |  | OFC - PMC \| HF | 154.5 | 34.1 | 963 | 4.525 | <0.0001 | **** | Tukey |
|  |  |  |  |  |  | OFC - ANT \| HF | 395.2 | 48.1 | 963 | 8.225 | <0.0001 | **** | Tukey |
|  |  |  |  |  |  | PMC - ANT \| HF | 240.7 | 48.8 | 963 | 4.93 | <0.0001 | **** | Tukey |
|  |  |  |  |  |  | HPC - OFC \| ISPC | 18.5 | 34.3 | 963 | 0.54 | 0.95 |  | Tukey |
|  |  |  |  |  |  | HPC - PMC \| ISPC | 33 | 33.2 | 962 | 0.995 | 0.75 |  | Tukey |
|  |  |  |  |  |  | HPC - ANT \| ISPC | 82.9 | 48.9 | 962 | 1.696 | 0.32 |  | Tukey |
|  |  |  |  |  |  | OFC - PMC \| ISPC | 14.5 | 34.3 | 962 | 0.422 | 0.97 |  | Tukey |
|  |  |  |  |  |  | OFC - ANT \| ISPC | 64.4 | 49.4 | 962 | 1.3 | 0.56 |  | Tukey |
|  |  |  |  |  |  | PMC - ANT \| ISPC | 49.9 | 49.6 | 962 | 1.01 | 0.75 |  | Tukey |
|  |  |  |  |  |  | HPC - OFC \| LF | 94 | 33.9 | 964 | 2.773 | 0.029 | * | Tukey |
|  |  |  |  |  |  | HPC - PMC \| LF | 82 | 34.4 | 963 | 2.382 | 0.081 |  | Tukey |
|  |  |  |  |  |  | HPC - ANT \| LF | -57.7 | 47.8 | 962 | -1.21 | 0.622 |  | Tukey |
|  |  |  |  |  |  | OFC - PMC \| LF | -12.1 | 34.3 | 963 | -0.35 | 0.99 |  | Tukey |
|  |  |  |  |  |  | OFC - ANT \| LF | -151.8 | 48 | 963 | -3.162 | 0.009 | ** | Tukey |
|  |  |  |  |  |  | PMC - ANT \| LF | -139.7 | 49 | 963 | -2.85 | 0.023 |  | Tukey |

**Table S7:**
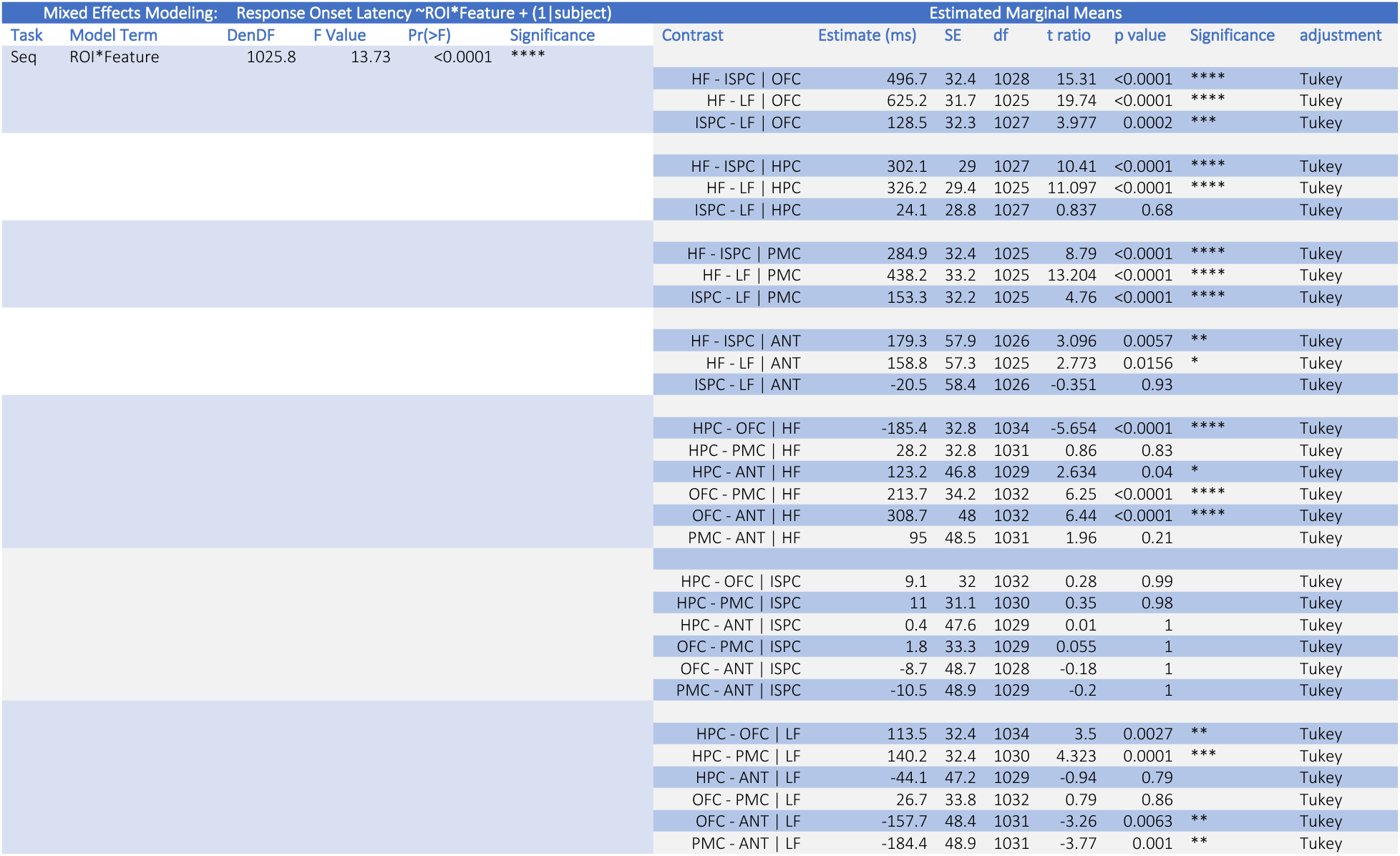
Mixed-effects modeling of electrophysiological event timing for the seq task. This table displays the significant interactions observed between ROI and feature when modeling the response onset latency (ROL) of different electrophysiological events during the seq task. A significant interaction was observed suggesting the timing of electrophysiological features differs between ROIs. OFC and PMC display significant differences in the timing of electrophysiological features, while the difference in HPC is less pronounced. Note: ISPC was assumed to have the same timing across connection types (See Table S8).

**Table S8:**
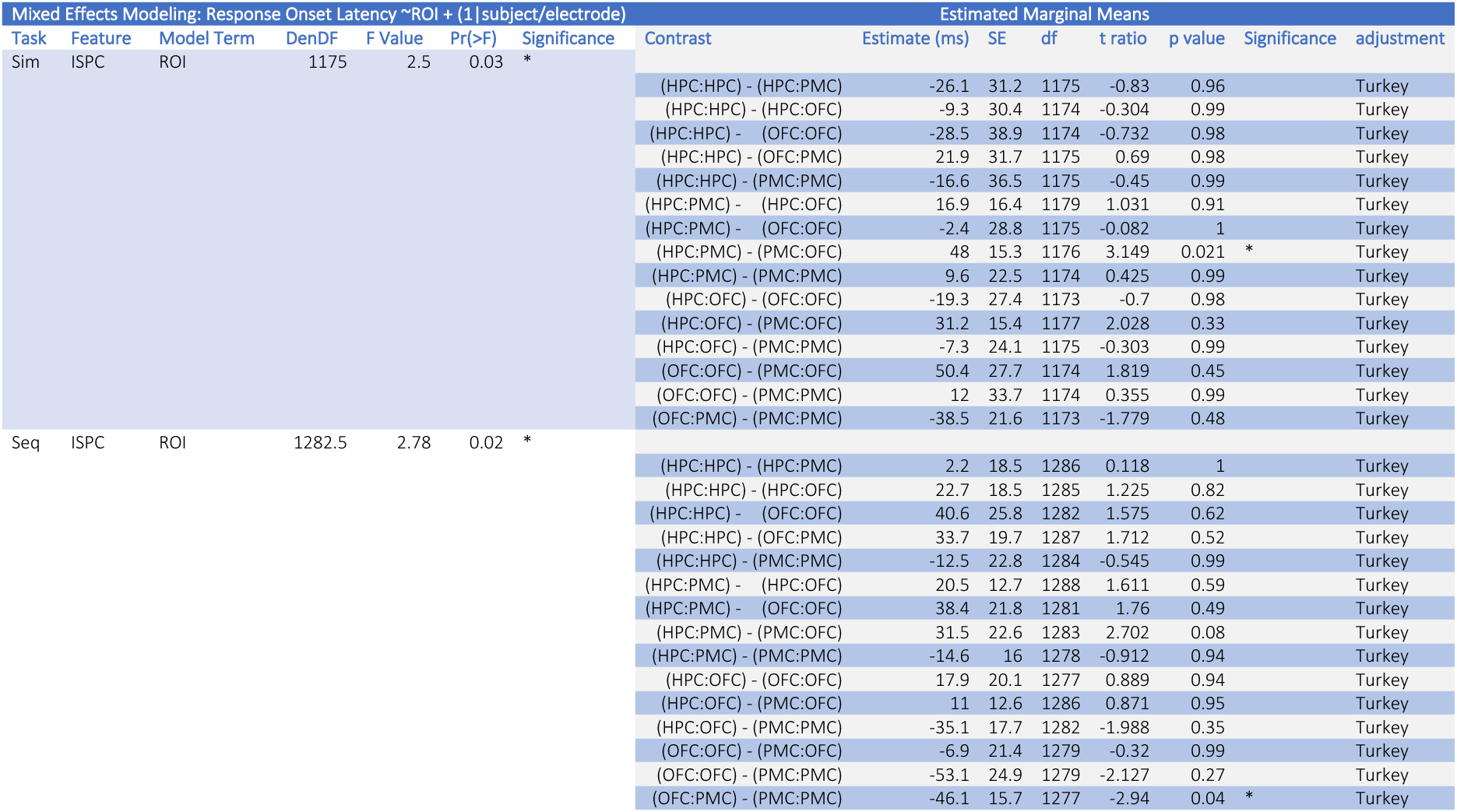
Mixed-effects modeling of ISPC in the first cohort. This table displays the ROL effects across different ISPC coupling sites. While significant differences were observed for a few ISPC comparisons, ISPC was assumed to have the same timing across connection types since the relative magnitude of this effect was much weaker than the effect observed in the power comparisons.

**Table S9:** Significant activation of single pulse electrical stimulation (STIM) This table displays the connections showing significant activation following STIM for different ROI types. Average connections per subject represent the total number of electrode pairs that could be active for a given subject. Activation percent represents the percentage of these connections within individual subjects that displayed significant causal effective connectivity. Values represent averages and standard errors calculated across subjects.

| ROI | Num Subs | Avg. Connect/Sub (± SE) | Activation Percent (%) |
| --- | --- | --- | --- |
| HPC->ANT | 8 | 19.75 ± 6.27 | 74.17 ± 6.50 |
| OFC->ANT | 7 | 29.14 ± 9.05 | 58.29 ± 12.20 |
| ANT->OFC | 7 | 40.29 ± 12.41 | 56.82 ± 10.18 |
| HPC->PMC | 8 | 47.38 ± 13.47 | 51.24 ± 10.20 |
| ANT->PMC | 9 | 25.55 ± 5.60 | 41.30 ± 10.63 |
| HPC->OFC | 6 | 45.5 ± 12.04 | 36.22 ± 7.10 |
| PMC->ANT | 9 | 18.89 ± 3.12 | 33.47 ± 6.24 |
| PMC->HPC | 9 | 47.44 ± 9.80 | 27.26 ± 7.88 |
| ANT->HPC | 9 | 26.11 ± 8.62 | 19.92 ± 5.60 |
| OFC->HPC | 7 | 57.0 ± 15.75 | 18.81 ± 7.15 |
| PMC->OFC | 7 | 87.57 ± 24.51 | 14.88 ± 3.63 |
| OFC->PMC | 7 | 77.29 ± 17.91 | 4.13 ± 1.48 |

**Table S10:**
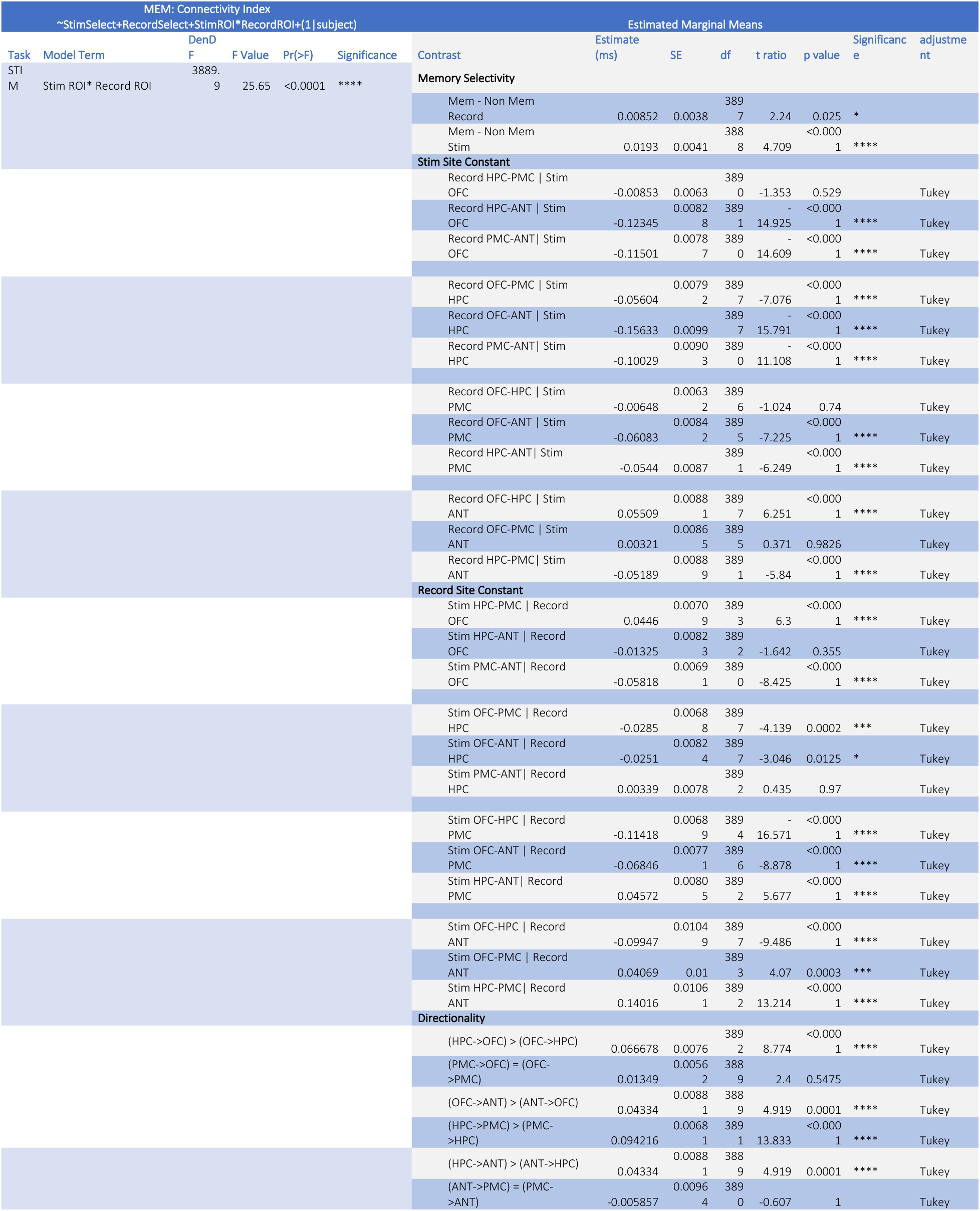
Connectivity index derived from ITPC. This table displays the significant interactions observed between stimulation and recording ROI when modeling the connectivity index following single pulse electrical stimulation. A significant interaction was observed suggesting the strength of coherence following electrical stimulation differs between stimulation and recording ROIs. Memory selectivity (Mem vs. Non Mem) is dependent on HF power during the memory task and outlined in the text.

**Table S11:**
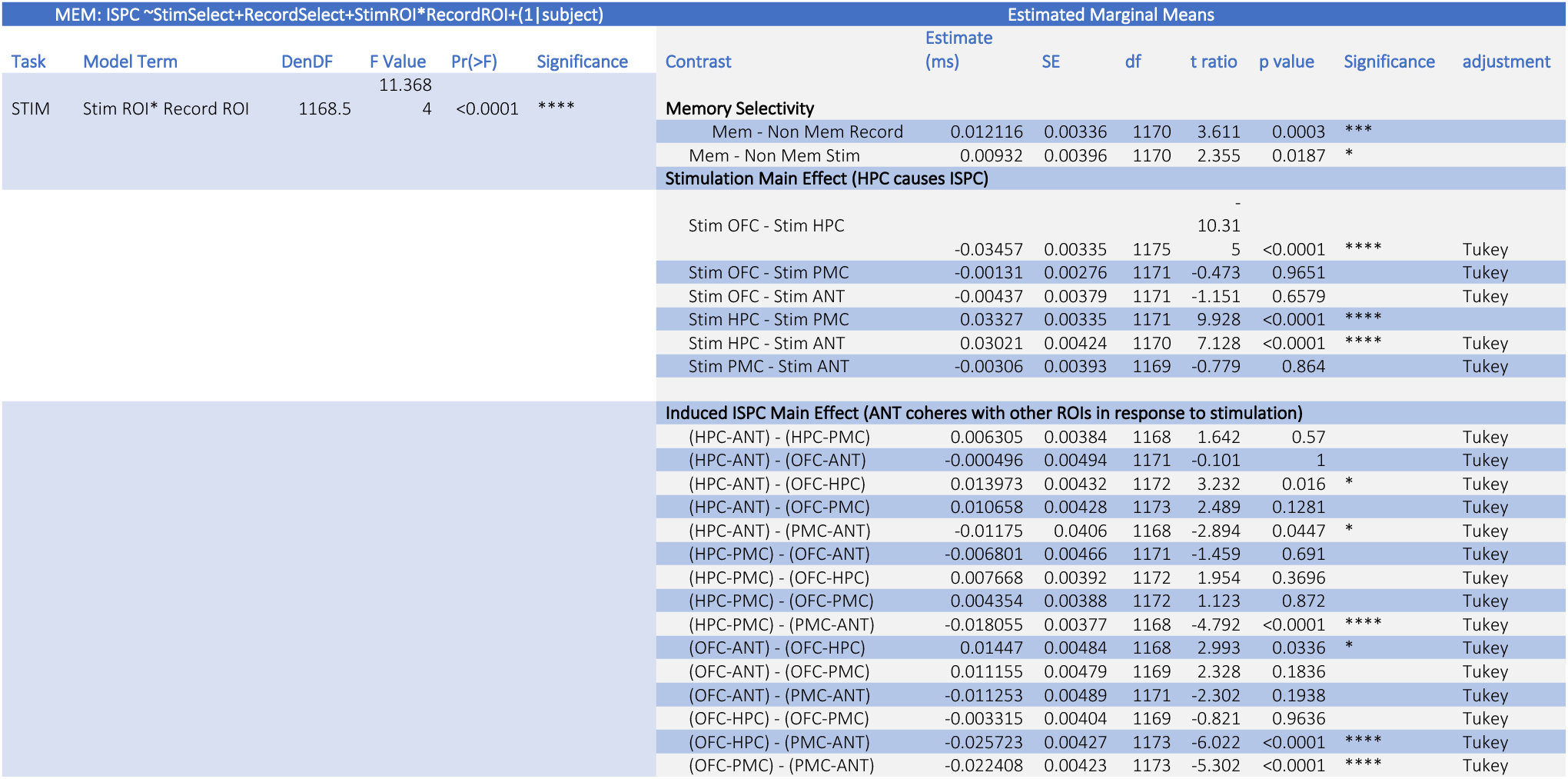
Stimulation induced ISPC. This table displays the significant interactions observed between stimulation and recording ROI when modeling the induced intersite phase coherence following single pulse electrical stimulation. A significant interaction was observed suggesting the strength of coherence following electrical stimulation differs between stimulation and recording ROIs. Memory selectivity (Mem vs. Non Mem) is dependent on HF power during the memory task and outlined in the text. Due to the large number of comparisons, marginal means were omitted from this table. However, screenshots of statistical outputs similar to those displayed in Table S10 can be provided upon request.

**Table S12:**
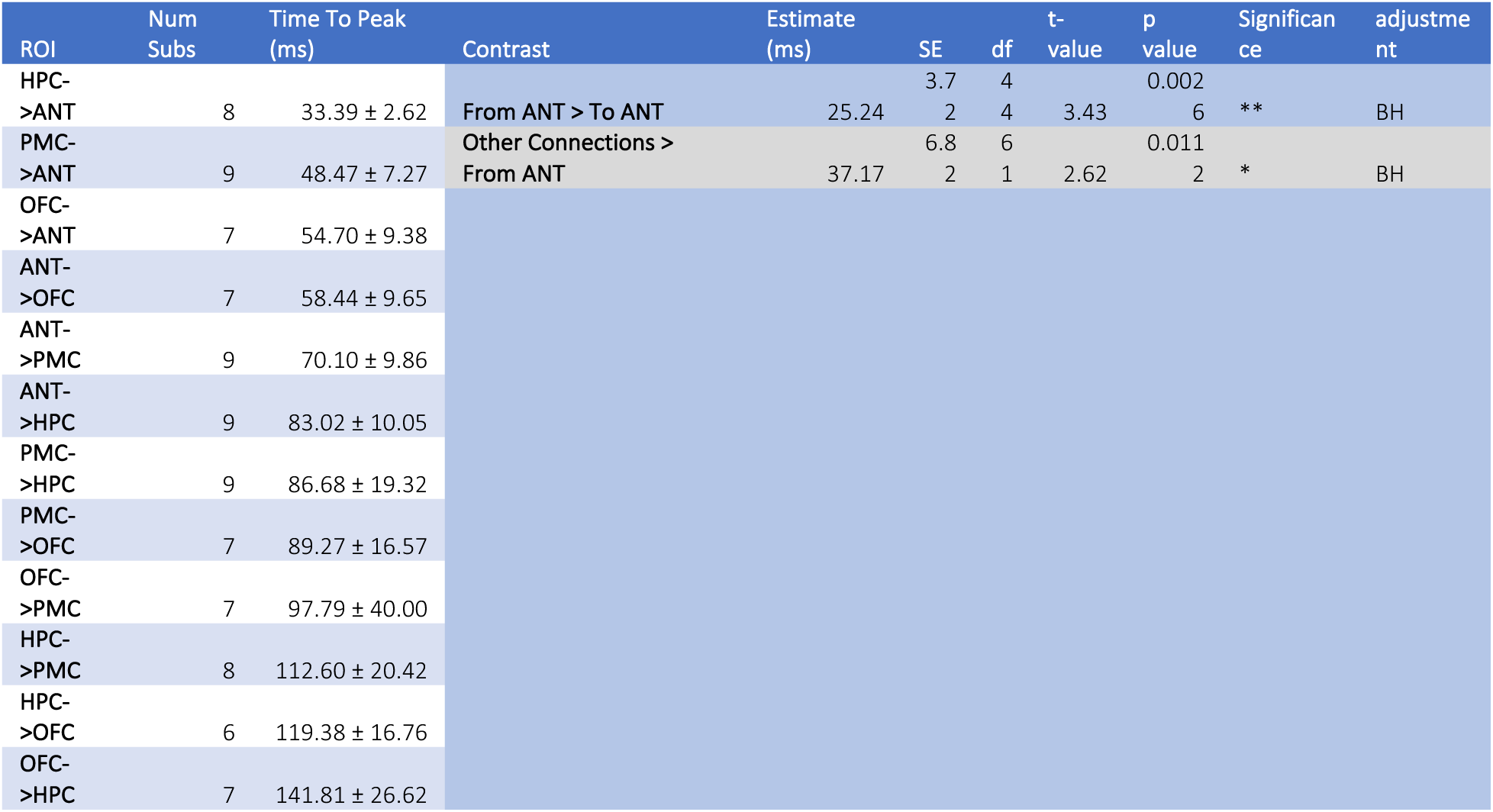
Time to first peak following single pulse electrical stimulation (STIM) When a significantly activated connection was found, the time to first peak was recorded for these connections then averaged for each subject and connection type. Values represent averages and standard errors calculated across subjects. Connections from ROIs to ANT were found to arrive significantly earlier than ANT to the other ROIs, which in turn were significantly earlier than all other connections.

